# Locally adaptive inversions modulate genetic variation at different geographic scales in a seaweed fly

**DOI:** 10.1101/2020.12.28.424584

**Authors:** Claire Mérot, Emma Berdan, Hugo Cayuela, Haig Djambazian, Anne-Laure Ferchaud, Martin Laporte, Eric Normandeau, Jiannis Ragoussis, Maren Wellenreuther, Louis Bernatchez

**Author notes:** Claire Mérot, (corresponding author).

## Abstract

Across a species range, multiple sources of environmental heterogeneity, at both small and large scales, create complex landscapes of selection, which may challenge adaptation, particularly when gene flow is high. One key to multidimensional adaptation may reside in the heterogeneity of recombination along the genome. Structural variants, like chromosomal inversions, reduce recombination, increasing linkage disequilibrium among loci at a potentially massive scale. In this study, we examined how chromosomal inversions shape genetic variation across a species range, and ask how their contribution to adaptation in the face of gene flow varies across geographic scales. We sampled the seaweed fly *Coelopa frigida* along a bioclimatic gradient stretching across 10° of latitude, a salinity gradient and a range of heterogeneous, patchy habitats. We generated a chromosome-level genome assembly to analyse 1,446 low-coverage whole genomes collected along those gradients. We found several large non-recombining genomic regions, including putative inversions. In contrast to the collinear regions, inversions and low recombining regions differentiated populations more strongly, either along an ecogeographic cline or at a fine-grained scale. These genomic regions were associated with environmental factors and adaptive phenotypes, albeit with contrasting patterns. Altogether, our results highlight the importance of recombination in shaping adaptation to environmental heterogeneity at local and large scales.

## Introduction

Across its range, a species experiences variable environmental conditions at both small and large geographic scales. This environmental heterogeneity makes local adaptation a complex process driven by multiple dimensions of selection and constrained by the distribution of genetic diversity within the genome and the intensity of gene flow acting on it (Savolainen et al. 2013; Tigano and Friesen 2016). Recombination plays a complex role in mediating this process (Stapley et al. 2017). On the one hand, recombination reduces Hill–Robertson interference, allowing natural selection to act on single loci (Otto and Barton 2001; Roze and Barton 2006). On the other hand, recombination homogenizes populations and reshuffles coadapted or locally adapted groups of alleles (Charlesworth and Charlesworth 1979; Lenormand and Otto 2000). Hence, the landscape of recombination influences adaptive trajectories, depending on the distribution of environmental heterogeneity, epistasis and gene flow (Charlesworth and Charlesworth 1979; Lenormand and Otto 2000; Yeaman 2013), and is expected to modulate the geographic distribution of adaptive and non-adaptive genetic diversity (Ortiz-Barrientos and James 2017; Stevison and McGaugh 2020).

Chromosomal inversions are major modifiers of the recombination landscape, whereby recombination between the standard and inverted arrangements is reduced in heterokaryotypes (Sturtevant 1917; Hoffmann et al. 2004). A single species can have multiple polymorphic inversions, each of them covering hundreds of kilobases or megabases, thus their impact can be widespread across the genome (Wellenreuther and Bernatchez 2018). For instance, five polymorphic inversions are present worldwide in *Drosophila melanogaster* (Kapun and Flatt 2019) and maize (*Zea mays*) harbours a 100Mb inversion (Fang et al. 2012). The last decade has shown that such inversion polymorphisms occur in a wide range of species and has brought important insights into the adaptive role of inversions (Hoffmann and Rieseberg 2008; Wellenreuther and Bernatchez 2018; Mérot, Oomen, et al. 2020). Inversions with a large effect on complex multitrait phenotypes, such as life-history, behaviour, and colour patterns, confirm that arrangements can behave as haplotypes of a “supergene”, linking together combinations of alleles within each arrangement (Joron et al. 2011; Schwander et al. 2014; Kirubakaran et al. 2016; Wellenreuther and Bernatchez 2018; Yan et al. 2020). Inversions are also notable for their associations with segregation distorters, involving epistatic selection which favours linkage between coadapted alleles at interacting loci (Sturtevant and Dobzhansky 1936; Fuller et al. 2020). Likewise, covariation between inversion frequencies and environmental variables, whether spatial, temporal or experimental, (Dobzhansky 1948; Schaeffer 2008; Kapun et al. 2016; Kirubakaran et al. 2016; Faria et al. 2019; Kapun and Flatt 2019; Huang and Rieseberg 2020) is consistent with selection for the suppression of recombination between locally adaptive loci (Kirkpatrick and Barton 2006; Charlesworth and Barton 2018) and/or coadaptive epistatic interactions between loci (Dobzhansky and Dobzhansky 1970; Charlesworth and Charlesworth 1973). Hence, when investigating adaptation with respect to multiple scales and at multiple sources of environmental variation, it is important to examine the role of large inversions or any recombination suppressors.

*Coelopa frigida* is a seaweed fly that inhabits piles of rotting seaweed, so-called wrackbeds (Fig. 1), on the east coast of North America and in Europe. *C. frigida* is known to harbour one large inversion on chromosome I (hereafter called *Cf-Inv(1)*) that is polymorphic in Europe and America (Butlin, Collins, et al. 1982; Mérot et al. 2018), as well as four additional large polymorphic inversions described in one British population (Aziz 1975). The inversion *Cf-Inv(1)* encapsulates 10% of the genome and has two arrangements: α and β. These alternative *Cf-Inv(1)* arrangements have opposing effects on body size, fertility and development time, a combination of traits which results in different fitnesses depending on the local characteristics of the wrackbed (Butlin, Read, et al. 1982; Day et al. 1983; Butlin and Day 1985; Edward and Gilburn 2013; Wellenreuther et al. 2017; Berdan et al. 2018; Mérot, Llaurens, et al. 2020). Almost nothing is known about the other inversions but, given that a large fraction of the *C. frigida* genome is affected by polymorphic inversions, one can expect that these inversions play a significant role in structuring genetic variation and contribute to local adaptation. Spatial genetic structure and connectivity in *C. frigida* remain poorly described, although occasional long distance migration bursts have been documented and regular dispersal is expected between nearby subpopulations occupying discrete patches of wrackbed (Egglishaw 1960; Dobson 1974). *C. frigida* occupies a wide climatic range of temperature (temperate to subarctic zones) as well as salinity (from freshwater to fully saline sites). Furthermore, *C. frigida* experiences high variability in the quality and the composition of its wrackbed habitat (Egglishaw 1960; Dobson 1974). These sources of habitat heterogeneity vary at both large and local geographic scales, for which, depending on the scale of dispersal, a linked genomic architecture may be favourable.

**Figure 1:**
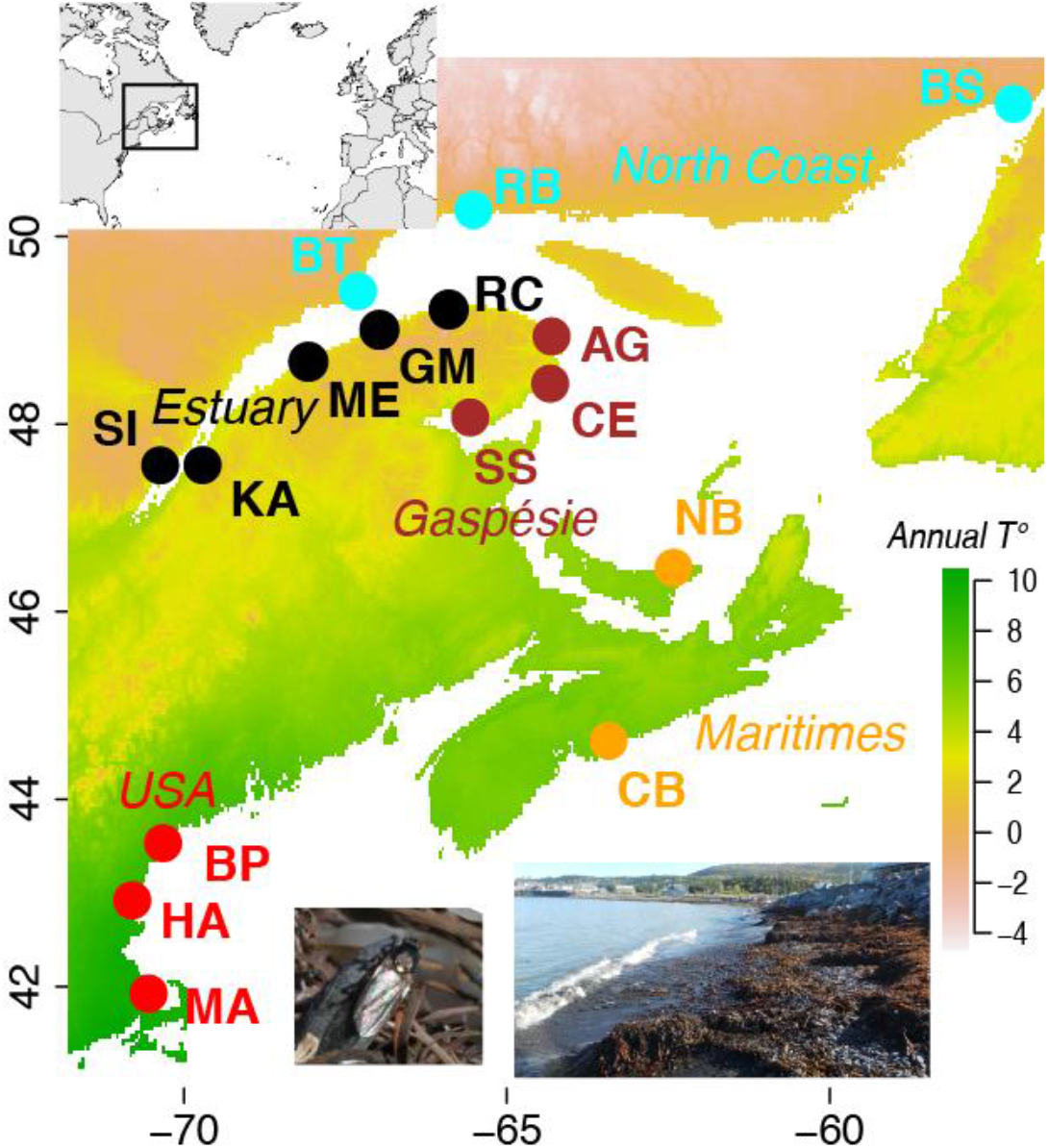
*Coelopa frigida* sampling across an environmental gradient. Map of the 16 sampling sites, coloured by geographic region. The background of the map displays the gradient of annual mean air temperature. The insert shows the location of the study area at a wider scale. Photos show *Coelopa frigida* and its habitat of seaweed beds.

In the present study, we investigated how chromosomal inversions contribute to local adaptation across different scales of environmental heterogeneity, and how they shape genetic diversity. Using the seaweed fly *Coelopa frigida* as a biological model, we adopted a systematic approach for localizing multiple chromosomal inversions, and analysed genetic variation across several dimensions of environmental variation including a 1,500 km climatic gradient, a salinity gradient, and fine scale, patchy habitat variation (Fig. 1). We built the first reference genome assembly for *C. frigida* and sequenced 1,446 whole genomes at low coverage. Using this comprehensive data set, we analysed patterns of genetic polymorphism along the genome to identify putative inversions. As connectivity between populations of *C. frigida* was previously unknown, we examined its geographic structure with respect to SNP markers. Finally, we tested genotype-environment and genotype-phenotype associations to determine the genomic architecture of adaptation to various sources of environmental variation acting at different geographic scales.

## Results

To facilitate our analyses, we built the first reference genome assembly for *Coelopa frigida* using a combination of long-read sequencing (PacBio) and linked-reads from 10xGenomics technology. A high-density linkage map (28,639 markers segregating across 6 linkage groups) allowed us to anchor and orientate more than 81% of the genome into 5 large chromosomes (LG1 to LG5) and one small sex chromosome (LG6). This karyotype was consistent with previous cytogenetic work on *C. frigida* (Aziz 1975) and with the 6 Muller elements (*A*=*LG4, B*=*LG5, C*=*LG2, D*=*LG3, E*=*LG1, F*=*LG6*, Fig. S1), which are usually conserved in Diptera (Vicoso and Bachtrog 2015; Schaeffer 2018). The final assembly included 6 chromosomes and 1,832 unanchored scaffolds with a N50 of 37.7 Mb for a total genome size of 239.7 Mb. This reference had a high level of completeness, with 96% (metazoan) and 92% (arthropods) of universal single-copy orthologous genes completely assembled. It was annotated with a highly complete transcriptome (87% complete BUSCOs in the arthropods) based on RNA-sequencing of several ontogenetic stages and including 35,999 transcripts.

To analyse genomic variation at the population-scale, we used low coverage (~1.4X) whole genome sequencing of 1,446 flies from 16 locations along the North American Atlantic coast (88-94 adult flies/location). Sampled locations spanned a North-South gradient of 1,500km, over 10° of latitude, a pronounced salinity gradient in the St Lawrence Estuary, and a range of habitats with variable seaweed composition and wrackbed characteristics (Fig. 1, Table S1). After alignment of the 1,446 sequenced individuals to the reference genome, we analysed genetic variation within a probabilistic framework accounting for low coverage (ANGSD, Korneliussen et al. 2014) and reported 2.83 million single nucleotide polymorphisms (SNPs) with minor allelic frequencies higher than 5% for differentiation analyses.

### • Two large chromosomal inversions structure intraspecific genetic variation

Decomposing SNPs genotype likelihoods through a principal component analysis (PCA) revealed that the 1^st^ and 2^nd^ principal components (PCs) contained a large fraction of genetic variance, respectively 21.6 % and 3.9 %, and allowed us to display the 1,446 flies as 9 discrete groups (Fig 2A). Along PC1, the three groups corresponded to three genotypes of the inversion *Cf-Inv(1)* (αα, αβ, ββ), as identified using two diagnostic SNPs (Mérot et al. 2018) with respectively 100% and 98.3% concordance (Table S2). Along PC2, three distinct groups were identified that corresponded neither to sex nor geographic origins, and thus possibly represented three genotypes for another polymorphic inversion.

**Figure 2:**
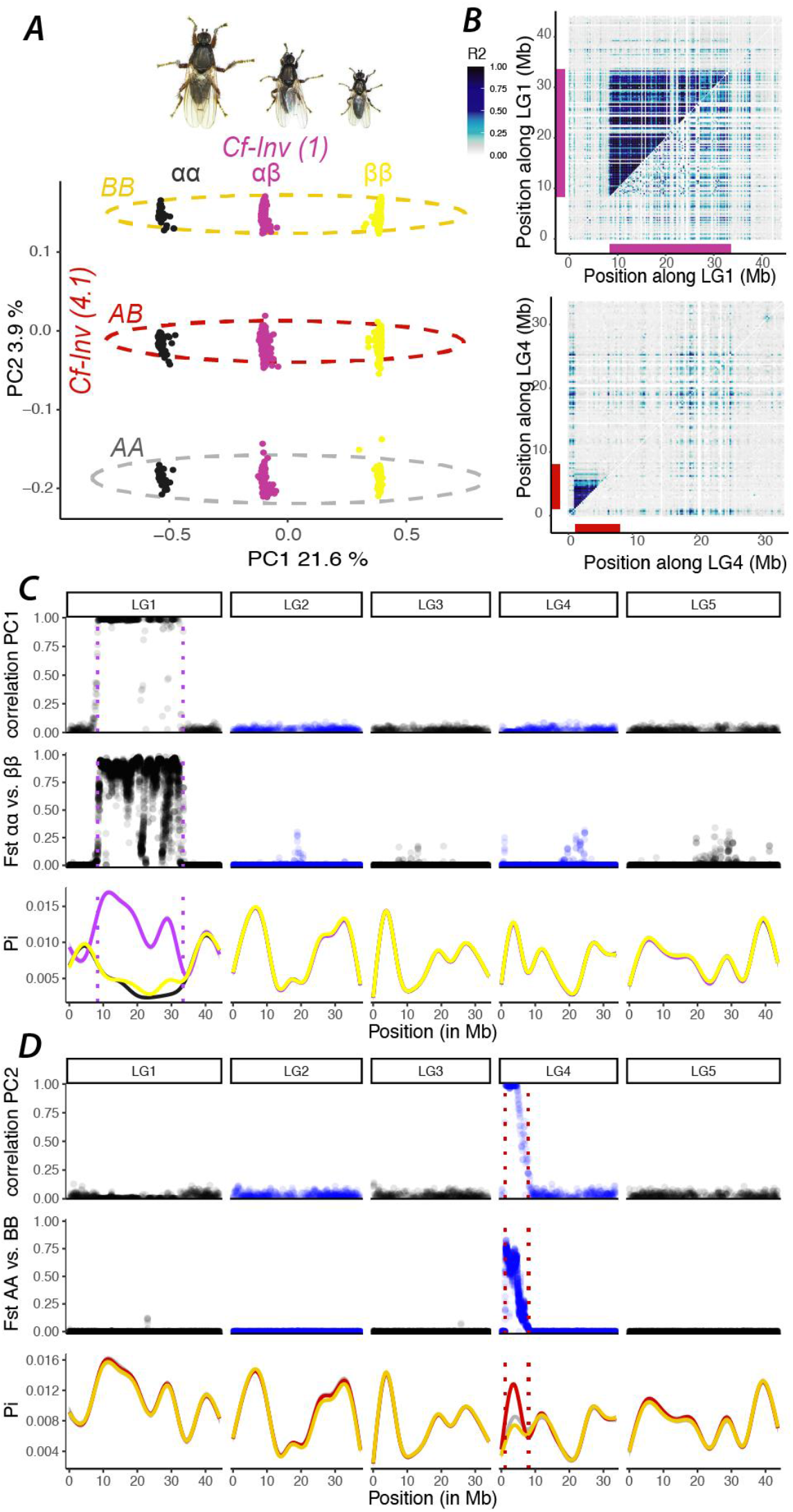
Two large chromosomal inversions structure within-species genetic variability. **(A)** Principal component analysis (PCA) of whole-genome variation. Individuals are coloured by karyotypes at the inversion *Cf-Inv(1)*, as determined previously with a SNP marker (Mérot et al. 2018). Ellipses indicate secondary grouping along PC2. **(B)** Linkage disequilibrium (LD) in LG1 and LG4. The upper triangles include all individuals and the lower triangles include homokaryotes for the most common arrangement for each inversion. Bars represent the position of the inversions. The colour scale shows the 2^nd^ higher percentile of the R^2^ value between SNPs summarized by windows of 250kb **(C)** Along the genome, correlation between PC1 scores of local PCAs performed on windows of 100 SNPs and PC1 scores of the PCA performed on the whole genome; F_ST_ differentiation between the two homokaryotypes of *Cf-Inv(1)* in sliding-windows of 25kb; and nucleotide diversity (π) within the three karyotypic groups of *Cf-Inv(1)* smoothed for visualization. Dashed lines represent the inferred boundaries of the inversion *Cf-Inv(1)* **(D)** Correlation between PC1 scores of local PCAs performed on windows of 100 SNPs and PC2 scores of the PCA performed on the whole genome; FST differentiation between the two homokaryotypes of *Cf-Inv(4.1)* in sliding-windows of 25kb; and nucleotide diversity (π) within the three karyotypic groups of *Cf-Inv(4.1)* smoothed for visualization. Dashed lines represent the inferred boundaries of the inversion *Cf-Inv(4.1)*.

To assess which regions of the genome reflected the patterns observed in the whole genome PCA, we performed local PCA on windows of 100 SNPs along each chromosome and evaluated the correlation between PC1 scores of each local PCA and PCs scores of the global PCA (Fig. 2C). PC1 was highly correlated with a region of 25.1 Mb on LG 1, indicating the genomic position of the large *Cf-Inv(1)* inversion (Table 1). PC2 was highly correlated with a smaller region of 6.9 Mb on LG4 (Fig. 2D), consistent with the hypothesis of an inversion, hereafter called *Cf-Inv(4.1)*. Several other characteristics were consistent with the hypothesis that these two regions are inversions. First, inside these regions, linkage disequilibrium (LD) was very high when considering all individuals, but low within each group of homokaryotypes (Fig 2B). This indicates that recombination is limited between the arrangements but occurs freely in homokaryotypes bearing the same arrangement. Second, F_ST_ was very high between homokaryotes in the inverted region (*Cf-Inv(1)* αα vs. ββ: 0.75, *Cf-Inv(4.1)* AA vs. BB: 0.51, Fig. 2C) compared to low values in the rest of the genome (*Cf-Inv(1)* αα vs. ββ: 0.002, *Cf-Inv(4.1)* AA vs. BB: 0.001, Fig. 2D). Third, the intermediate group on the PCA was characterized by a higher proportion of observed heterozygotes for SNPs in the inverted region than the extreme groups, confirming that this is probably the heterokaryotypic group (Fig. S2).

**Table 1:** Name, position and characteristics of the putative inversions and regions appearing as cluster of outlier windows in the local PCA analysis. For putative inversions, absolute nucleotide divergence (d_XY_) in non-coding regions was calculated between homokaryotypic groups, and corrected by the mean of nucleotide diversity (π) within homokaryotypic groups by windows of 25kb. Numbers between square brackets indicate confidence intervals drawn by bootstrapping windows of 25kb.

| Name | Status | Chr | start | stop | size<br>(MB) | $d_{xy}$ |
| --- | --- | --- | --- | --- | --- | --- |
| *Cf-Inv(1)* | *Known inversion* | LG1 | 8342182 | 33487673 | 25.1 | 1.84% [1.80-1.88] |
| *Cf-Inv(4.1)* | *Probable inversion* | LG4 | 1088816 | 7995568 | 6.9 | 0.64% [0.61-0.67] |
| *Cf-Inv(4.2)* | *Probably two linked* | LG4 | 22421881 | 25145365 | 2.7 | 0.079% [0.061-0.096] |
| *Cf-Inv(4.3)* | *inversions* | LG4 | 30622035 | 31991919 | 1.4 | 0.32% [0.31-0.34] |
| *Cf-Lrr(2)* | *Low-recombination region* | LG2 | 14083320 | 20869940 | 6.8 |  |
| *Cf-Lrr(3)* | *Low-recombination region* | LG3 | 7486933 | 13829649 | 6.3 |  |
| *Cf-Lrr(5)* | *Low-recombination region* | LG5 | 15940464 | 32665323 | 16.7 |  |

Nucleotide diversity, as measured by π, was similar between karyotypic groups along the genome, and higher in the heterokaryotypes than in the homokaryotypes in inverted regions (Fig. 2C-D). For both inversions, nucleotide diversity was comparable between homokaryotes. Absolute nucleotide divergence between arrangements was strong in inverted regions (Table 1, Fig. S3). Assuming a mutation rate comparable to *Drosophila* (5 ×10^−9^ mutations per base per generation (Assaf et al. 2017)), and approximately 5 to 10 generations per year, we thus estimated, from absolute divergence at non-coding regions that the arrangements split at least 180,000 to 376,000 years ago for *Cf-Inv(1)* and at least 61,000 to 134,000 years ago for *Cf-Inv(4.1)*.

### • *C. frigida* exhibit other regions including non-recombining haplotypic blocks

To further examine the heterogeneity of genetic structure along the genome, we reanalysed the local PCAs using a method based on multidimensional scaling (MDS) which identifies clusters of PCA windows displaying a common pattern. This method has been previously used to identify and locate non-recombining haplotypic blocks (Li and Ralph 2019; Huang et al. 2020; Todesco et al. 2020). Besides the aforementioned *Cf-Inv(1)* and *Cf-Inv(4.1)* inversions, which caused the 1^st^ and 2^nd^ axis of the MDS, we identified five outlier genomic regions across the different MDS axes (Fig.3, Fig. S4). In all five regions, a large proportion of variance was captured along the 1^st^ PC (>50%), and linkage disequilibrium was high (Fig. 3A).

**Figure 3:**
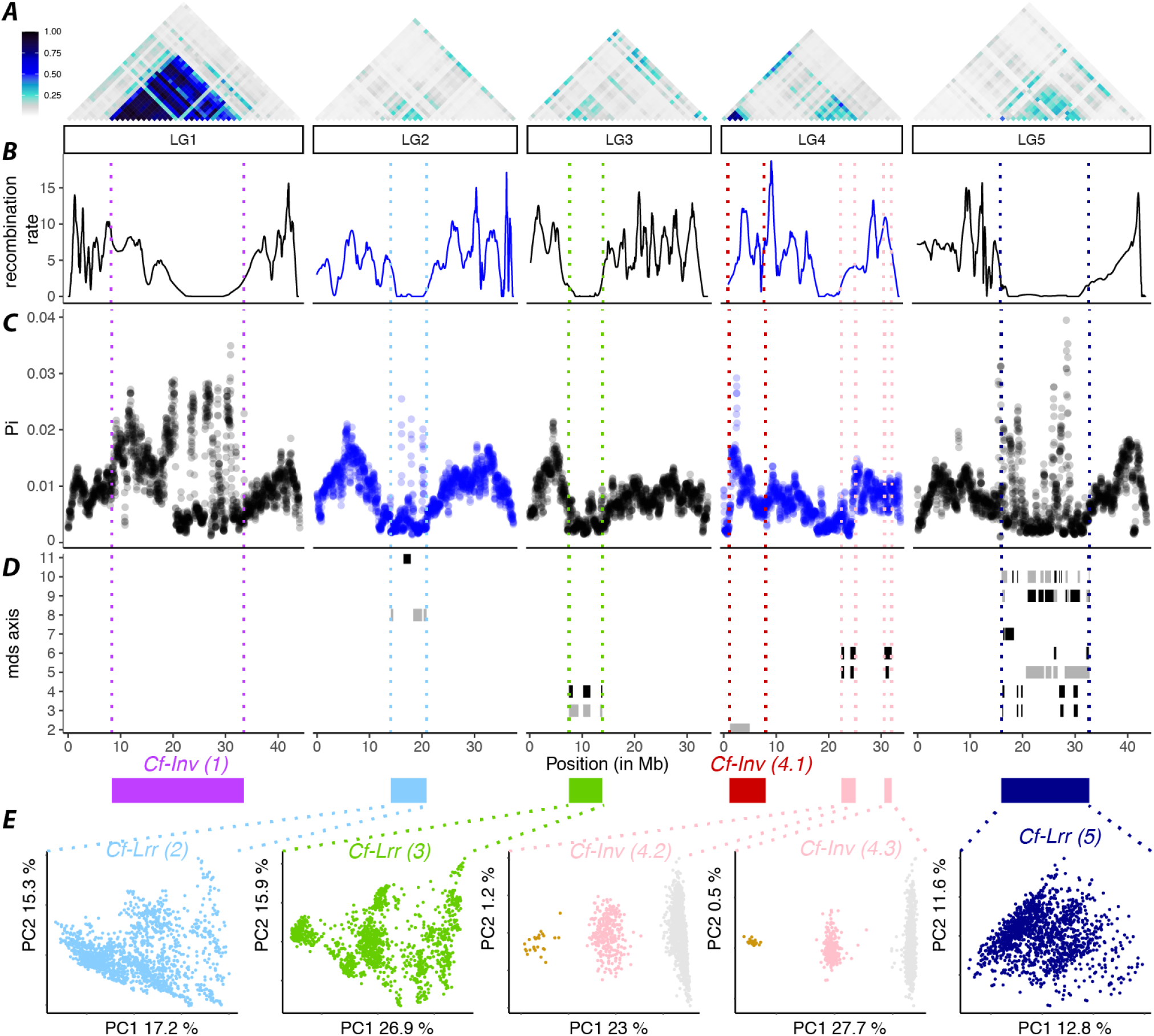
Detecting other regions exhibiting non-recombining haplotypic blocks. **(A)** LD across the 5 major chromosomes expressed as the 2^nd^ higher percentile of the R^2^ value between SNPs summarized by windows of 1Mb. **(B)** Recombination rate (in cM/Mb) inferred from the linkage map, smoothened with a loess function accounting for 10% of the markers. **(C)** Nucleotide diversity (π) by sliding-windows of 100kb (step 20kb) averaged across the different geographic populations. **(D)** Position along the genome of clusters of local PCA windows scored as outliers (>4sd) along each axis of the MDS, at the upper end in black, and the lower end in grey. Coloured rectangles indicate the position of the inversions and the regions of interest gathering outlier clusters or putative inversions. Dashed lines represent their inferred boundaries across all plots. **(E)** PCA performed on SNPs within each region of interest. For the two regions on LG4 that appear as two linked putative inversions (*Cf-Inv(4.2)* and *Cf-Inv(4.3)*), three clusters were identified with high-confidence and coloured as putative homokaryotes and heterokaryotes. The same colours are used in both regions since karyotyping was consistent across all individuals.

Two regions on LG4 represented convincing putative inversions of 2.7Mb and 1.4Mb, respectively. In both regions, the PCA displayed three groups of individuals with high clustering confidence; the central group contained a high proportion of heterozygotes and the extreme groups were differentiated (Fig. 3E, Fig. S5). Within these two regions, nucleotide diversity was comparable between haplogroups groups and the absolute divergence (d_XY_) between homokaryotypes was lower than for *Cf-Inv(1)* and *Cf-Inv(4.1)*, suggesting younger inversions which could have diverged as recently as 6,000 to 68,000 years ago. Karyotype assignment was the same between the two putative inversions, indicating that they are either tightly linked or belong to a single inversion. Two lines of evidence support the hypothesis that these are two inversions. First, the high density of linkage map markers and the non-null recombination rate across this area of 50 cM provided confidence in the genome assembly and supported a gap of 5 Mb between the two inversions. Moreover, previous cytogenetic work showed that one chromosome of *C. frigida* exhibits a polymorphic inversion on one arm (possibly *Cf-Inv(4.1)*) and, on the other arm, two polymorphic inversions which rarely recombine (Aziz 1975). Both inversions were subsequently analysed together and called *Cf-Inv(4.2)* and *Cf-Inv(4.3)*.

The other three regions, spanning 6.8 Mb on LG2, 6.3 Mb on LG3 and 16.7 Mb on LG5, represented complex areas that behaved differently from the rest of the genome. Recombination was locally reduced, both in the linkage map and in wild populations, as indicated by strong linkage disequilibrium (Fig. 3A-B). These three regions were all highly heterogeneous; within each region, nucleotide diversity showed highly contrasting pattern across subregions (Fig. 3C). A fraction of these subregions exhibited low nucleotide diversity, which may correspond to centromeric or pericentromeric regions (Fig. 3C, Fig. S6), as well as a high density of transposable elements, such as LINEs or LTRs (Fig S7). However, these low diversity subregions were interspersed with subregions of high diversity, particularly on LG5 (Fig 3C). Some of those high diversity subregions also corresponded to clusters of outlier windows in the local PCA analysis and appeared as non-recombining haplotypic blocks of medium size (1Mb-2Mb) in partial LD (Fig. S8-S10). In the absence of more information about the mechanisms behind the reduction in recombination, we consider those three regions of the genome to be simply “low recombining regions” (subsequently called *Cf-Lrr(2), Cf-Lrr(3), Cf-Lrr(5)*). Accordingly, the fraction of the genome subsequently called “collinear” excluded both these regions and the inversions (*Cf-Inv(1), Cf-Inv(4.1), Cf-Inv(4.2)*, and *Cf-Inv(4.3)*).

### • Geographic structure shows distinctive signals in inverted and low recombining regions

Geography also played a major role in structuring genetic variation. Our 3^rd^ PC, which explained 1.4% of variance, represented genetic variation along the North-South gradient (Fig. 4A). Differentiation between pairs of populations, measured as F_ST_ on a subset of LD-pruned SNPs, also followed the North-South gradient but was globally weak (F_ST_ = 0.003 to 0.016, Fig. S11)). We also detected a strong signal of isolation by distance (IBD) when examining the correlation between genetic distances and Euclidean distances among the 16 populations (R^2^=0.45, F=97, p<0.001, Table S3). Considering least cost distances along the shorelines instead of Euclidian distances between locations improved the model fit (R^2^=0.63, F=199, p<0.001, ΔAIC=47, Table 2, Table S3). This supports a pattern of isolation by resistance (IBR, see Methods), in which dispersal occurs primarily along the coastline and is limited across the mainland or the sea.

**Figure 4:**
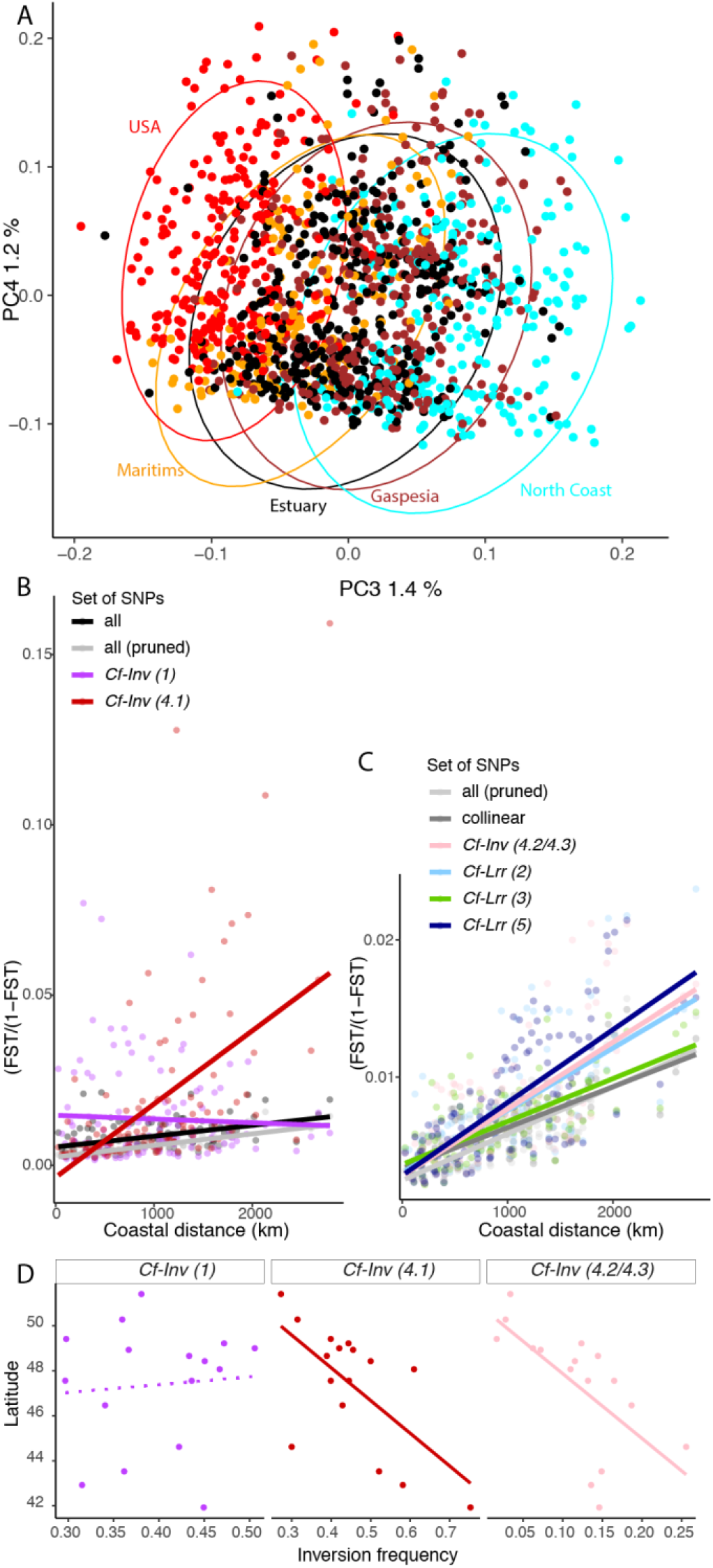
Genetic variation is geographically structured along a North-South gradient and displays isolation-by-resistance. **(A)** 3^rd^ and 4^th^ principal components of a PCA on whole-genome variation. Individuals are coloured by their geographic region, as in Fig. 1 **(B-C)** Isolation by resistance displayed as the association between genetic distance (F_ST_/(1-F_ST_) and the distance by the least-cost path following the coast. Colours denote the subset of SNPs used for the calculation of the F_ST_. The results are displayed in two panels with different y scales to better display the lower values. **(D)** Latitudinal variation of inversion frequencies.

**Table 2:** Association between genetic distance and geographic distances measured as least-cost distances along the shoreline (Isolation-by-resistance) for the different fractions of the genome. Numbers between square brackets indicate the limits of the 95% distribution of the slope coefficient. The comparison to collinear regions displays the output of a full model comparing each region to the collinear genome, providing the direction and the significance (*) of the interaction term.

| SNP subset | R<sup>2</sup><br>adjusted | F | p-<br>value | intercept | slope coefficient | Comparison<br>to collinear<br>regions |
| --- | --- | --- | --- | --- | --- | --- |
| All | 0.19 | 29.3 | <0.001 | 0.0085 | 0.0020 [0.0013-0.0027] |  |
| Collinear | 0.54 | 138.6 | <0.001 | 0.0062 | 0.0019 [0.0015-0.0022] |  |
| LD pruned | 0.63 | 199.5 | <0.001 | 0.0057 | 0.0021 [0.0018-0.0024] |  |
| *Cf-Inv(1)* | -0.01 | 0.3 | 0.59 | 0.0137 | -0.0006 [-0.0032-0.0018] | - * |
| *Cf-Inv(4.1)* | 0.29 | 49.4 | <0.001 | 0.0172 | 0.0134 [0.0096-0.0172] | + * |
| *Cf-Inv(4.2/4.3)* | 0.50 | 121.5 | <0.001 | 0.0075 | 0.0030 [0.0025-0.0036] | + * |
| *Cf-Lrr(2)* | 0.44 | 95.4 | <0.001 | 0.0074 | 0.0028 [0.0023-0.0034] | + * |
| *Cf-Lrr(3)* | 0.49 | 113.1 | <0.001 | 0.0066 | 0.0019 [0.0016-0.0023] | n.s. |
| *Cf-Lrr(5)* | 0.55 | 147.2 | <0.001 | 0.0080 | 0.0033 [0.0028-0.0038] | + * |

These IBD and IBR patterns varied significantly along the genome. When considering all SNPs, pairwise differentiation was more heterogeneous (F_ST_=0.002 to 0.021, Fig. 4B) and IBR was much weaker, albeit significant (R^2^=0.19, F=29, p<0.001) than when considering LD-pruned SNPs or collinear SNPs. We thus calculated pairwise F_ST_ between pairs of populations based on different subsets of SNPs, either from each inversion, from each low-recombing region, or from the collinear genome.

All of the inversions exhibited increased differentiation between populations in comparison with the collinear genome (Table S3, Fig. S12). However, the global geographic patterns differed between inversions. In the inverted region *Cf-Inv(1)*, there was no association between genetic and geographic distances (Fig. 4B, Table 2), a result which significantly contrasts with the collinear genome (Fig. S13). This result was due to highly variable pairwise genetic differentiation between populations in the inverted region *Cf-Inv(1)*. Conversely, genetic differentiation between geographic populations in the inverted regions of LG4 showed significant IBD/IBR patterns with a significantly steeper slope of regression between genetic and geographic distances compared to collinear regions (Fig. 4B-C, Table 2, Table S4, Fig. S13). The divergence between northern and southern populations was mirrored by a sharp and significant latitudinal cline of inversion frequencies, ranging from 0.27 to 0.75 for *Cf-Inv(4.1)* (GLM: z=-8.1, p<0.001, R^2^=0.41) and from 0.02 to 0.26 for *Cf-Inv(4.2/4.3)* (GLM: z=-6.6, p<0.001, R^2^=0.37). The association between latitude and inversion frequency was significantly stronger than for randomly chosen SNPs with similar average frequencies (Fig 4D, Fig. S14-S15).

Although the entire genome (with the exception of inversion *Cf-Inv(1)*) showed IBD and IBR, it was significantly increased in two of the three low recombining regions compared to the collinear regions. When compared to collinear regions of the same size, the slope of the regression between genetic and geographic distances was significantly steeper for *Cf-Lrr(2)* and *Cf-Lrr(5)* but not for *Cf-Lrr(3)* (Fig. 4C, Table 2, Table S4, Fig. S13). Overall, the geographic differentiation in the four inverted regions and two low recombining regions showed patterns differing from the collinear genome, indicating the influence of processes other than the migration-drift balance, possibly at different geographic scales for *Cf-Inv(1)* vs. others.

### • Adaptive diversity colocalizes with inversions and low recombining regions

To investigate putative patterns of adaptive variation in *C. frigida*, we analyzed the association between SNP frequencies and environmental variables at large (thermal latitudinal gradient and salinity gradient in the St. Lawrence R. Estuary) and local (abiotic and biotic characteristics of the wrackbed habitat) spatial scales (Fig. 1, Fig. S16, Table S1). Analyses with two different genotype-environment association methods (latent factor mixed models and Bayesian models) showed consistent results, highlighting high peaks of environmental associations and large clusters of outlier SNPs in the inverted or low recombining regions (Fig. 5A-E, Table 3, Table S5, Fig. S17-18). However, different inversions were implicated depending on environmental factor and spatial scale. We considered SNPs consistently identified as outliers across both analyses to be putatively adaptive.

**Figure 5:**
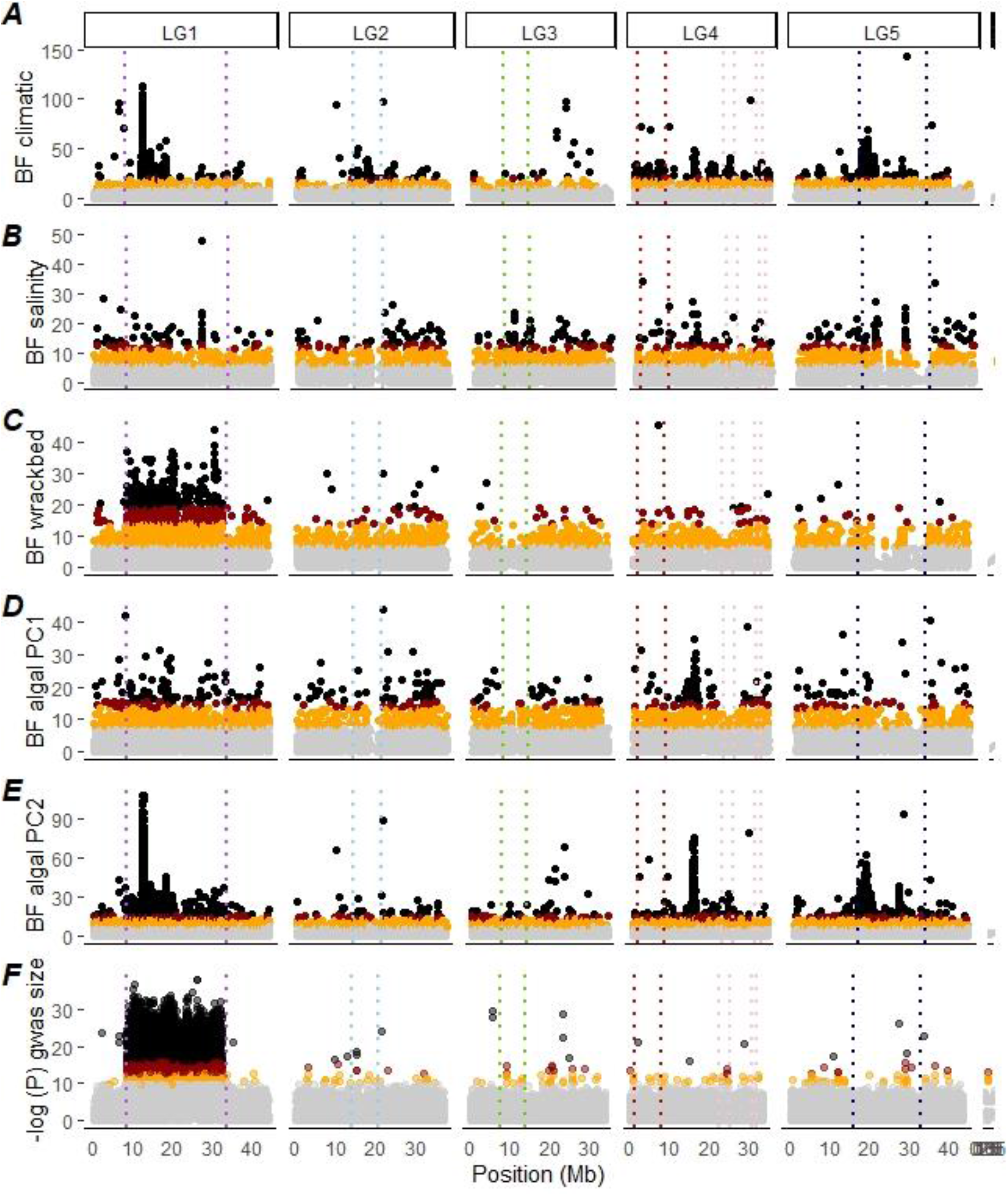
Environmental and phenotypic associations. Candidate SNPs associated with **(A)** climatic variation along the North-South gradient, **(B)** salinity variation along the Estuarian gradient, **(C)** variations in abiotic characteristics of the wrackbed habitat, **(D-E)** variation in wrackbed algal composition. The Manhattan plot shows the Bayesian factor from the environmental association analysis performed in Baypass, controlling for population structure. **(F)** Candidate SNPs associated with wing size. The Manhattan plot shows the p-values from the GWAS. Points are coloured according to false-discovery rate (black: <0.00001, red: <0.0001, orange: <0.001). Dashed lines represent the inferred boundaries of inversions and low-recombing regions.

**Table 3:** Genomic repartition of candidate SNPs associated with environmental variables. Repartition of the candidate SNPs associated with each environmental variation using the combination of two GEA methods. N is the number of outliers SNPs within a given region, % is the proportion of the outliers found in this region and OR indicate the odd-ratio. Values in bold with a star indicate significant excess of candidate SNPs in a Fisher exact test. Results obtained for each GEA method are presented in Table S5.

|  | Tested SNPs |  | Climate |  |  | Salinity |  |  | Bed abiotic characteristics |  |  | Algal composition (PC1) |  |  | Algal composition (PC2) |  |  |
| --- | --- | --- | --- | --- | --- | --- | --- | --- | --- | --- | --- | --- | --- | --- | --- | --- | --- |
|  | N | % | N | % | OR | N | % | OR | N | % | OR | N | % | OR | N | % | OR |
| All | 1155978 |  | 3635 |  |  | 509 |  |  | 780 |  |  | 372 |  |  | 2740 |  |  |
| Collinear | 814279 | 70% | 556 | 15% | 0.2 | 301 | 59% | 0.8 | 163 | 21% | 0.3 | 254 | 68% | 1.0 | 390 | 14% | 0.2 |
| *Cf-Inv(1)* | 176963 | 15% | 1474 | 41% | **2.6\*** | 64 | 13% | 0.8 | 584 | 75% | **4.9\*** | 77 | 21% | **1.4\*** | 1494 | 55% | **3.6\*** |
| *Cf-Inv(4.1)* | 57323 | 5.0% | 480 | 13% | **2.7\*** | 15 | 2.9% | 0.6 | 11 | 1.4% | 0.3 | 14 | 3.8% | 0.8 | 33 | 1.2% | 0.2 |
| *Cf-Inv(4.2/4.3)* | 17019 | 1.5% | 111 | 3.1% | **2.1\*** | 8 | 1.6% | 1.1 | 8 | 1.0% | 0.7 | 3 | 0.8% | 0.5 | 26 | 0.9% | 0.6 |
| *Cf-Lrr(2)* | 20458 | 1.8% | 93 | 2.6% | **1.4\*** | 6 | 1.2% | 0.7 | 9 | 1.2% | 0.7 | 3 | 0.8% | 0.5 | 15 | 0.5% | 0.3 |
| *Cf-Lrr(3)* | 16313 | 1.4% | 11 | 0.3% | 0.2 | 28 | 5.5% | **3.9\*** | 0 | 0.0% | 0.0 | 3 | 0.8% | 0.6 | 7 | 0.3% | 0.2 |
| *Cf-Lrr(5)* | 53623 | 4.6% | 910 | 25% | **5.4\*** | 87 | 17% | **3.7\*** | 5 | 0.6% | 0.1 | 18 | 4.8% | 1.0 | 775 | 28% | **6.1\*** |

At a large geographic scale, associations with climatic variation along the latitudinal gradient showed a strong excess of outlier SNPs in the four inversions and the low recombining regions of LG2 and LG5. These regions exhibited 2 to 5 times more outliers than expected by chance (Table 3) with particularly strong peaks of environmental association (BF >50, Fig. 5A), and a signal significantly stronger than for random blocks of collinear genome of the same size (Fig. S19). However, this was not the case for *Cf-Lrr(3)*. These results were consistent whether or not the model was controlled by the geographic population structure (Fig. S17-18). Variation of frequencies of *Cf-Inv(4.1)* and *Cf-In(4.2/4.3)* were also significantly associated with climatic variation, when considered as single loci (GLM: *Cf-Inv(4.1)*: z=-7.76, p<0.001; *Cf-Inv(4.2/4.3)*: z=-6.45, p<0.001, with the model explaining 36% and 37% of the variance in inversion frequency, Fig. S20). Variation along the salinity gradient, which also spanned variation in tidal amplitude, was significantly associated with a more limited number of SNPs. A large excess of such outliers were found in *Cf-Lrr(3)* and *Cf-Lrr(5)* (Table 3), the only two regions in which the signal of association was stronger than in the collinear genome (Fig. S19).

At a finer geographic scale, outlier SNPs associated with wrackbed abiotic characteristics (depth, temperature and salinity) were strongly enriched in the inverted region *Cf-Inv(1)* with an odds ratio of 5, including outliers with very strong support (BF >20, Fig. 4C). SNP associations with wrackbed abiotic characteristics were stronger than in collinear regions in *Cf-Inv(1)*, and more marginal in *Cf-Inv(4.2/4.3)* (Fig. S19). This was mirrored by the *Cf-Inv(1)* frequency, which significantly covaried with wrackbed (GLM, z=3.5, p<0.001, R^2^=0.26, Fig. S20). Variation in algal composition of the wrackbed, driven by the relative abundance of two dominant families of seaweed, Fucaceae or Laminariaceae, was significantly associated with widespread SNPs, although the inversion *Cf-Inv(1)* was overrepresented by an odds ratio of 1.4. Variation in secondary components of the substrate were more difficult to interpret as they covaried with latitude and temperature (Fig. S16). Despite this, these secondary components were also associated with a large number of SNPs in *Cf-Inv(1)* and in *Cf-Lrr(5)* with odds ratio of 3.6 to 6 (Fig. 5E), and a distribution of association scores significantly higher than in collinear blocks (Fig. S19).

### • Genotype-phenotype association

As wrackbed composition and *Cf-Inv(1)* are known to influence adult size (Butlin, Read, et al. 1982; Edward and Gilburn 2013), we used a GWAS to uncover genetic variation associated with wing size. Among the 124,701 candidate SNPs identified by the GWAS, more than 99.8% were located in *Cf-Inv(1)* (Fig. 5F). When variation in karyotype was removed (by running the analysis with only homokaryotypes), we found almost no candidate SNPs associated with size variation (0 for αα individuals, and up to 3 SNPs when the FDR was lowered to p=0.01 for the ββ individuals, Fig. S21). We ran gene ontology using two data sets: the candidates identified by GWAS and all genes present in *Cf-Inv(1)*. Both analyses showed an enrichment in several biological processes all consistent with large differences in wing size and life-history, such as morphogenesis, muscle development or neural system development (Table S6-S7).

Given the extreme temperature range inhabited by *C. frigida* (temperate to subarctic) we also investigated thermal adaptation. We evaluated the recovery time after a chill coma in the F2s used to build the linkage map. Cold shock resistance localized to a QTL on LG4, which explained about 13% of the variation (Fig. S22). The main peak was located on LG4 around 25-28Mb. This broad QTL encompassed multiple outliers SNPs associated with climatic variation, and multiple annotated genes, among them two heat shock proteins, which may represent relevant candidates for thermal adaptation (Uniprot P61604 at position 25,128,992 and P29844 at position 26,816,283). This peak was located between the two putative inversions *Cf-Inv(4.2)* and *Cf-Inv(4.3)*, and there was a secondary peak at 8MB, the putative breakpoint of *Cf-Inv(4.1)*.

## Discussion

Analyses of more than 1, 400 whole genomes of *C. frigida* flies revealed four large chromosomal inversions affecting a large fraction of the genome (36.1Mb, 15%), and three low recombining genomic regions. These megabases-long stretches of the genome appear to play a predominant role in shaping genetic variation across two large scale environmental gradients as well as heterogeneous patchy habitats. Yet different inversions showed contrasting patterns, which may be related to different selective forces acting on them. In particular, the newly-discovered inversions on LG4 displayed clinal variation along a geoclimatic gradient. In contrast, the largest inversion *Cf-Inv(1)* was associated with body size and covaried at a fine geographic scale with wrackbed habitat characteristics, confirming previous work (Day et al. 1983; Butlin and Day 1985; Mérot et al. 2018). Below, we discuss how our results provide new insights into the evolutionary role played by recombination-limited regions including inversions, and how our data suggest that those regions are involved in local adaptation at different geographic scales in the face of high gene flow.

### • Low-coverage sequencing provides insights into genetic variation across a species range and individual genomes

Studying all aspects of genetic variation across a species range is more accurate and powerful when sampling encompasses both fine and coarse geographical scales across multiple environmental conditions. When searching for signatures of adaptation or putative inversions, high density genetic markers are required to identify patterns (Fuentes-Pardo and Ruzzante 2017). This creates the need to balance effort across the number of samples, the portion of the genome sequenced (*i.e*., reduced representation or whole genome sequencing), and the depth of sequencing. To maximise insights, we sequenced the whole genome of 1,446 wild-collected flies, but reduced individual coverage to about 1.4X. Simulations have shown that sequencing many samples at low depth (1X) provides robust estimates of population genetic statistics, namely allele frequencies, F_ST_, and other population parameters, and may be more powerful than sequencing few samples at higher depth (Alex Buerkle and Gompert 2013; Lou et al. 2020). Consequently, this strategy has been used efficiently in a few pioneer studies in human genomics (Martin et al. 2020), conservation genomics (Therkildsen et al. 2019) and population genomics (Clucas et al. 2019). Additionally, thanks to a low cost barcoding library preparation (Therkildsen and Palumbi 2017), individual information was retained, which allowed parameters that require this information (LD, Hobs) to be accurately calculated as well as the use of phenotypic association studies. Importantly, allele frequencies were also unbiased by *a priori* or unbalanced pooling as may happen in pool-seq (Fuentes-Pardo and Ruzzante 2017), and any grouping could be subsequently chosen for the analyses.

Individual whole genome sequencing at low coverage allowed us to uncover the genetic structure associated with inversions in *C. frigida* and to analyze environmental parameters and phenotypes potentially associated with those inversions. First, the large sample size brought power to make the most of a recently developed method of indirect inversion detection (Li and Ralph 2019; Huang et al. 2020). For instance, we would probably have missed the inversion(s) *Cf-Inv(4.2/4.3)* with smaller sample size, since the rare homokaryotype frequency was below 2% (26/1446 individuals). Second, the high density of markers along the genome provided accurate locations for the major inversions although characterizing the exact breakpoints was impossible without long-read sequencing (Ho et al. 2019). Third, the retention of individual information allowed us to split the dataset into subgroups of karyotypes, as determined from the analyses of sequences, and to characterize LD, heterozygosity, nucleotide diversity and the differentiation within and between karyotypes for all inversions. This aspect was critical to our study and allowed us to describe contrasting patterns from a geographic and ecological point of view.

### • Polymorphic inversions structure within-species genetic diversity

Despite the wide geographic area covered, the major factor shaping genetic variation in *C. frigida* was not geographic distance but chromosomal inversions. Despite more than 1,500km (or 3,000 km of coastal distance) between the most distant populations, geographic genetic differentiation was very weak (Maximal F_ST_ <0.02). This is much lower than other coastal specialised insects such as the saltmarsh beetle *Pogonus chalceus* (F_ST_ ~0.2, (Van Belleghem et al. 2018) but comparable to small Dipterans with large distributions like *Drosophila melanogaster* or *D. simulans* which typically exhibit F_st_ around 0.01-0.03, probably resulting from both high migration rate and large effective population size *N_e_* (Machado et al. 2016; Kapun et al. 2020). Despite this weak genetic structure, we detected a strong signal of isolation by distance indicating that dispersal among populations and subsequent gene flow decreases with distance. Furthermore, our analyses also showed that the least cost distance along the coastline better explained genetic variation than Euclidean distance. This isolation by resistance pattern probably results from a stepping stone dispersal process (Gandon and Rousset 1999) where the absence of suitable habitat patches in mainland and marine areas drives gene flow along the shore and constraints genetic connectivity.

In contrast with the overall weak geographic genetic structure, the frequencies of the different inversion arrangements were highly differentiated. Differentiation was restricted to the inverted regions, with fixed allelic differences between arrangements. Such a high genotypic divergence between alternative arrangements is comparable to many other ancient inversions (Hoffmann and Rieseberg 2008; Wellenreuther and Bernatchez 2018), and reflects the accumulation of neutral and non-neutral mutations between two sequences that experience a reduction in recombination (Berdan et al. 2021). Divergence was stronger between arrangements of *Cf-Inv(1)* than between arrangements of the LG4 inversions. Several non-exclusive hypotheses can explain this. First, it is possible that *Cf-Inv(1)* is older, leaving more time for mutations to accumulate. Second, *Cf-Inv(1)* is a complex structural variant, which involved at least three separate inversion events (Aziz 1975; Day et al. 1982), and such complexity is known to suppress double crossovers and gene conversion, which maintain some exchange in simpler inversions (Korunes and Noor 2019). Finally, the distribution of karyotypes across the populations will strongly affect mutation accumulation by dictating the frequency of the arrangements (and thus their *N_e_*) as well as the extent of recombination suppression. *Cf-Inv(1)* is polymorphic in all populations studied with a higher than expected proportion of heterokaryotypes. Conversely, *Cf-Inv(4.1)* has high frequencies of opposing homokaryotypes at each end of the cline. It is probably a combination of age, extent of gene flux (i.e., double crossing over and gene conversion between arrangements) and karyotype distribution that explains the variation in differentiation between our inversions.

### • Chromosomal inversions are involved in adaptation to heterogeneous environments

Across geographic and ecological gradients, inversions may contribute strongly to genetic differentiation and often appear as islands of differentiation (Hoffmann et al. 2004). For instance, in the mosquito *Anopheles gambiae*, genetic differentiation along a latitudinal cline is almost entirely concentrated in two inversions (Cheng et al. 2012). In the marine snail *Littorina saxatilis*, genetic variation between habitats is largely driven by several inverted regions (Morales et al. 2019). *Coelopa frigida* follows this trend: pairwise F_ST_ values between populations are higher in inverted regions compared to collinear regions, albeit at a different geographic scale for the different inversions. Along the North-South gradient, differentiation between populations was higher and isolation by distance was stronger in *Cf-Inv(4.1)* and *Cf-Inv(4.2/4.3)* than in collinear regions. F_ST_ based on SNPs in an inverted region combined two levels of genetic variation because differentiation between populations was driven by frequency variation at each highly differentiated arrangement. Such frequencies showed strong latitudinal clines, resembling the clines observed for several inversions in *Drosophila* that are maintained by selection-migration balance (Kapun et al. 2016). In sharp contrast, the genetic differentiation in the inverted region *Cf-Inv(1)* did not depend on geographic distances among populations. This pattern was related to the heterogeneous frequency of the α and β arrangements, which vary at a fine spatial scale but do not vary clinally. Yet, both the clines of *Cf-Inv(4.1)/ Cf-Inv(4.2/4.3)* and the heterogeneity of *Cf-Inv(1)* contrasted with the homogeneous frequency of collinear variants, supporting the hypothesis that inversion distribution reflects spatial variation in selection pressures.

Genotype-environment associations (GEA) confirmed the putative role of inversions in adaptation to small scale and large scale variation of ecological conditions in *C. frigida*. Here, one question that may arise is whether the SNPs located in an inverted region are more likely to be detected as outliers than collinear SNPs. We avoided such artefacts by following the guidelines and best practices from Lotterhos (2019) that used simulations to confirm the absence of bias when inversions or low recombining regions were neutral. However, genome scan analyses are still more likely to detect adaptive regions with strong divergence that are resistant to swamping by migration, while dispersed, transient or small-effect adaptive alleles are harder to detect (Yeaman 2015). Moreover, because of the high linkage disequilibrium associated with an inversion, several SNPs may not be causative but simply linked to an adaptive variant. Hence, the high density of outlier SNPs in inverted regions neither means that they are full of adaptive alleles, nor that they are the only variants relevant for local adaptation. Nevertheless, the strengths of association between environment and the frequencies of some SNPs found in the inverted regions, as well as the association between environment and inversion frequencies, support inversions as major and true players of adaptation to heterogeneous environments in *C. frigida*. As such, the seaweed fly *C. frigida* joins an accumulating number of studies pioneered by Dobzhansky (1947; 1948) which have provided examples of species carrying several ecologically-relevant inversions that are involved in local adaptation despite high gene flow (Anderson et al. 1991; Schaeffer 2008; Joron et al. 2011; Cheng et al. 2012; Kapun et al. 2016; Kirubakaran et al. 2016; Lindtke et al. 2017; Wellenreuther and Bernatchez 2018; Kapun and Flatt 2019; Huang et al. 2020). All of this is consistent with a model in which inversions are particularly relevant for adaptation with gene flow, because they preserve linkage between locally adapted alleles (Kirkpatrick and Barton 2006; Charlesworth and Barton 2018) and/or coadaptive epistatic alleles (Dobzhansky and Dobzhansky 1970; Charlesworth and Charlesworth 1973). However, although each inversion contains hundreds of genes, identifying multiple coselected or coadapted loci remains challenging because of LD, and calls for future experimental or transcriptomic work dissecting genetic variation in inversions.

In many empirical cases, when several inversions are found in the same species, they tend to vary along the same environmental axis. For instance, in the silverside fish *Menidia menidia*, several inverted haploblocks covary along a latitudinal gradient (Tigano et al. 2020; Wilder et al. 2020). The same tendency is observed for three inversions differentiating mountain and plain African honeybees *Apis mellifera scutellata* (Christmas et al. 2019), and dune and non-dune ecotypes of the sunflower *Helianthus petiolaris* (Huang et al. 2020; Todesco et al. 2020). In contrast, for *C. frigida*, we observed two contrasting evolutionary patterns: The inversion *Cf-Inv(1)* was associated with wrackbed characteristics and composition, which represent patchy habitats at a fine geographic scale. It also functions as a supergene for body size, a trait which is usually polygenic yet appears in *C. frigida* to be controlled largely, if not entirely, by this inversion. The ecological and phenotypic associations are consistent with previous work on European and American populations (Day et al. 1983; Butlin and Day 1985; Berdan et al. 2018; Mérot et al. 2018). They reflect how the quality, composition and depth of the wrackbed, possibly reflecting its stability, differently favour the opposite life-history strategies associated with the inversion. The β arrangement provides quick growth and smaller size while the α arrangement provides high reproductive success linked to a larger size but at the expense of longer development time. This ecologically-related trade-off combined with heterozygote advantage results in strong balancing selection (Butlin 1983; Mérot, Llaurens, et al. 2020). Conversely, the inversions *Cf-Inv(4.1)* and *Cf-Inv(4.2/4.3)* show no deviation from Hardy-Weinberg disequilibrium and display a strong geographic structure along a latitudinal cline. As *Cf-Inv(4.1)* and *Cf-Inv(4.2/4.3)* are associated with climatic variables, we suggest that they possibly play a role in thermal adaptation. Additional support for this hypothesis come from the close proximity of *Cf-Inv(4.1)* and *Cf-Inv(4.2/4.3)* with a QTL for recovery after chill coma although we cannot exclude that the presence and the position of that QTL may suffer from mapping bias caused by low recombination (Noor et al. 2001). To summarize, these inversions describe contrasting patterns driven by different shapes of selection, with *Cf-Inv(1)* being a cosmopolitan polymorphism under balancing selection, while *Cf-Inv(4.1)* and *Cf-Inv(4.2/4.3)* represent geographically-structured polymorphisms, possibly under spatially variable selection.

### • Exploring low-recombination regions: what are they and why do they matter?

In addition to the aforementioned inversions, we also identified additional regions that spanned large fractions of each chromosome (6 to 16MB) and were characterized by distinct haploblocks, high LD, and low recombination. With the current data, we can only speculate about what those regions are and what are the mechanisms underlying the observed patterns. Different types of data suggest different answers to this question. For example, the enrichment in transposable elements (Fig. S7) may indicate pericentromeric regions or transposon-rich centromeres, which are challenging to assemble and characterize (Talbert and Henikoff 2020). However, we did not observe the typical enrichment of AT content (Fig. S7). The landscape of nucleotide diversity was also very heterogeneous: parts of those low recombining regions are deserts of diversity (Fig. 3C), as expected under increased selection at linked sites (also called “linked selection”, (Cutter and Payseur 2013)), which leads to genetic hitchhiking around loci affected by positive or negative selection (Begun and Aquadro 1992; Charlesworth 1996). Yet, peaks of high diversity are observed in *Cf-Lrr(2)* and *Cf-Lrr(5)*. These may reflect signatures of associative overdominance (Ohta 1971), due to masking of recessive deleterious loci in heterozygotes, as observed in some low recombining regions of human and *Drosophila* genomes (Becher et al. 2020; Gilbert et al. 2020). Recent admixture from related lineages can also form distinct haploblocks (Li and Ralph 2019) and would generate similar patterns of high diversity but we consider this hypothesis unlikely in our case as no sympatric sister-species is known. Another possibility is that haploblocks coinciding with peaks of diversity are misassembled structural variants embedded in a low recombining region, such that haploblocks that are seemingly separated could be adjacent. Our reference genome was scaffolded and ordered based on a linkage map from one family. Hence, inversions that were heterozygous in the mother, as well as any low recombining regions, could cluster into large regions with low rates of crossing-over in the map, where marker ordering may be less accurate. Additional data such as long-reads or connected molecules like Hi-C are needed to improve the quality of the assembly in those specific areas and better characterize their DNA content. Despite these cautionary notes, our analysis provides an early annotation of regions that do not behave like the rest of the genome in terms of geographic genetic structure and association with environmental factors.

The low recombining regions may also play a role in shaping the distribution of adaptive and non-adaptive variation since they differentiated populations more strongly than collinear regions. One potential reason would be increased variance in F_ST_ statistics in low recombining regions that can emerge even under a purely neutral model (Booker et al. 2020). The effect of selection at linked sites, which reduces diversity in low recombining regions, is also known to inflate differentiation, sometimes repeatedly between different pairs of populations (Burri et al. 2015; Hoban et al. 2016). While these processes may explain extreme F_ST_ values in low diversity and low recombining subregions, they are unlikely to explain the pattern that we observed in high diversity subregions, at least in *Cf-Lrr(2)* and *Cf-Lrr(5)*, which include GEA outliers as well as haplotypic variants whose frequency correlates with environmental variation. Without certainty about the mechanisms behind the reduced recombination, we can only propose hypotheses about the evolutionary processes at play. If these regions are complex or misassembled structural variants, they would represent additional adaptive chromosomal rearrangements contributing to adaptation in *C. frigida*, with different arrangements bearing one or several locally adapted alleles. If those regions are centromeric, or simply rarely recombining, they would highlight the importance of selection at linked sites in structuring intraspecific variation and the relevance of low recombining regions in protecting locally adapted alleles. Evidence for an important evolutionary role of low recombining regions is increasingly reported and we should analyse genomic landscapes in the light of recombination. For instance, in three-spine stickleback (*Gasterosteus aculeatus*), putatively adaptive alleles tend to occur more often in regions of low recombination in populations facing divergent selection pressures and high gene flow (Samuk et al. 2017). Similarly, regions of low recombination are enriched in loci involved in parallel adaptation to alpine habitat in the Brassicaceae *Arabidopsis lyrata* (Hämälä and Savolainen 2019). To what extent LD in low recombination regions affect such inferences yet remains an open question (Stevison and McGaugh 2020). Some statistics are biased by recombination heterogeneity (e.g. outliers based on F_ST_ in sliding-windows(Booker et al. 2020) or on PCA (Lotterhos 2019), QTL from mapping families (Noor et al. 2001)) but other approaches appear robust when following best practices (e.g. genotype-environment associations, selective sweep detection (Lotterhos 2019)). Overall, further work is needed, both on the methodological and empirical points of view, in order to better understand the contribution of recombination heterogeneity in structuring intraspecific variation and modulating migration-selection balance.

## Conclusion

Our findings support the growing evidence that large chromosomal inversions play a major evolutionary role in some organisms characterised by extensive connectivity across a large geographical range. In this flying insect, as in several marine species, migratory birds, and widespread plants, chromosomal rearrangements strongly affect genetic diversity and represent a key component of the genetic architecture for adaptation in the face of gene flow. Critically, the different inversions are under different selective constraints across a range of geographical scales and contribute to adaptation to different environmental factors. Thus, inversions appear to be an architecture that allows some species to cope with gene flow as well as various sources and scales of environmental heterogeneity. While inversions present one solution to the problem of adaptation with gene flow it is still unknown how prevalent inversions are in nature. This is because structural variants are just beginning to be characterized in non-model species. In one of the best-studied clades, *Drosophila*, the answer is contradictory, closely related species *Drosophila melanogaster* and *Drosophila simulans* exhibit respectively > 500 vs. only 14 polymorphic inversions while both species are ecologically successful, and distributed worldwide with high connectivity between populations (Aulard et al. 2004). This is possibly due to the dichotomous nature of reduced recombination. While reduced recombination holds together complexes of adaptive alleles it also hampers the generation of new (and potentially adaptive) allele combinations and reduces the efficiency of purifying selection in linked regions (Felsenstein 1974). Overall, our analysis highlights the importance of regions of low recombination in structuring adaptive and non-adaptive intraspecific genetic variation. With recombination varying both along the genome and between individuals or haplotypes, inversions may represent only the simplest aspect of the complex relationship between recombination, selection and gene flow that we are just starting to uncover through the prism of structural variants (Stapley et al. 2017). By optimising whole genome sequencing to include many individuals across a species range as done here, future work will have the possibility to better understand how the interplay between structural variation and recombination may matter for the evolution of biodiversity.

## Methods

### A reference genome assembly for *Coelopa frigida*

To generate a reference genome, we sequenced female siblings of *C. frigida* homozygous αα for the inversion *Cf-Inv(1)*, obtained by three generations of sib-mating from parents collected in St Irénée (QC, Canada). A pool of DNA from three siblings was sequenced on four cells of Pacific Biosystems Sequel sequencer, producing 16.1 Gbp (~64x coverage) of long reads, and one sibling was sequenced with 10xGenomics Chromium on 1 lane of an Illumina HiSeqXTen sequencer, yielding 82 Gbp (~300x of coverage) of 150bp paired-end linked-reads. Long reads were assembled using the Smrt Analysis v3.0 pbsmrtpipe workflow and FALCON (Chin et al 2013), resulting in 2959 contigs (N50 = 320 kb), for a total assembly size of 233.7 Mbp. This assembly was polished by using the linked reads, first by correcting for sequence errors with Pilon (Walker et al. 2014) and, second, by correcting for misassemblies with Tigmint (Jackman et al. 2018). The resulting assembly consisted of 3096 contigs (N50 = 320 kb). The contigs were scaffolded using the long-range information from linked-reads with ARKS-LINKS (Coombe et al. 2018), resulting in 2539 scaffolds (N50 = 735kb). Scaffolds were anchored and oriented into chromosomes using *Chromonomer* (Catchen et al. 2020), based on the order of markers in a linkage map (see below). The final assembly consisted of 6 chromosomes and 1832 unanchored scaffolds (N50 = 37.7Mb) for a total of 239.7Mb (195.4 Mb into chromosomes). The completeness of this reference was assessed with BUSCO version 3.0.1 (Simão et al. 2015). The genome was annotated by mapping a transcriptome assembled from RNA sequences obtained from 8 adults (split by sex and by karyotype at the *Cf-Inv(1)* inversions), 4 pools of 3 larvae and pools of *C. frigida* at different stages. The transcriptome was annotated using the Triannotate pipeline. More details are provided as supplementary methods.

### A high-density linkage map and QTL analyses

#### • Sequencing, genotyping and building the map

We generated an outbred F2 family of 136 progenies by crossing two F1 individuals of *C. frigida* obtained by crossing wild individuals collected in Gaspésie (QC, Canada). The mother of the F2 family was homozygous αα at *Cf-Inv(1)*. The progeny, both parents, and two paternal grandparents were sequenced using a double-digest restriction library preparation (ddRAD-seq) using ApeK1 on an IonProton (ThermoFisher), with greater depth for the parents. Reads were trimmed and aligned on the scaffolded assembly with bwa-mem. Genotype likelihoods were obtained with SAMtools mpileup. Only markers with at least 3X coverage in all individuals were kept. More details are provided in supplementary methods

A linkage map was built using *Lep-MAP3* (Rastas 2017) following a pipeline available at https://github.com/clairemerot/lepmap3_pipeline. Markers with more than 30% of missing data, non-informative markers, and markers with extreme segregation distortion (χ^2^ test, *P* < 0.001) were excluded. Markers were assigned to linkage groups (LGs) using the *SeparateChromosomes* module with a logarithm of odds (LOD) of 15, a minimum size of 5 and assuming no recombination in males, as is generally the case in Diptera (Satomura et al. 2019). This assigned 28,615 markers into 5 large LGs, and 25 sex-linked markers into 2 small LGs than were subsequently merged as one (LG6). Within each LG, markers were ordered with 5 iterations of the *OrderMarker* module. The marker order from the run with the best likelihood was retained and refined 3 times with the *evaluateOrder* flag with 5 iterations each. When more than 1000 markers were plateauing at the same position, all of them were removed, as suggested by *Lep-MAP* guidelines. Exploration for more stringent filtering or different values of LOD resulted in collinear maps.

#### • Estimating recombination rate

To estimate recombination rate across the genome, we compared the positions of the markers along the genetic map with their position along the genome assembly with MAREYMAP (Rezvoy et al. 2007). Local recombination rates were estimated with a Loess method including 10% of the markers for fitting the local polynomial curve.

#### • Quantitative trait locus (QTL) analysis

All individuals used to build the map were scored for recovery at room temperature after a chill coma induced by holding them for 10 minutes at −20°C. We distinguished three categories: “0”, the fly stands immediately or in less than 5 minutes; “1”, the fly recovers with difficulty after 5 to 15 minutes; “2”, the fly has not recovered after more than 15 minutes. A phased map was obtained by performing an additional iteration of the *OrderMarker* module. QTL analysis was carried out using R/qtl (Broman et al. 2003). LOD scores correspond to the −log_10_ of the associated probabilities between genotype and phenotype with the Haley-Knott method. The LOD threshold for significance was calculated using 1000 permutations.

### Population level sequencing

#### • Sampling and characterisation of size and karyotype

We analysed 1446 adult *C. frigida*, sampled at 16 locations spanning over 10° of latitude (Fig.1A) in September/October 2016. Sampling, genotyping and phenotyping are described in detail in Mérot et al. (2018). Size was estimated using wing length as a proxy for 1426 flies. Genomic DNA was extracted using a salt extraction protocol (Aljanabi and Martinez 1997) with a RNase A treatment (Qiagen). A total of 1438 flies were successfully genotyped at the *Cf-Inv(1)* inversion using a molecular marker (Mérot et al. 2018).

#### • Library preparation, sequencing and processing of the sequences

Whole genome high-quality libraries were prepared for each fly by adapting a protocol described in (Baym et al. 2015; Therkildsen and Palumbi 2017) and detailed in Supplementary Materials. Briefly, DNA extracts were quantified, distributed in 17 plates with randomisation (96-well) and normalised at 1ng/μL. Each sample extract underwent a step of tagmentation, which fragments DNA and incorporates partial adapters, two PCR steps that attached barcode sequences (384 combinations) while amplifying the libraries, and a size selection step using an Axygen magnetic bead cleaning protocol. Final concentrations and fragment size distributions were estimated to combine equimolar amounts of 293 to 296 libraries into 5 separate pools. Sequencing on 5 lanes of Illumina HiSeq 4000 yielded an average of 327Mb per sample, which resulted in approximately 1.4X coverage (range: 121-835Mb, 0.5X-3.5X)

Paired-end 150bp reads were trimmed and filtered for quality with FastP (Chen et al. 2018), aligned to the reference genome with BWA-MEM (Li and Durbin 2009), and filtered to keep mapping quality over 10 with Samtools v1.8 (Li et al. 2009). Duplicate reads were removed with MarkDuplicates (PicardTools v1.119.) We realigned around indels with GATK IndelRealigner (McKenna et al. 2010) and soft clipped overlapping read ends using clipOverlap in bamUtil v1.0.14 (Breese and Liu 2013). Reads, in bam format, were analysed with the program ANGSD v0.931 (Korneliussen et al. 2014), which accounts for genotype uncertainty and is appropriate for low coverage whole genome sequencing (Lou et al. 2020). Input reads were filtered to remove low-quality reads and to keep mapping quality above 30 and base quality above 20.

As a first step, we ran ANGSD to estimate the spectrum of allele frequency, minor allele frequency, depth and genotype likelihoods on the whole dataset. Genotype likelihoods were estimated with the GATK method (-GL 2). The major allele was the most frequent allele (-doMajorMinor 1). We filtered to keep positions covered by at least one read in at least 50% of individuals, with a total coverage below 4338 (3 times the number of individuals) to avoid including repeated regions in the analysis. From this list of variant and invariant positions, we selected SNPs with a minor allele frequency (MAF) of above 5% and subsequently used this list with their respective major and minor alleles for most analyses (PCA, inversion detection, F_ST_, genotype-environment associations). Using PLINK 1.9, we selected a subset of SNPs pruned for physical linkage, removing SNPs with a variance inflation factor greater than two (VIF>2) in 100 SNP sliding windows shifted by 5 SNPs after each iteration.

#### • Principal component analysis (PCA) and inversion detection

An individual covariance matrix was extracted from the genotype likelihoods in beagle format using PCAngsd (Meisner and Albrechtsen 2018) and decomposed into principal components with R, using a scaling 2 transformation, which adds an eigenvalue correction (Legendre and Legendre 1998). To analyse genetic variation along the genome, we performed the same decomposition in non-overlapping windows of 100 SNPs. For each “local PCA”, we analysed the correlation between PC1 scores and PC scores from the PCA performed on the whole genome. This allowed us to locate two (inversion) regions underlying the structure observed on PC1 and PC2 (Fig2A). We set the boundaries of those regions as windows with a coefficient of correlation above one standard deviation.

To scan the genome for other putative inversions or non-recombining haploblocks, we used the R package *Lostruct* (Li and Ralph 2019) which measures the similarity between local PCA (PC1 and PC2 for each 100SNP window) using Euclidean distances. Similarity was mapped using multidimensional scaling (MDS) of up to 20 axes. Clusters of outlier windows (presenting similar PCA patterns) were defined along each MDS axis as those with values beyond 4 standard deviations from the mean, following Huang et al. (2020). Adjacent clusters with less than 20 windows between them were pooled, and clusters with less than 5 windows were not considered. Different window sizes (100 to 1000), different subset of PCs (1 to 3 PCs) and different thresholds yielded consistent results. A typical signature of a polymorphic inversion is three groups of individuals appearing on a PCA: the two homokaryotypes for the alternative arrangements and, as an intermediate group, the heterokaryotypes. All clusters of outlier windows were thus examined either by a PCA as single blocks, or divided into several blocks when discontinuous. We then used K-means clustering to identify putative groups of haplotypes. Clustering accuracy was maximised by rotation and the discreteness was evaluated by the proportion of the between-cluster sum of squares over the total.

#### • Inversion analysis

For the four inversions (*Cf-Inv(1), Cf-Inv(4.1), Cf-Inv(4.2/4.3)*), K-means assignment on PC scores was used as the karyotype of the sample. Differentiation among karyotypes was measured with F_ST_ statistics, using ANGSD to estimate joint allele frequency spectrum, realSFS functions to compute F_ST_ in sliding windows of 25KB with a step of 5KB, and subsampling the largest groups to balance sample size. Observed proportion of heterozygotes (Hobs) was calculated for each karyotype and each SNP using the function -doHWE in ANGSD, and then averaged across sliding windows of 25KB with a step of 5KB using the R package *windowscanr*. Nucleotide diversity (π) within each arrangement, and nucleotide divergence (dxy) between arrangements was calculated in sliding windows of 25KB (step 5KB) considering all positions (variants and invariants), controlling for missing positions and using the function -doThetas (ANGSD) and the script https://github.com/mfumagalli/ngsPopGen/blob/master/scripts/calcDxy.R, following the recommendation of Korunes and Samuk (2021). For each 25kb window, nucleotide divergence was corrected for within arrangement genetic variation by subtracting the mean of the nucleotide diversity in both arrangements. The 95% confidence intervals were estimated by bootstrapping the data using per-window corrected d_XY_ estimates (1,000 replicates). Based on this value of corrected *d_XY_* between each inversion’s arrangements calculated on non-coding windows in inverted regions, we estimated an approximate time of divergence using a constant molecular clock. We assumed a mutation rate comparable to *Drosophila*, with *μ* equal to 5 ×10^−9^ mutations per base per generation (Assaf et al. 2017). Given that generation time in *C. frigida* strongly varies, from 8 to 20 days at 25°C up to months in colder conditions, we considered a range of 5 to 10 generations per year. As arrangements are expected to keep some gene flux after the formation of the inversion, due to double crossing-over and gene conversion (Navarro et al. 1997; Korunes and Noor 2019), the age estimates should be considered as a minimum value.

#### • Linkage disequilibrium (LD)

Intrachromosomal LD was calculated among a reduced number of SNPs, filtered with more stringent criteria (MAF > 10%, at least one read in 75% of the samples). Pairwise R^2^ values were calculated with NGS-LD (Fox et al. 2019) based on genotype likelihood obtained by ANGSD, and grouped into windows of 1MB. Plots display the 2^nd^ percentile of R^2^ values per pair of windows. For LG1 and LG4, R^2^ was calculated, first within all samples, then within individuals homozygous for the most common orientation of each inversion, subsampling the largest groups to balance sample size, and plotted by windows of 250kb.

#### • Geographic structure

*F_ST_* differentiation between all pairs of populations was estimated based on the allele frequency spectrum per population (-doSaf) and using the realSFS function in ANGSD. Positions were restricted to the polymorphic SNPs (> 5% MAF) previously polarised as major or minor allele (options –sites and –doMajorMinor 3), and which were covered in at least 50% of the samples in a given population. Populations were randomly subsampled to a similar size of 88 individuals. The weighted *F_ST_* values between pairs of population were computed by including either all SNPs, LD-pruned SNPs, or SNPs from a region of interest (inversions/low recombining regions) or SNPs outside those regions (collinear SNPs).

Isolation by distance (IBD) was tested for each subset of SNPs using a linear model in which pairwise genetic distance (*F_ST_*/(1-*F_ST_*)) was included as the response variable and geographic Euclidean distance was incorporated as an explanatory term. Isolation by resistance (IBR) refers to constrained dispersal due to environmental features that limit movement and was tested in the same way as IBD, except that physical distances were calculated along the shoreline, assuming that the open water or mainland may oppose dispersal of *C. frigida*. The distance via least cost path was measured through areas of the map between −40 meters of depth and 20 meters of altitude using the R package *marmap*. Both models of IBD and IBR were compared to a null model using an ANOVA F-test, and to each other using adjusted *R*^2^ and AIC. To compare IBD and IBR patterns in each inversion/low recombining region to the collinear genome, we built a full model explaining pairwise genetic distances by physical distances and genomic region (collinear vs. inversion) as a cofactor, and assessed the significance of the interaction term as well as the direction of the interaction slope coefficient. We repeated this analysis 100 times with randomly chosen collinear regions including the same number of contiguous SNPs as each inversion/low recombining regions. This provided a distribution of the significance of the interaction term and its slope coefficient (Fig. S13). For *Cf-Inv(1)*, no contiguous block with the same number of SNPs could be found in the genome, hence we gathered 3 blocks of 1/3 the number of SNPs in each of the 100 random replicates.

Finally, we examined the direct association between inversions and latitude, treating inversions as single bi-allelic loci. The association was tested by a GLM with a logistic link function for binomial data, the response variable being the number of individuals carrying/not carrying the minor arrangement and the explanatory variable being latitude. To assess whether this association deviates from null expectations, we randomly sampled 1000 SNPs, with an average frequency similar to each inversion, and built 1000 full models explaining frequency by latitude and genomic region (a collinear SNP vs. an inversion) as a cofactor, and assessed the significance of the interaction term (Fig. S15).

#### • Environmental associations

Environment at each location was described by large scale climatic/abiotic conditions and local wrackbed characteristics (Table S1), as described in Mérot et al. (2018). Large scale conditions included annual means in precipitation, air temperature, sea surface temperature, sea surface salinity and tidal amplitude extracted from public databases. Wrackbed characteristics were measured upon collection and included abiotic variable (depth, internal temperature and salinity) and algal composition (relative proportions of Laminariaceae, Fucaceae, Zoosteraceae, plant debris and other seaweed species). Correlation between variables was tested with a Pearson correlation test, and variation was reduced by performing a PCA for each group of correlated environmental variables (climatic, salinity/tidal amplitude, abiotic characteristic of the wrackbed, algal composition) and retaining significant PCs following the Kaiser-Guttman and Broken Stick criteria (Borcard et al. 2011) (see Fig. S16).

Minor allele frequency was calculated for each population from the list of SNPs previously polarised as major or minor allele (–sites and –doMajorMinor 3), and covered by at least 50% of the individuals in each population. Allelic frequency was thus estimated with confidence (>50X of coverage at population level) for a total of 1,155,978 SNPs. A genetic environment association (GEA) which evaluated SNPs frequencies as function of environmental variables was performed through a combination of two methods as recommended by de Villemereuil et al. (2014): (i) latent factor mixed models (LFMM2; Frichot et al. 2013; Caye et al. 2019), (ii) Bayes factor (BAYPASS; Gautier 2015). Those three methods had also been shown to be robust to the presence of large inversions (Lotterhos 2019).

LFMM was run with the R package *lfmm2* (Caye et al. 2019) on the full set of 1,155,978 SNPs, using a ridge regression which performed better in simulations including inversions (Lotterhos 2019), and parametrized using a K-value of 4 latent factors (as evaluated from a PCA on a LD-pruned dataset). False discovery rate was assessed following the recommendations of François et al. (2016), using a Benjamini-Hochberg correction. Using Baypass v2.2 (Gautier 2015), a Bayes factor (BF), evaluating the strength of an association between SNP frequency and environment, was computed as the median of three runs under the standard model on the full set of 1,155,978 SNPs. Environmental variables were scaled using the -scalecov option. We ran this analysis twice: first, without controlling for population structure and, second, by controlling with a covariance matrix extracted from an initial BayPass model run on the subset of LD-pruned SNPs without environmental covariables. To calculate a significance threshold for BF, we simulated pseudo-observed data with 10,000 SNPs using the “simulate.baypass” function and kept the 0.1% quantile as the significance threshold.For each GEA method, and the combination of the two, the repartition of candidate SNPs for association with environment inside and outside inversions/low recombining regions was compared to the original repartition of SNPs. Deviation from this original repartition was tested with a Fisher’s exact test, and the magnitude of the excess/deficit of outlier SNPs in each region of the genome was reported as the odd ratio.

We also compared the distribution of association scores in each inversion/low recombining region to the collinear genome. This test was performed on absolute values of the z scores from LFMM, using a generalised linear model with quasinormal family (square root link) and genomic region (collinear vs. inversion) as an explanatory factor. We repeated this analysis on 100 randomly chosen collinear blocks including the same number of SNPs as each inversion/low recombining regions times (Fig. S19). Finally, we examined the direct association between inversions and environment variables, treating each inversion as a single locus, as described above for latitude using GLM models and comparing to 1000 randomly drawn SNPs (Fig. S20).

#### • Phenotypic associations and gene ontology analysis

We performed a genome wide association study (GWAS) for wing size using ANGSD latent genotype model (EM algorithm, -doAssso=4) where genotype is introduced as a latent variable and then the likelihood is maximized using weighted least squares regression (Jørsboe and Albrechtsen 2020). We considered a false discovery rate (FDR) of 0.001. The GWAS was applied on the whole dataset (1,426 flies with size information) and then on each subset of homokaryotypes at the inversion *Cf-Inv(1)* (140 αα and 436 ββ flies with size information).

Using BEDtools, we extracted the list of genes overlapping with significantly associated SNPs, or within a window of 5kb upstream or downstream a gene. We then tested for the presence of overrepresented GO terms using GOAtools (v0.6.1, pval = 0.05) and filtered the outputs of GOAtools to keep only GO terms for biological processes of levels 3 or more, and with an FDR value equal below 0.1. We performed the same GO enrichment analysis for the list of genes found in the two largest inversions (*Cf-Inv(1)* and *Cf-Inv(4.1)*).

## Data availability

The genome assembly, GBS reads used to build the linkage map and WGS paired reads used for population genomics are available on NCBI under the projects, respectively (PRJNA688905, PRJNA689789, and PRJNA689963). Raw information about GBS and WGS samples is provided as supplementary tables S9 & S10. The pipelines for analyses of WGS data are available at https://github.com/enormandeau/wgs_sample_preparation and https://github.com/clairemerot/angsd_pipeline.

## Acknowledgments

We are very grateful to M. Lionard who sampled in Blanc Sablon and to L. Johnson, E. Tamigneaux, D. Malloch for their advice during fieldwork. We thank C. Babin and P. Berube for their support in the lab. B. Boyle and N. Therkildsen provided key advice about sequencing and libraries preparation. We thank Y. Dorant, M. Leitwein, and C. Rougeux for their help and advice with analyses, and C. Venney for proofreading the manuscript. We thank the editor and two anonymous reviewers for having thoroughly commented on an earlier version of the manuscript. The sequencing service was provided by the Norwegian Sequencing Centre (www.sequencing.uio.no), a national technology platform hosted by the University of Oslo and supported by the “Functional Genomics” and “Infrastructure” programs of the Research Council of Norway and the Southeastern Regional Health Authorities, by McGill Sequencing Platform and by the genomic platform at IBIS (University Laval http://www.ibis.ulaval.ca/). This research was supported by a Discovery research grant from the Natural Sciences and Engineering Research Council of Canada (NSERC) to L.B., by the Canadian Research Chair in genomics and conservation of aquatic resources held by L.B. and by the Swedish Research Council grant 2012-3996 to M.W. The genome assembly was supported by the Canada 150 Sequencing Initiative (CanSeq150), CFI and Genome Canada Technology Platform grants to J. R. C.M. was supported by a postdoctoral fellowship from the Fonds de Recherche Québec (FRQNT FRQS) and a Banting Postdoctoral Fellowship from the NSERC.

