## Supplementary materials for "Locally adaptive inversions modulate genetic variation at different geographic scales in a seaweed fly"

### Supplementary Methods 1: genome assembly

- **Sequencing**

To generate a reference genome, we sequenced sibling *Coelopa frigida* females from a 3-generation inbred family, obtained by crossing descendants from a pair of wild *Coelopa frigida* collected in St Irénée (QC, Canada). The inbred family was genotyped as homozygous for the α arrangement at the inversion *Cf-Inv(1R)* with a PCR assignment test developed in Mérot et al (2018). DNA was extracted individually from each sibling with a Phenol-Chloroform-Isopropanol extraction protocol adapted to preserve high-molecular weight DNA.

On a pool of DNA from three female siblings, long-read sequencing was performed on 4 cells of Pacific Biosystems Sequel sequencer at McGill University following standard procedures. It produced a total of 16.1 Gbp (~64x coverage) of sequencing data. One additional female from the same inbred family was sequenced following the 10xGenomics approach. High-molecular-weight DNA, with fragment size between 50 and 200kb, was loaded onto a Chromium controller chip, along with 10x Chromium reagents and gel beads following manufacturers recommended protocols. The resulting library was sequenced on one lane of an Illumina HiSeqXTen sequencer at McGill University. Sequencing yield 82 Gbp (~300x of coverage). The resulting linked-reads are illumina paired-end reads of 150bp, with barcode information linking short-reads belonging to the same molecules over long distances.

- **Assembling and polishing**

An initial assembly was carried out on the PacBio long reads using the Smrt Analysis v3.0 pbsmrtpipe analysis workflow tool from SMRT Link v5.0 (smrtlink-release_5.0.0.6792, pbsmrtpipe version: 0.51.2). The genome assembly and polishing was run with FALCON (Chin et al 2013), using the polished_falcon_fat pipeline option along with sequencing read metadata (data.xml), pipeline settings (preset.json) and output directory. From the default json settings the genome size was set to 255Mb (HGAP_GenomeLength_str) and seed coverage (HGAP_SeedCoverage_str) was set to 30X. The Falcon assembly method operates in two phases: First, overlapping sequence reads were compared to generate accurate consensus sequences with read N50 greater than 10Kbp. Next, overlaps between the corrected longer reads are used to generate a string graph. The graph is reduced so that multiple edges formed by heterozygous structural variation are replaced to represent a single haplotype. Contigs are formed by using the sequences of nonbranching paths. Two supplemental graph cleanup operations are defined to improve assembly quality by removing spurious edges from the string graph: tip removal and chimeric duplication edge removal. Tip removal discards sequences with errors that prevent 5′ or 3′ overlaps. Chimeric duplication edges may result from the raw sequence information or during the first sequence cleanup step and artificially increase the copy number of a duplication. In a second and final workflow stage, the polished_falcon_fat workflow will use the Quiver/Arrow consensus tool to perform the final error correction of the assembly and generated the polished assembly. This assembly yielded 2959 contigs (N50 = 320 kb), for a total assembly size of 233.7 Mbp.

This initial assembly was subsequently polished by using the linked reads. First, the contigs were corrected locally for sequence errors by running Pilon (Walker et al. 2014) with the following parameters (--changes --tracks --diploid --fix indels,gaps,local --mindepth 5) using Illumina paired-end short reads from the 10Xgenomics sequencing (~300X of the *C. frigida* genome) aligned with BWA-MEM on the initial assembly. Second, the information on long-range links provided by short-read barcodes was extracted using LongRanger (10xGenomics) and used by Tigmint (Jackman et al. 2018) with default parameters to correct misassemblies or break contigs. The resulting assembly included 3096 contigs with a N50 of 320 kb.

- **Scaffolding**

To scaffold the genome assembly, we used the program ARKS (D=true) and LINKS (Coombe et al. 2018; Yeo et al. 2018) with the following parameters (-k 30 -e 0.5 -l 3 -v 1), which relies on linked-reads to scaffold contigs. The resulting assembly included 2539 scaffolds with a N50 of 735 kb. At this stage, some scaffolds were manually broken following the identification of misassemblies in the preliminary analysis of the population genomic data and the genetic map. Finally, scaffolds were assembled into chromosomes using *Chromonomer* (Catchen et al. 2020), which anchors and orientates scaffolds based on the order of markers in a linkage map (see below). The default parameters were used. The final assembly accounted 6 chromosomes and 1832 unanchored scaffolds with a N50 of 37.7Mb for a total of 239.7Mb (195.4 Mb into chromosomes). The completeness of this reference was assessed with BUSCO version 3.0.1 (Simão et al. 2015).

### Supplementary Methods 2: transcriptome assembly

- **Rearing and crosses**

Larvae of *C. frigida* were collected from the field in May 2017 from Skeie (58.69733, 5.54083) and Østhassel (58.07068, 6.64346), Norway. All larvae were brought back to the Tjärnö Marine Laboratory in Strömstad, Sweden where they were raised to adulthood at 25°C. Eclosed adults from Østhassel were used to generate a development time series. Adults were introduced to three replicate containers with 50% *Saccharina latissima* and 50% *Fucus spp.* substrate. They were left for 24 hours to lay eggs after which they were removed. At this time egg samples were taken and samples were taken every subsequent 48 hours. Samples were stored in RNAlater in -20°C until extraction. The eclosed adults from Skeie were used to generate homokaryotypic crosses. Adult virgins were collected and one of the hind legs was removed for genotyping. DNA extraction and genotyping were performed as described in (Mérot et al. 2018) Three separate successful αα x αα and three separate successful ββ x ββ crosses were obtained. Thirty individuals (15 ♀ and 15 ♂) spread over these crosses were introduced to new containers containing 90 g *Saccharina latissima* and 45 g *Fucus spp.* to make an αα and a ββ line. Six days after the creation of these lines, two replicates of 3 larvae each were flash-frozen in liquid nitrogen and stored at -80°C until extraction. The adults that eclosed from these lines were used to make subsequent crosses (not described here) and then were flash-frozen in liquid nitrogen and stored at -80°C until extraction.

- **RNA extraction and library prep**

RNA from all samples were extracted using a TriZOL protocol. Briefly, 500 µl of TriZOL was added to each sample and then the sample was homogenized in a shaker using glass beads. The sample was then incubated at room temperature for 5 minutes after which it was centrifuged at 12,000 rcf 4°C for 10 minutes. The supernatant was transferred to a new tube and 100 µl of chloroform was added. After shaking the tube was incubated at room temperature for 3 minutes after which it was centrifuged at 10,000 rcf 4°C for 15 minutes. The upper aqueous phase was transferred to a new tube and 100 µl of phenol and 100 µl of chloroform were added. After shaking the sample was centrifuged for 7 minutes at 10,000 rcf 4°C. Then 500 µl of isopropanol was added and the sample was incubated for 10 minutes at room temperature. After a centrifugation step of 10 minutes at 12,000 rcf 4°C the pellet was washed with 500 µl of 75% EtOH, centrifuged at 7,500 rcf 4°C for 5 minutes, and air-dried for 10 minutes. Samples were re-suspended in 50 µl H20. After at least 24 hours all samples were further cleaned using the Zymo Clean & Concentrator Kit following manufacturers instructions.

- **Library preparation and Sequencing**

For the ontogeny series, we extracted RNA from 6 time points (eggs, 48 hours, 96 hours, 192 hours, 288 hours, and 384 hours). For each time point, we separately extracted samples from two of the three replicates. The concentration of these extractions was measured using a QBIT and equal amounts of RNA from each time point from one of the two samples were pooled to make a pool. Thus we had two pooled samples covering the same time distribution but not created from any of the same samples.

These pooled samples, as well as the 2 larval αα pooled samples, 2 larval ββ pooled samples, 2 αα adult males, 2 αα adult females, 2 ββ adult males, 2 ββ adult females were submitted to SciLifeLab in Uppsala, Sweden for library preparation and sequencing. All RNA was purified with Agencourt RNA clean XP before library preparation. Library preparation was done with the TruSeq stranded mRNA library preparation kit including polyA selection. Samples were sequenced along with 36 other libraries (not described here) on a NovaSeq S1 flowcell with 100 bp paired-end reads (v1 sequencing chemistry).

- **Transcriptome assembly**

Individual assemblies for each of the adult samples, both of the ontogenetic pools, both of the αα larval pools, and both of the ββ larval pools were done using Trinity v2.9.1 (11 assemblies total) (Haas et al. 2013). Prior to assembly, all reads were trimmed and adaptors removed using cutadapt 2.3 with Python 3.7.2. All assemblies were run through TransRate 1.0.1 (Smith-Unna et al. 2016) a quality assessment tool for *de novo* transcriptomes that looks for artefacts, such as chimeras and incomplete assembly, and provides individual transcripts and overall assembly scores. We retained all transcripts from each assembly classified by TransRate as 'good'. These contigs were then merged using CD-hit V4.8.1 (Fu et al. 2012) with a sequence identity threshold of 0.95, a word size of 10, and local sequence alignment coverage for the longer sequence at 0.005. Finally, the transcriptome was mapped to the genome assembly using GMAP version 2018-07-04 (Wu and Watanabe 2005). The mapping coordinates for each transcript were extracted and in the event that two transcripts mapped to the same coordinates, only the longer transcript was retained. The final transcriptome was annotated using the Trinotate pipeline (Grabherr et al. 2011).

### Supplementary methods 3: linkage map sequencing

- **Sequencing and genotyping**

We generate an outbred F2 family of 136 progenies by crossing two F1 individuals of *Coelopa frigida* from different crosses obtained from wild individuals collected in Gaspésie (QC, Canada). The mother of the F2 family was genotyped homozygous for the α arrangement at the inversion *Cf-Inv(1).* DNA from the progeny, both parents, and two paternal grandparents, was extracted following a salt-based extraction protocol (Aljanabi and Martinez 1997) with a RNAase A treatment. DNA was quantified using QuantiT Picogreen dsDNA Assay Kit (Invitrogen) and concentration was normalized to 10 ng/µl. Libraries were prepared and sequenced at the plateforme d'analyses génomiques of the Institut de Biologie Intégrative et des Systèmes (IBIS, Université Laval, Québec, Canada). Libraries were constructed following the procedure described by Elshire (Elshire et al. 2011) adapted for Ion proton sequencing as reported in Abed et al 2019 (Abed et al. 2019). Briefly, genomic DNA was digested with the restriction enzyme ApeK1 by incubating at 37°C for two hours followed by enzyme inactivation by incubation at 65°C for 20 min. Sequencing adaptors and a unique individual barcode were ligated to each sample using a ligation master mix including T4 ligase. The ligation reaction was completed at 22°C for 2 hours followed by 65°C for 20 min to deactivate the enzymes. Libraries were size-selected using a BluePippin prep (Sage Science), amplified by PCR. Libraries were prepared for sequencing using a Ion CHEF, Hi-Q reagents and P1 V3 chips and the sequencing was performed for 300 flows on the Ion Proton (ThermoFisher). After an initial run of sequencing (96-plex for the progeny and one chip for the parents/grandparents), all libraries were re-pooled in 96-plex and sequenced again to normalized read depth between individuals. Parents were sequenced at a greater depth than progeny to make an accurate catalogue of diploid genotypes possible in the cross. The father was very poorly sequenced, likely because of low-quality DNA, so the grand-parents were both re-sequenced at great depth to infer the father genotype when possible.

The library sequences for the 136 offspring and their parents were trimmed using cutadapt (-e 0.2, -m 50) and then split per sample using process_radtags (-c -r -t 80 -q -s 0 --barcode_dist_1 2 -E phred33 -e apekI). This resulted in an average of 1.23 million reads (stdev = 0.198 million reads) of 80bp per offspring, as well as 2940951 reads for the male and 5173632 reads for the female. The paternal grandparents had 4496703 and 4513732 for the male and female, respectively. These prepared reads were aligned on the scaffolded assembly with bwa mem (-k 19 -c 500 -O 0,0 -E 2,2 -T 0) and SAMtools (samtools view -Sb -q 1 -F 4 -F 256 -F 2048). Genotype likelihoods were obtained with SAMtools mpileup following the pipeline and parameters provided in lep-map3 documentation. Only markers with at least 3X of coverage in all individuals were kept. We explored more stringent filtering such as 6X and 10X, which led to very similar and collinear maps albeit with less marker density, an aspect that was the priority for efficient scaffolding.

### Supplementary methods 4: low-coverage whole-genome sequencing

- **Library preparation and sequencing**

DNA quality was evaluated with nanodrop and on a 1% agarose gel electrophoresis. Only samples with acceptable ratios that showed clear high molecular weight bands were retained for library preparation. Following (Therkildsen and Palumbi 2017), we remove DNA fragments shorter than 1kb by treating each extract with Axygen magnetic beads in a 0.4:1 ratio, and eluted the DNA in 10mM Tris-Cl, pH 8.5. We measured DNA concentrations with QuantiT Picogreen dsDNA Assay Kit (Invitrogen) and normalised all samples at a concentration of 5ng/µL. Then, sample DNA extracts were randomized, distributed in 17 plates (96-well) and re-normalised at 1ng/µL. Whole-genome high-quality libraries were prepared for each fly sample according to the protocol described in (Baym et al. 2015; Therkildsen and Palumbi 2017). Briefly, a tagmentation reaction using the enzyme from the Nextera kit, which simultaneously fragments the DNA and incorporates partial adapters, was carried out in a 2.5 µl volume with approximately 1 ng of input DNA. Then, we used a two-step PCR procedure with a total of 12 cycles (8+4) to add the remaining Illumina adapter sequence with dual index barcodes and amplify the libraries. The PCR was conducted with the KAPA Library Amplification Kit and custom primers derived from Nextera XT set of barcodes A,B,C and D (total 384 combinations). Amplification products were purified from primers and size-selected with a two-steps Axygen magnetic beads cleaning protocol, first with a ratio of 0.35:1, keeping the supernatant (medium and short DNA fragments), second with a ratio of 0.7:1, keeping the beads (medium fragments). The final concentrations of the libraries were quantified with QuantiT Picogreen dsDNA Assay Kit (Invitrogen) and fragment size distribution was estimated with an Agilent BioAnalyzer for a subset of 10 to 20 samples per plate. Equimolar amounts of 293 to 296 libraries were combined into 5 separate pools for sequencing on 5 lanes of paired-end 150bp reads on an Illumina HiSeq 4000 at the Norwegian Sequencing Center at the University of Oslo.

- **Sequence filtering and processing**

Raw reads were trimmed and filtered for quality with FastP (Chen et al. 2018). Reads were aligned to the reference genome with BWA-MEM (Li and Durbin 2009) and filtered with samtools v1.8 (Li et al. 2009) to keep only unpaired, orphaned, and concordantly paired reads with a mapping quality over 10. Duplicate reads were removed with the MarkDuplicates module of Picard Tools v1.119. Then, we realigned reads around indels with the GATK IndelRealigner (McKenna et al. 2010). Finally, to avoid double-counting the sequencing support during SNP calling, we used the clipOverlap program in the bamUtil package v1.0.14 (Breese and Liu 2013) to soft clip overlapping read ends and we kept only the read with the highest quality score in overlapping regions. This pipeline was inspired by (Therkildsen and Palumbi 2017) and is available at https://github.com/enormandeau/wgs_sample_preparation.

For most of the analysis, we used the program ANGSD v0.931 (Korneliussen et al. 2014), a software specifically designed to take genotype uncertainty into account instead of basing the analysis on called genotypes, which was appropriated for the low coverage of our data. The pipeline of analysis is available at https://github.com/clairemerot/angsd_pipeline. For all analysis, input reads were filtered to remove reads with a samtools flag above 255 (not primary, failure and duplicate reads, tag -remove_bads = 1), with mapping quality below 30 (-minMapQ = 30) and to remove bases with quality below 20 (-minQ 20). Note that to reduce the computational and analytic burden due to small scaffolds, all analyses were performed on a reduced genome including the 6 chromosomes and only 135 unanchored scaffolds, selected because they were longer than 25kb and bear more than 100 SNPs/scaffold. This reduced genome represents more than 89% of the total reference and more than 98.5% of all SNPs.

Abed A, Légaré G, Pomerleau S, St-Cyr J, Boyle B, Belzile FJ. 2019. Genotyping-by-sequencing on the ion torrent platform in barley. In: Barley. Springer. p. 233–252.

Aljanabi SM, Martinez I. 1997. Universal and rapid salt-extraction of high quality genomic DNA for PCR-based techniques. *Nucleic acids research* 25:4692–4693.

Baym M, Kryazhimskiy S, Lieberman TD, Chung H, Desai MM, Kishony R. 2015. Inexpensive multiplexed library preparation for megabase-sized genomes. *PloS one* 10:e0128036.

Breese MR, Liu Y. 2013. NGSUtils: a software suite for analyzing and manipulating next-generation sequencing datasets. *Bioinformatics* 29:494–496.

Cabanettes F, Klopp C. 2018. D-GENIES: dot plot large genomes in an interactive, efficient and simple way. *PeerJ* 6:e4958.

Catchen J, Amores A, Bassham S. 2020. Chromonomer: a tool set for repairing and enhancing assembled genomes through integration of genetic maps and conserved synteny. *bioRxiv*.

Chen S, Zhou Y, Chen Y, Gu J. 2018. fastp: an ultra-fast all-in-one FASTQ preprocessor. *Bioinformatics* 34:i884–i890.

Coombe L, Zhang J, Vandervalk BP, Chu J, Jackman SD, Birol I, Warren RL. 2018. ARKS: chromosome-scale scaffolding of human genome drafts with linked read kmers. *BMC bioinformatics* 19:234.

Elshire RJ, Glaubitz JC, Sun Q, Poland JA, Kawamoto K, Buckler ES, Mitchell SE. 2011. A Robust, Simple Genotyping-by-Sequencing (GBS) Approach for High Diversity Species. *PLOS ONE* 6:e19379.

Fu L, Niu B, Zhu Z, Wu S, Li W. 2012. CD-HIT: accelerated for clustering the next-generation sequencing data. *Bioinformatics* 28:3150–3152.

Grabherr MG, Haas BJ, Yassour M, Levin JZ, Thompson DA, Amit I, Adiconis X, Fan L, Raychowdhury R, Zeng Q. 2011. Full-length transcriptome assembly from RNA-Seq data without a reference genome. *Nature biotechnology* 29:644–652.

Haas BJ, Papanicolaou A, Yassour M, Grabherr M, Blood PD, Bowden J, Couger MB, Eccles D, Li B, Lieber M. 2013. De novo transcript sequence reconstruction from RNA-seq using the Trinity platform for reference generation and analysis. *Nature protocols* 8:1494–1512.

Jackman SD, Coombe L, Chu J, Warren RL, Vandervalk BP, Yeo S, Xue Z, Mohamadi H, Bohlmann J, Jones SJ. 2018. Tigmint: correcting assembly errors using linked reads from large molecules. *BMC bioinformatics* 19:1–10.

Korneliussen TS, Albrechtsen A, Nielsen R. 2014. ANGSD: analysis of next generation sequencing data. *BMC bioinformatics* 15:356.

Li H, Durbin R. 2009. Fast and accurate short read alignment with Burrows–Wheeler transform. *bioinformatics* 25:1754–1760.

Li H, Handsaker B, Wysoker A, Fennell T, Ruan J, Homer N, Marth G, Abecasis G, Durbin R. 2009. The sequence alignment/map format and SAMtools. *Bioinformatics* 25:2078–2079.

McKenna A, Hanna M, Banks E, Sivachenko A, Cibulskis K, Kernytsky A, Garimella K, Altshuler D, Gabriel S, Daly M. 2010. The Genome Analysis Toolkit: a MapReduce framework for analyzing next-generation DNA sequencing data. *Genome research* 20:1297–1303.

Mérot C, Berdan EL, Babin C, Normandeau E, Wellenreuther M, Bernatchez L. 2018. Intercontinental karyotype–environment parallelism supports a role for a chromosomal inversion in local adaptation in a seaweed fly. *Proc Biol Sci* [Internet] 285. Available from: http://rspb.royalsocietypublishing.org/content/285/1881/20180519.abstract

Simão FA, Waterhouse RM, Ioannidis P, Kriventseva EV, Zdobnov EM. 2015. BUSCO: assessing genome assembly and annotation completeness with single-copy orthologs. *Bioinformatics* 31:3210–3212.

Smit A, Hubley R, Green P. 2015. RepeatMasker Open-4.0. 2013–2015.

Smith-Unna R, Boursnell C, Patro R, Hibberd JM, Kelly S. 2016. TransRate: reference-free quality assessment of de novo transcriptome assemblies. *Genome research* 26:1134–1144.

Therkildsen NO, Palumbi SR. 2017. Practical low-coverage genomewide sequencing of hundreds of individually barcoded samples for population and evolutionary genomics in nonmodel species. *Molecular ecology resources* 17:194–208.

Walker BJ, Abeel T, Shea T, Priest M, Abouelliel A, Sakthikumar S, Cuomo CA, Zeng Q, Wortman J, Young SK. 2014. Pilon: an integrated tool for comprehensive microbial variant detection and genome assembly improvement. *PloS one* 9.

Wu TD, Watanabe CK. 2005. GMAP: a genomic mapping and alignment program for mRNA and EST sequences. *Bioinformatics* 21:1859–1875.

Yeo S, Coombe L, Warren RL, Chu J, Birol I. 2018. ARCS: scaffolding genome drafts with linked reads. *Bioinformatics* 34:725–731.

### Table S1: Environmental variables at sampled locations in North America

The “other seaweeds” category includes red, brown and green algae that did not belong to Fucaceae or Laminariceae. From Mérot *et al*, 2018

| | | GPS coordinates | | Climatic and abiotic variables extracted from databases | | | | | Wrackbed abiotic characteristics | | | Wrackbed algal composition (%) | | | | |
|---|---|---|---|---|---|---|---|---|---|---|---|---|---|---|---|---|
| | Location | Latitude (°) | Longitude (°) | Air T° (°C) | Precipitations (mm) | Sea T° (°C) | Sea Salinity (‰) | Tidal Amplitude (m) | Mean depth (m) | Mean T° (°C) | Salinity | Fucaceae | Laminariaceae | Plant Debris | Other Seaweeds | Zoosteraceae |
| **AG** | Anse du Griffon (QC, C.) | 48.93491 | -64.30589 | 2.7 | 104.6 | 5.1 | 28.4 | 1.36 | 0.55 | 18.7 | 154 | 5 | 90 | 0 | 5 | 0 |
| **BP** | Black Point (ME, USA) | 43.53059 | -70.32209 | 8.1 | 113.9 | 8.8 | 31.5 | 3.06 | 0.2 | 16.2 | 8 | 25 | 5 | 0 | 20 | 50 |
| **BS** | Blanc Sablon (QC, C.) | 51.41545 | -57.15290 | 0.8 | 112.6 | 3.6 | 31.0 | 1.40 | 0.4 | 10.0 | | 68 | 30 | 0 | 2 | 0 |
| **BT** | Baie Trinité (QC, C) | 49.41716 | -67.30285 | 1.4 | 97.3 | 5.4 | 28.8 | 2.89 | 0.75 | 40.9 | 164 | 98 | 2 | 0 | 0 | 0 |
| **CB** | Cow Bay (NS, C.) | 44.62190 | -63.42112 | 6.4 | 140.9 | 7.1 | 30.3 | 1.46 | 0.35 | 17.1 | 10 | 5 | 70 | 0 | 25 | 0 |
| **CE** | Cap Espoir (QC) | 48.43087 | -64.32778 | 3.4 | 109.6 | 6.1 | 29.8 | 1.12 | 0.15 | 18.8 | 114 | 3 | 95 | 5 | 2 | 0 |
| **GM** | Grands Méchins (QC, C) | 49.00427 | -66.97155 | 2.8 | 96.2 | 5.1 | 28.9 | 2.44 | 0.3 | 11.9 | 44 | 40 | 40 | 20 | 0 | 0 |
| **HA** | Hampton (NH, USA) | 42.92098 | -70.79826 | 8.7 | 114.4 | 9.3 | 31.4 | 2.87 | 0.45 | 37.3 | 179 | 50 | 2 | 0 | 50 | 0 |
| **KA** | Kamouraska (QC, C.) | 47.56294 | -69.87375 | 3.9 | 94.4 | 5.4 | 15.5 | 4.72 | 0.35 | 17.3 | 19 | 75 | 5 | 20 | 0 | 0 |
| **MA** | Manomet Point (MA, USA) | 41.92654 | -70.54451 | 9.8 | 119.9 | 10.0 | 31.8 | 3.13 | 0.15 | 12.0 | 9 | 90 | 2 | 0 | 2 | 5 |
| **ME** | Métis (QC, C) | 48.66408 | -68.07221 | 2.4 | 93.0 | 4.4 | 24.6 | 3.06 | 0.3 | 19.4 | 95 | 60 | 0 | 14 | 1 | 25 |
| **NB** | Naufrage Beach (PEI, C.) | 46.46795 | -62.41561 | 5.7 | 109.2 | 8.3 | 30.0 | 0.73 | 0.65 | 11.1 | 14 | 70 | 2 | 0 | 1 | 0 |
| **RB** | Rivière du Bouleau (QC) | 50.28161 | -65.51516 | 1.3 | 98.9 | 4.9 | 30.2 | 1.53 | 0.2 | 17.0 | 80 | 80 | 20 | 0 | 0 | 0 |
| **RC** | Rivière à Claude (QC) | 49.22086 | -65.89794 | 2.3 | 98.2 | 5.1 | 29.4 | 2.40 | 0.5 | 20.7 | 53 | 45 | 5 | 50 | 0 | 0 |
| **SI** | Saint Irénée (QC) | 47.55973 | -70.20425 | 2.7 | 105.1 | 5.4 | 15.5 | 4.72 | 0.15 | 6.8 | 0 | 80 | 15 | 0 | 5 | 0 |
| **SS** | Saint Siméon (QC) | 48.06991 | -65.56586 | 4.2 | 103.3 | 7.0 | 29.1 | 1.66 | 0.2 | 14.9 | 113 | 20 | 50 | 0 | 5 | 25 |

### Table S2: Concordance with SNP marker

Individuals genotyped with the PCA on the whole-genome data for the inversion *Cf-Inv(1)* were all consistent with the genotype previously obtained in the lab (Mérot et al. 2018), except for the ones listed below. They were re-verified in the lab with an independent PCR and corrected for 2 of them (BS16-0088 and SI16-0021). For all the others, PCA genotyping was concordant with genotyping based on the enzymatic test with AluI but not DraI.

| | | PCA genotype | Initial genotype with AluI | Initial genotype with DraI | Verification of genotype with AluI | Verification of genotype with DraI |
|---|---|---|---|---|---|---|
| North | BS16-0088 | αβ | αβ | ββ | αβ | αβ |
| Gaspesie | SI16-0021 | αβ | αα | NA | αβ | αβ |
| | GM16-0056 | αβ | αβ | ββ | αβ | ββ |
| | KA16-0037 | αβ | αβ | ββ | αβ | ββ |
| USA | BP16-0035 | αβ | αβ | ββ | αβ | ββ |
| | BP16-0042 | αβ | αβ | ββ | αβ | ββ |
| | BP16-0071 | αβ | αβ | ββ | αβ | ββ |
| | BP16-0080 | αβ | αβ | ββ | αβ | ββ |
| | BP16-0090 | αβ | αβ | ββ | αβ | ββ |
| | HA16-0009 | αβ | αβ | ββ | αβ | ββ |
| | HA16-0034 | αβ | αβ | ββ | αβ | ββ |
| | HA16-0084 | αβ | αβ | ββ | αβ | ββ |
| | HA16-0090 | αβ | αβ | ββ | αβ | ββ |
| | HA16-0101 | αβ | αβ | ββ | αβ | ββ |
| | MA16-0006 | αβ | αβ | ββ | αβ | ββ |
| | MA16-0032 | αβ | αβ | ββ | αβ | ββ |
| | MA16-0037 | αβ | αβ | ββ | αβ | ββ |
| | MA16-0055 | αβ | αβ | ββ | αβ | ββ |
| | MA16-0057 | αβ | αβ | ββ | αβ | ββ |
| | MA16-0073 | αβ | αβ | ββ | αβ | ββ |
| | MA16-0083 | αβ | αβ | ββ | αβ | ββ |
| | MA16-0088 | αβ | αβ | ββ | αβ | ββ |
| | MA16-0090 | αβ | αβ | ββ | αβ | ββ |
| | MA16-0096 | αβ | αβ | ββ | αβ | ββ |
| | MA16-0100 | αβ | αβ | ββ | αβ | ββ |
| | MA16-0107 | αβ | αβ | ββ | αβ | ββ |

### Table S3: Isolation-by-distance / Isolation-by-resistance

Linear models testing the association between genetic distance (calculated on LD-pruned SNPs) and geographic distances measured as Euclidian distances or least-cost distances along the shoreline.

| Model | F | p-value | intercept | slope coefficient | $R^2$ adjusted | AIC |
|---|---|---|---|---|---|---|
| Null model | | | | | | -1078 |
| Euclidian distances | 96.8 | <0.001 | 0.006 | 0.0018 [0.0014-0.0022] | 0.45 | -1148 |
| Least-cost distances | 199.5 | <0.001 | 0.006 | 0.0021 [0.0018-0.0024] | 0.63 | -1195 |

### Table S4: Isolation-by-distance within the different genomic regions

Linear models testing the association between genetic distance and geographic distances measured as Euclidian distances in the different subsets of SNPs. Numbers between brackets indicate the limits of the 95% distribution of the slope coefficient.

| SNP subset | F | p-value | intercept | slope coefficient | $R^2$ adjusted | mean pairwise Fst |
|---|---|---|---|---|---|---|
| All | 16.3 | <0.001 | 0.0085 | 0.0015 [0.0008-0.0023] | 0.11 | 0.0084 |
| Collinear | 62.6 | <0.001 | 0.0062 | 0.0015 [0.0011-0.0019] | 0.34 | 0.0062 |
| LD pruned | 96.8 | <0.001 | 0.0057 | 0.0018 [0.0014-0.0022] | 0.45 | 0.0057 |
| *Cf-Inv(1)* | 0.8 | 0.37 | 0.0137 | -0.0011 [-0.0036-0.0014] | 0.00 | 0.0133 |
| *Cf-Inv(4.1)* | 39.6 | <0.001 | 0.0172 | 0.0124 [0.0085-0.0162] | 0.24 | 0.0164 |
| *Cf-Inv(4.2/4.3)* | 41.6 | <0.001 | 0.0075 | 0.0022 [0.0015-0.0028] | 0.25 | 0.0074 |
| *Cf-Lrr(2)* | 74.6 | <0.001 | 0.0074 | 0.0026 [0.0020-0.0032] | 0.38 | 0.0073 |
| *Cf-Lrr(3)* | 57.5 | <0.001 | 0.0066 | 0.0016 [0.0012-0.0020] | 0.32 | 0.0066 |
| *Cf-Lrr(5)* | 82.4 | <0.001 | 0.0080 | 0.0028 [0.0022-0.0035] | 0.41 | 0.0079 |

### Table S5: Enrichment in outlier SNPs associated with environment for the different GEA methods

**Baypass uncontrolled**

| | Tested SNPs | | Climate | | | Salinity | | | Bed abiotic characteristics | | | Algal composition (Laminaria/Fucus) | | | Algal composition (PC2) | | |
|---|---|---|---|---|---|---|---|---|---|---|---|---|---|---|---|---|---|
| | N | % | N | % | OR | N | % | OR | N | % | OR | N | % | OR | N | % | OR |
| All | 1155978 | | 4369 | | | 1584 | | | 499 | | | 1747 | | | 2960 | | |
| Collinear | 814279 | 70% | 913 | 21% | 0.3 | 1025 | 65% | 0.9 | 321 | 64% | 0.9 | 1303 | 75% | 1.1 | 657 | 22% | 0.3 |
| *Cf-Inv(1)* | 176963 | 15% | **1298** | **30%** | **1.9*** | 112 | 7.1% | 0.5 | **106** | **21%** | **1.4*** | 192 | 11% | 0.7 | **988** | **33%** | **2.2*** |
| *Cf-Inv(4.1)* | 57323 | 5.0% | **362** | **8.3%** | **1.7*** | 44 | 2.8% | 0.6 | 22 | 4.4% | 0.9 | 75 | 4.3% | 0.9 | 35 | 1.2% | 0.2 |
| *Cf-Inv(4.2/4.3)* | 17019 | 1.5% | **168** | **3.8%** | **2.6*** | 22 | 1.4% | 0.9 | 12 | 2.4% | 1.6 | 25 | 1.4% | 1.0 | 20 | 0.7% | 0.5 |
| *Cf-Lrr(2)* | 20458 | 1.8% | **232** | **5.3%** | **3.0*** | 31 | 2.0% | 1.1 | **17** | **3.4%** | **1.9*** | 22 | 1.3% | 0.7 | 34 | 1.1% | 0.6 |
| *Cf-Lrr(3)* | 16313 | 1.4% | 18 | 0.4% | 0.3 | **97** | **6.1%** | **4.3*** | 3 | 0.6% | 0.4 | 32 | 1.8% | 1.3 | 15 | 0.5% | 0.4 |
| *Cf-Lrr(5)* | 53623 | 4.6% | **1378** | **32%** | **6.8*** | **253** | **16%** | **3.4*** | 18 | 3.6% | 0.8 | 98 | 5.6% | 1.2 | **1211** | **41%** | **8.8*** |

**Baypass controlled**

| | Tested SNPs | | Climate | | | Salinity | | | Bed abiotic characteristics | | | Algal composition (Laminaria/Fucus) | | | Algal composition (PC2) | | |
|---|---|---|---|---|---|---|---|---|---|---|---|---|---|---|---|---|---|
| | N | % | N | % | OR | N | % | OR | N | % | OR | N | % | OR | N | % | OR |
| All | 1155978 | | 7230 | | | 1670 | | | 8765 | | | 1855 | | | 10713 | | |
| Collinear | 814279 | 70% | 1223 | 17% | 0.2 | 986 | 59% | 0.8 | 819 | 9.3% | 0.1 | 1231 | 66% | 0.9 | 1747 | 16% | 0.2 |
| *Cf-Inv(1)* | 176963 | 15% | **2521** | **35%** | **2.3*** | 237 | 14% | 0.9 | **7772** | **89%** | **5.8*** | **421** | **23%** | **1.5*** | **6594** | **62%** | **4.0*** |
| *Cf-Inv(4.1)* | 57323 | 5.0% | **1195** | **17%** | **3.5*** | 44 | 2.6% | 0.5 | 38 | 0.4% | 0.1 | 54 | 2.9% | 0.6 | 108 | 1.0% | 0.2 |
| *Cf-Inv(4.2/4.3)* | 17019 | 1.5% | **180** | **2.5%** | **1.7*** | 20 | 1.2% | 0.8 | 36 | 0.4% | 0.3 | 26 | 1.4% | 1.0 | 70 | 0.7% | 0.4 |
| *Cf-Lrr(2)* | 20458 | 1.8% | **298** | **4.1%** | **2.3*** | 39 | 2.3% | 1.3 | 43 | 0.5% | 0.3 | 17 | 0.9% | 0.5 | 84 | 0.8% | 0.4 |
| *Cf-Lrr(3)* | 16313 | 1.4% | 23 | 0.3% | 0.2 | **99** | **5.9%** | **4.2*** | 6 | 0.1% | 0.0 | 21 | 1.1% | 0.8 | 50 | 0.5% | 0.3 |
| *Cf-Lrr(5)* | 53623 | 4.6% | **1790** | **25%** | **5.3*** | **245** | **15%** | **3.2*** | 51 | 0.6% | 0.1 | 85 | 4.6% | 1.0 | **2060** | **19%** | **4.1*** |

**Lfmm k=4**

| | Tested SNPs | | Climate | | | Salinity | | | Bed abiotic characteristics | | | Algal composition (Laminaria/Fucus) | | | Algal composition (PC2) | | |
|---|---|---|---|---|---|---|---|---|---|---|---|---|---|---|---|---|---|
| | N | % | N | % | OR | N | % | OR | N | % | OR | N | % | OR | N | % | OR |
| All | 1155978 | | 9712 | | | 3520 | | | 3092 | | | 2748 | | | 6240 | | |
| Collinear | 814279 | 70% | 3571 | 37% | 0.5 | 2355 | 67% | 0.9 | 1810 | 59% | 0.8 | 1860 | 68% | 1.0 | 2534 | 41 % | 0.6 |
| *Cf-Inv(1)* | 176963 | 15% | **2428** | **25%** | **1.6*** | 483 | 14% | 0.9 | **890** | **29%** | **1.9*** | **522** | **19 %** | **1.2*** | **2103** | **34%** | **2.2*** |
| *Cf-Inv(4.1)* | 57323 | 5.0% | **1370** | **14%** | **2.8*** | 147 | 4.2% | 0.8 | 134 | 4.3% | 0.9 | 118 | 4.3% | 0.9 | 225 | 3.6% | 0.7 |
| *Cf-Inv(4.2/4.3)* | 17019 | 1.5% | **515** | **5.3%** | **3.6*** | 56 | 1.6% | 1.1 | 55 | 1.8% | 1.2 | 39 | 1.4% | 1.0 | 71 | 1.1% | 0.8 |
| *Cf-Lrr(2)* | 20458 | 1.8% | **228** | **2.3%** | **1.3*** | 54 | 1.5% | 0.9 | 45 | 1.5% | 0.8 | 32 | 1.2% | 0.7 | 55 | 0.9% | 0.5 |
| *Cf-Lrr(3)* | 16313 | 1.4% | 76 | 0.8% | 0.6 | **113** | **3.2%** | **2.3*** | 30 | 1.0% | 0.7 | 23 | 0.8% | 0.6 | 62 | 1.0% | 0.7 |
| *Cf-Lrr(5)* | 53623 | 4.6% | **1524** | **16%** | **3.4*** | **312** | **8.9%** | **1.9*** | **128** | **4.1%** | **0.9** | **154** | **5.6%** | **1.2*** | **1190** | **19%** | **4.1*** |

### Table S6: Gene ontology enrichment for SNPs associated with size by GWAS

| id | level | name | Outliers SNPs | All SNPs | p value | fdr |
|---|---|---|---|---|---|---|
| GO:0048853 | 3 | forebrain morphogenesis | 6/2412 | 7/17848 | 3.75E-05 | 0.030 |
| GO:0030259 | 3 | lipid glycosylation | 6/2412 | 7/17848 | 3.75E-05 | 0.030 |
| GO:0021764 | 3 | amygdala development | 5/2412 | 6/17848 | 0.000239 | 0.079 |
| GO:0016477 | 3 | cell migration | 13/2412 | 258/17848 | 1.29E-05 | 0.024 |
| GO:0007018 | 3 | microtubule-based movement | 9/2412 | 188/17848 | 0.000144 | 0.079 |
| GO:0035148 | 3 | tube formation | 0/2412 | 58/17848 | 0.000356 | 0.085 |
| GO:0048485 | 4 | sympathetic nervous system development | 5/2412 | 5/17848 | 4.49E-05 | 0.030 |
| GO:0001649 | 4 | osteoblast differentiation | 9/2412 | 18/17848 | 0.000227 | 0.079 |
| GO:0098597 | 4 | observational learning | 5/2412 | 6/17848 | 0.000239 | 0.079 |
| GO:0048820 | 4 | hair follicle maturation | 5/2412 | 6/17848 | 0.000239 | 0.079 |
| GO:0051969 | 4 | regulation of transmission of nerve impulse | 4/2412 | 4/17848 | 0.000333 | 0.085 |
| GO:1902644 | 4 | tertiary alcohol metabolic process | 4/2412 | 4/17848 | 0.000333 | 0.085 |
| GO:0033561 | 4 | regulation of water loss via skin | 7/2412 | 12/17848 | 0.000346 | 0.085 |
| GO:1900271 | 4 | regulation of long-term synaptic potentiation | 6/2412 | 9/17848 | 0.000352 | 0.085 |
| GO:0015074 | 5 | DNA integration | 47/2412 | 154/17848 | 1.15E-06 | 0.009 |
| GO:0051348 | 5 | negative regulation of transferase activity | 18/2412 | 54/17848 | 0.000161 | 0.079 |
| GO:0048745 | 5 | smooth muscle tissue development | 5/2412 | 6/17848 | 0.000239 | 0.079 |
| GO:0014044 | 5 | Schwann cell development | 5/2412 | 6/17848 | 0.000239 | 0.079 |
| GO:0061535 | 5 | glutamate secretion, neurotransmission | 5/2412 | 6/17848 | 0.000239 | 0.079 |
| GO:0061534 | 5 | gamma-aminobutyric acid secretion, neurotransmission | 5/2412 | 6/17848 | 0.000239 | 0.079 |
| GO:0045761 | 5 | regulation of adenylate cyclase activity | 10/2412 | 22/17848 | 0.000273 | 0.085 |
| GO:0051971 | 5 | positive regulation of transmission of nerve impulse | 4/2412 | 4/17848 | 0.000333 | 0.085 |
| GO:0045765 | 5 | regulation of angiogenesis | 12/2412 | 31/17848 | 0.00042 | 0.099 |

### Table S7: Gene ontology enrichment for genes in the inversion *Cf-Inv(1)*

| id | level | name | Inversion genes | All genes | p value | fdr |
|---|---|---|---|---|---|---|
| GO:0030259 | 3 | lipid glycosylation | 6/2629 | 7/18808 | 4.57E-05 | 0.036 |
| GO:0009886 | 3 | post-embryonic animal morphogenesis | 5/2629 | 174/18808 | 2.47E-06 | 0.008 |
| GO:0016477 | 3 | cell migration | 14/2629 | 264/18808 | 9.32E-06 | 0.015 |
| GO:0048485 | 4 | sympathetic nervous system development | 5/2629 | 5/18808 | 5.32E-05 | 0.036 |
| GO:0035120 | 4 | post-embryonic appendage morphogenesis | 0/2629 | 73/18808 | 2.63E-05 | 0.036 |
| GO:0015074 | 5 | DNA integration | 61/2629 | 175/18808 | 1.47E-06 | 0.008 |
| GO:0030073 | 5 | insulin secretion | 6/2629 | 7/18808 | 4.57E-05 | 0.036 |

### Table S8: Gene ontology enrichment for genes in the inversion *Cf-Inv(4.1)*

| id | Lev. | name | Inversion genes | All genes | p value | fdr |
|---|---|---|---|---|---|---|
| GO:0007613 | 3 | memory | 12/686 | 80/18808 | 3.13E-05 | 0.024 |
| GO:0006810 | 3 | transport | 89/686 | 1622/18808 | 9.64E-05 | 0.045 |
| GO:0008344 | 3 | adult locomotory behavior | 10/686 | 65/18808 | 0.000114 | 0.045 |
| GO:0031570 | 3 | DNA integrity checkpoint | 7/686 | 33/18808 | 0.000155 | 0.045 |
| GO:0021532 | 4 | neural tube patterning | 4/686 | 4/18808 | 1.75E-06 | 0.009 |
| GO:0008610 | 4 | lipid biosynthetic process | 24/686 | 256/18808 | 2.39E-05 | 0.024 |
| GO:0035845 | 4 | photoreceptor cell outer segment organization | 4/686 | 7/18808 | 5.62E-05 | 0.035 |
| GO:0000076 | 4 | DNA replication checkpoint | 5/686 | 14/18808 | 9.68E-05 | 0.045 |
| GO:0048568 | 4 | embryonic organ development | 7/686 | 33/18808 | 0.000155 | 0.045 |
| GO:0009111 | 4 | vitamin catabolic process | 3/686 | 4/18808 | 0.000188 | 0.045 |
| GO:0043320 | 4 | natural killer cell degranulation | 3/686 | 4/18808 | 0.000188 | 0.045 |
| GO:0006873 | 4 | cellular ion homeostasis | 17/686 | 172/18808 | 0.000191 | 0.045 |
| GO:0007628 | 4 | adult walking behavior | 5/686 | 16/18808 | 0.000199 | 0.045 |
| GO:0048678 | 4 | response to axon injury | 5/686 | 16/18808 | 0.000199 | 0.045 |
| GO:0000038 | 5 | very long-chain fatty acid metabolic process | 8/686 | 22/18808 | 6.11E-07 | 0.007 |
| GO:0031076 | 5 | embryonic camera-type eye development | 4/686 | 5/18808 | 8.52E-06 | 0.012 |
| GO:1990403 | 5 | embryonic brain development | 4/686 | 6/18808 | 2.48E-05 | 0.024 |
| GO:0033559 | 5 | unsaturated fatty acid metabolic process | 7/686 | 33/18808 | 0.000155 | 0.045 |
| GO:0042365 | 5 | water-soluble vitamin catabolic process | 3/686 | 4/18808 | 0.000188 | 0.045 |
| GO:0002323 | 5 | natural killer cell activation involved in immune response | 3/686 | 4/18808 | 0.000188 | 0.045 |
| GO:0016036 | 5 | cellular response to phosphate starvation | 3/686 | 4/18808 | 0.000188 | 0.045 |

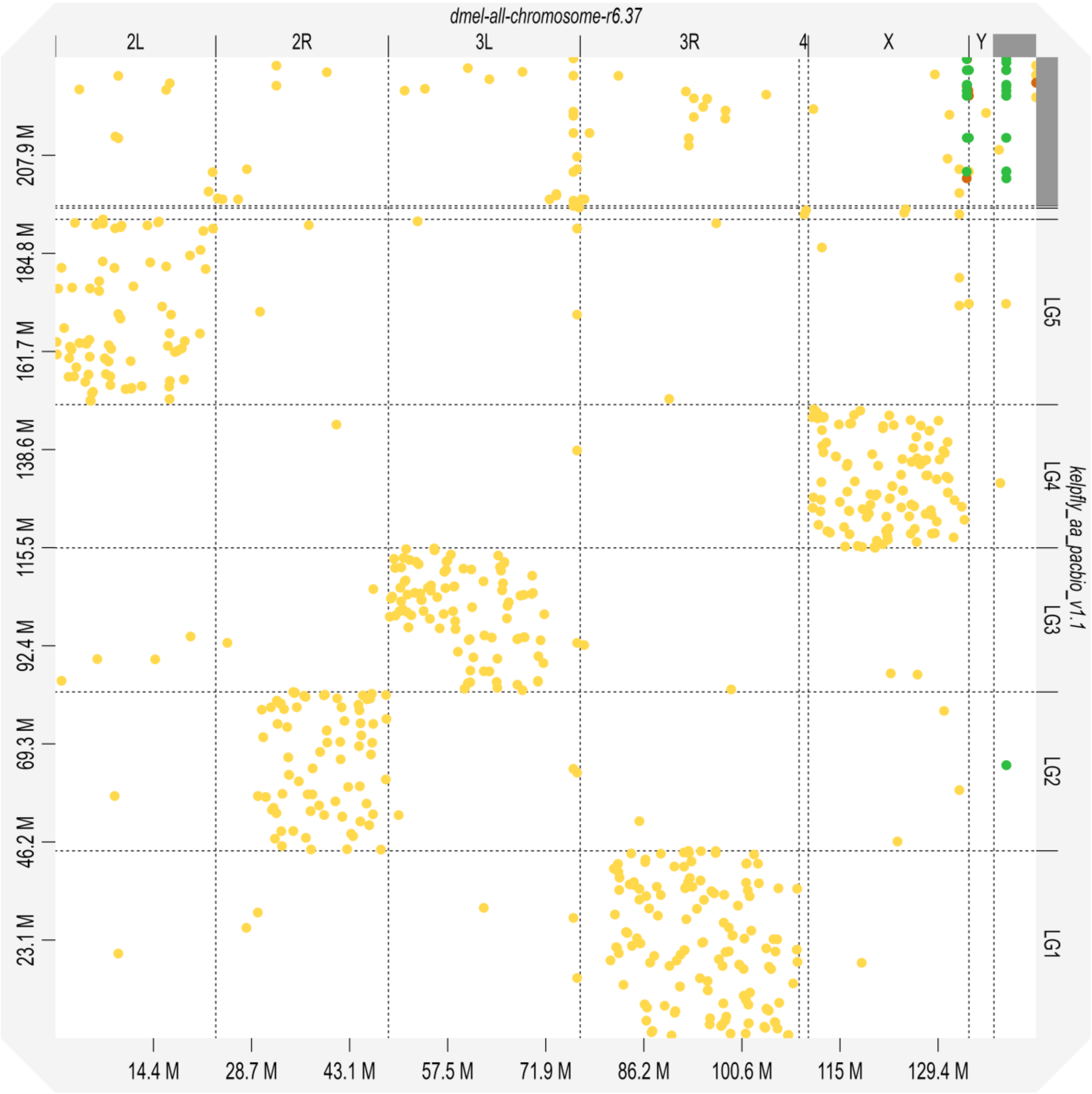

### Fig. S1: Dot plot comparing the genome of *Coelopa frigida* to *Drosophila melanogaster*

Alignment between the genome of *Coelopa frigida* and the reference genome of *Drosophila melanogaster r6.37* (Flybase) using minimap 2 and visualized with D-GENIES (Cabanettes and Klopp 2018). Colours represent the level of identity from 0 to 0.25 in yellow, 0.25 to 0.5 in orange, and above 0.5 in green.

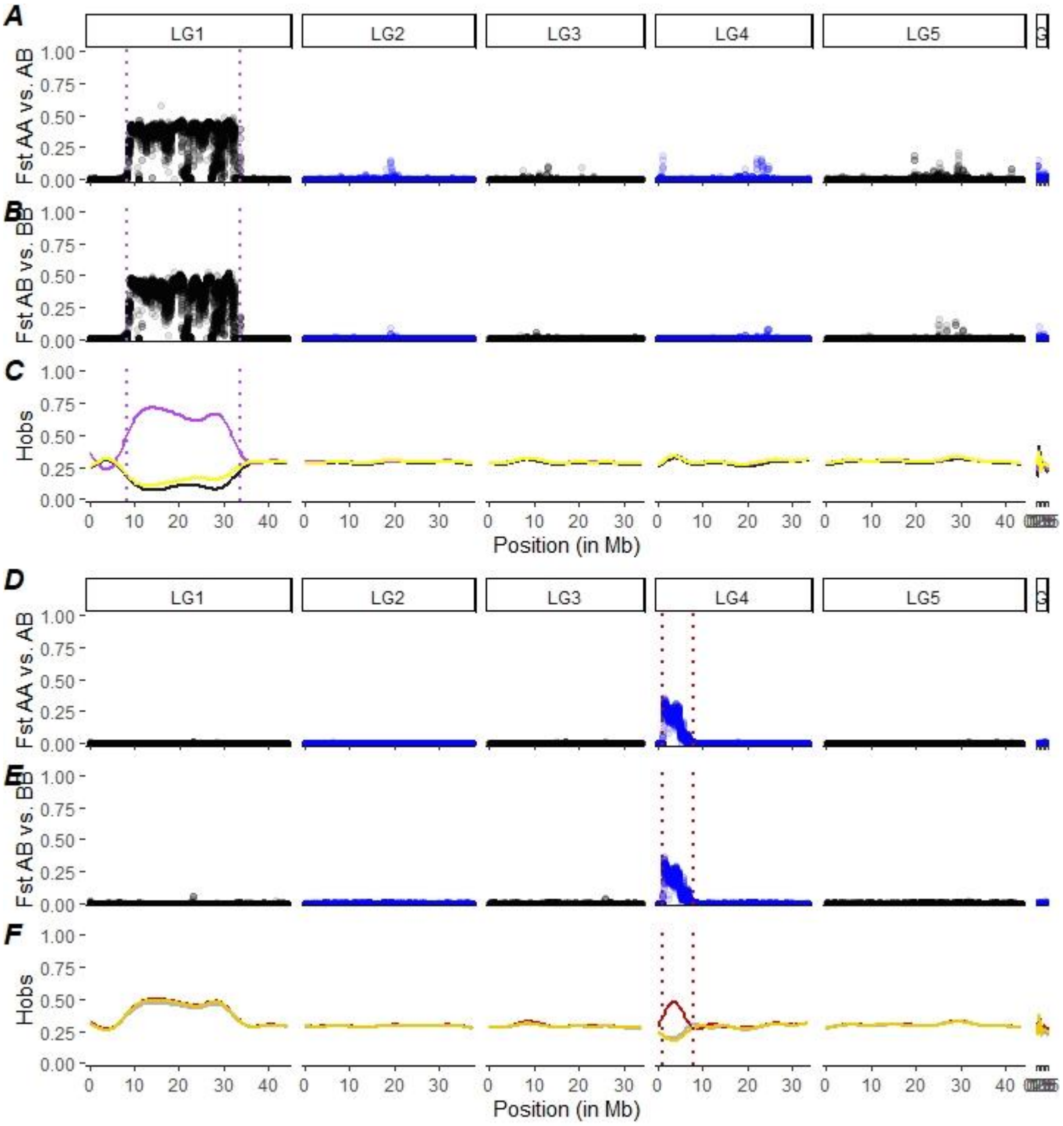

### Fig. S2: Differentiation and observed number of heterozygotes in haplogroups

(A-B) $F_{ST}$ between homokaryotypes and heterokaryotypes for the inversion *Cf-In(1)* (AA stands for αα, AB for αβ, and BB for ββ). (C) Observed proportion of heterozygotes within each haplogroup of the inversion *Cf-Inv(1)* (D-E) $F_{ST}$ between homokaryotypes and heterokaryotypes for the inversion *Cf-Inv(4.1)* (AA and BB are homokaryotypes at that inversion and AB is the heterokaryotype). (F) Observed proportion of heterozygotes within each haplogroup of the inversion *Cf-Inv(1)*. $F_{ST}$ is calculated by sliding-windows of 25kb, and the observed proportion of heterozygotes is smoothed for visualization

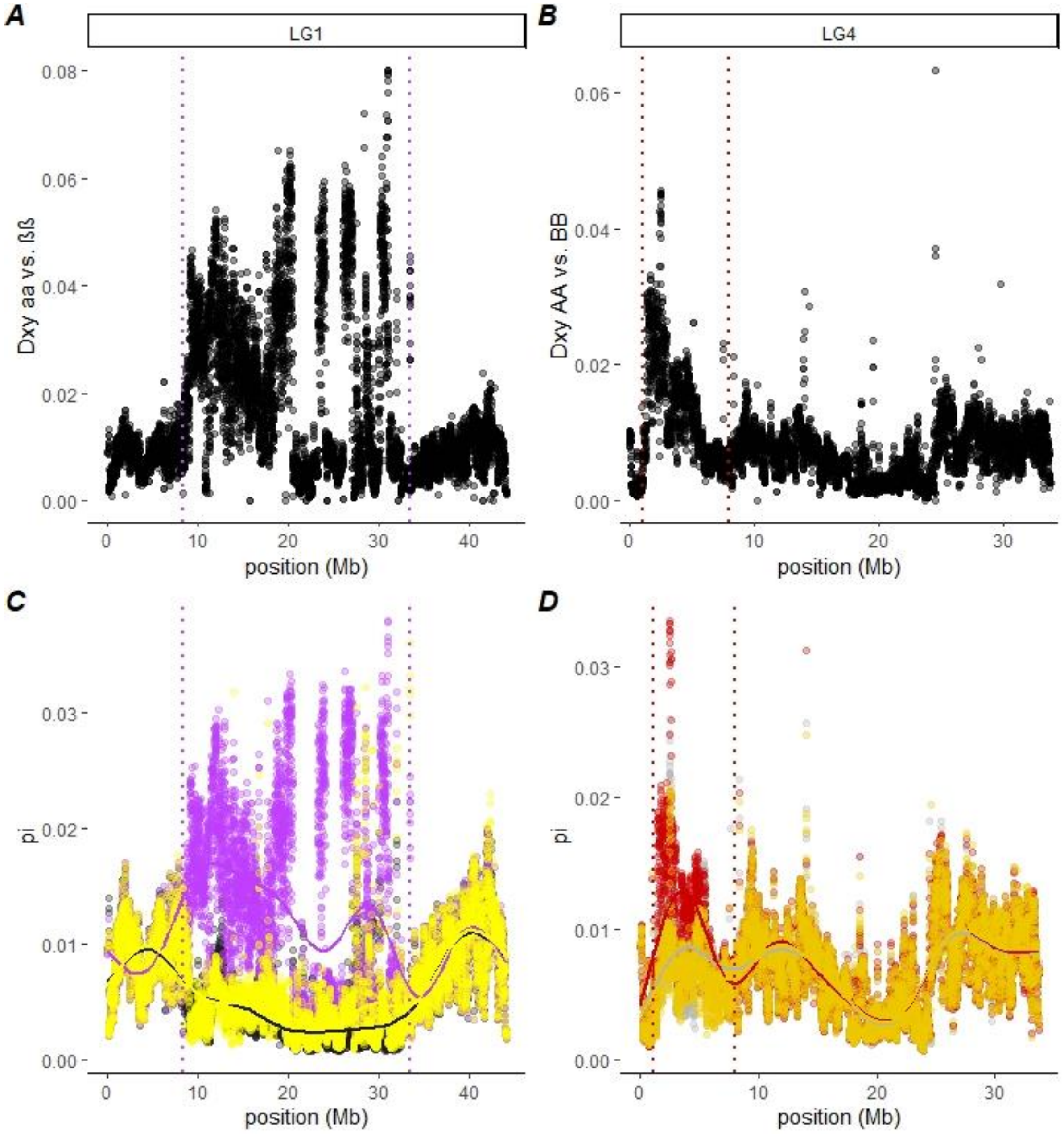

### Fig. S3: Nucleotide diversity and divergence in inversions

(A) Absolute nucleotide divergence ($d_{XY}$) between αα and ββ homokaryotypes for the inversion *Cf-Inv(1).* (B) Absolute nucleotide divergence ($d_{XY}$) between AA and BB homokaryotypes for the inversion *Cf-Inv(4.1).* (C) Nucleotide diversity (π) in each karyotypic group (black = αα, purple = αβ, yellow= ββ) of the inversion *Cf-Inv(1).* (D) Nucleotide diversity (π) in each karyotypic group (grey = AA, red= AB, gold=BB) of the inversion *Cf-Inv(4.1).* $d_{XY}$ and π are calculated by windows of 25kb.

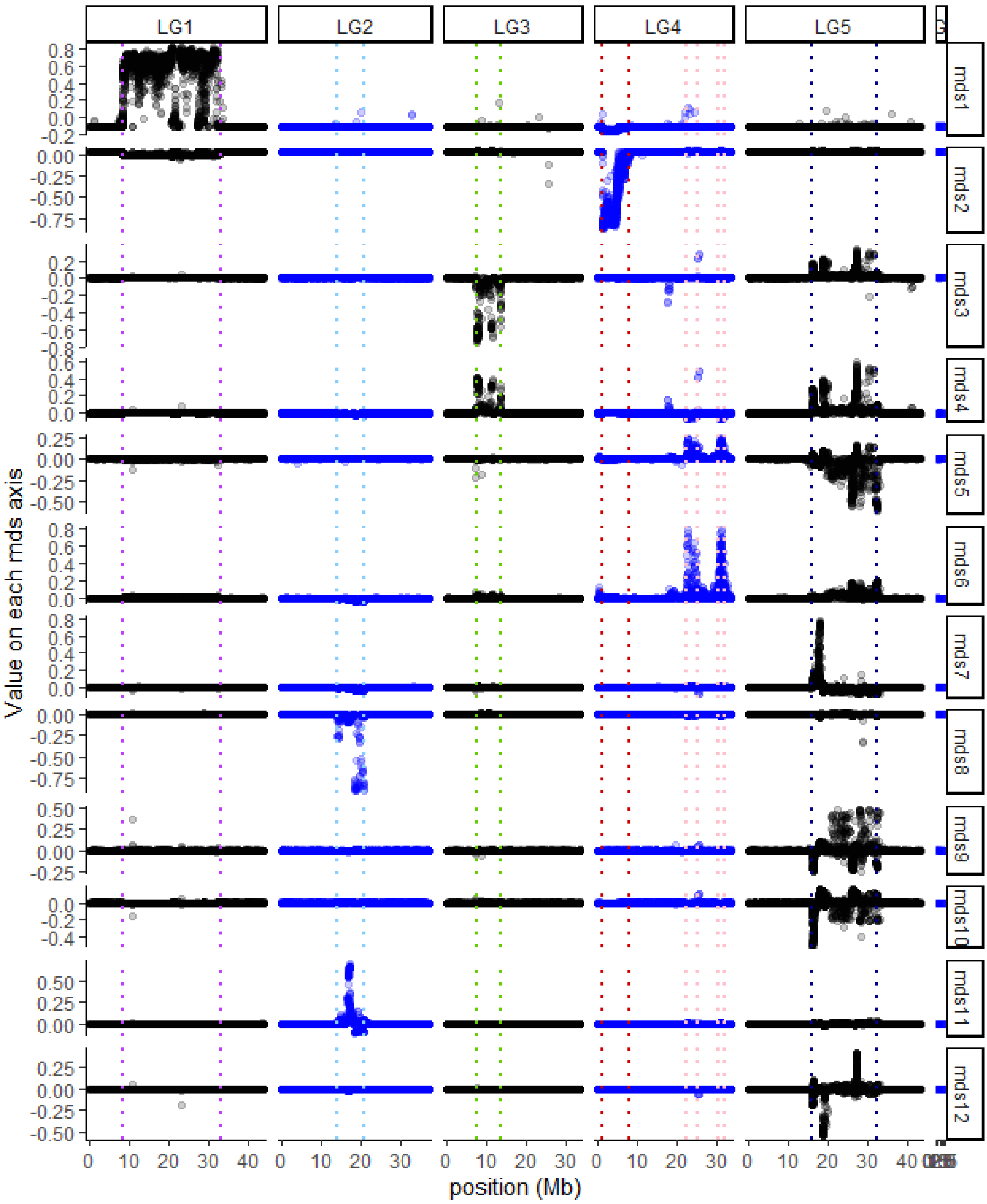

### Fig. S4: Multidimensional scaling of local PCAs.

For each MDS axis (up to 12), the Y-axis represents the MDS value of each local PCA matrix (based on windows of 100 SNPs) and the x-axis is the position along the chromosome. Dotted coloured lines denote the boundaries of the inversions and low-recombining regions.

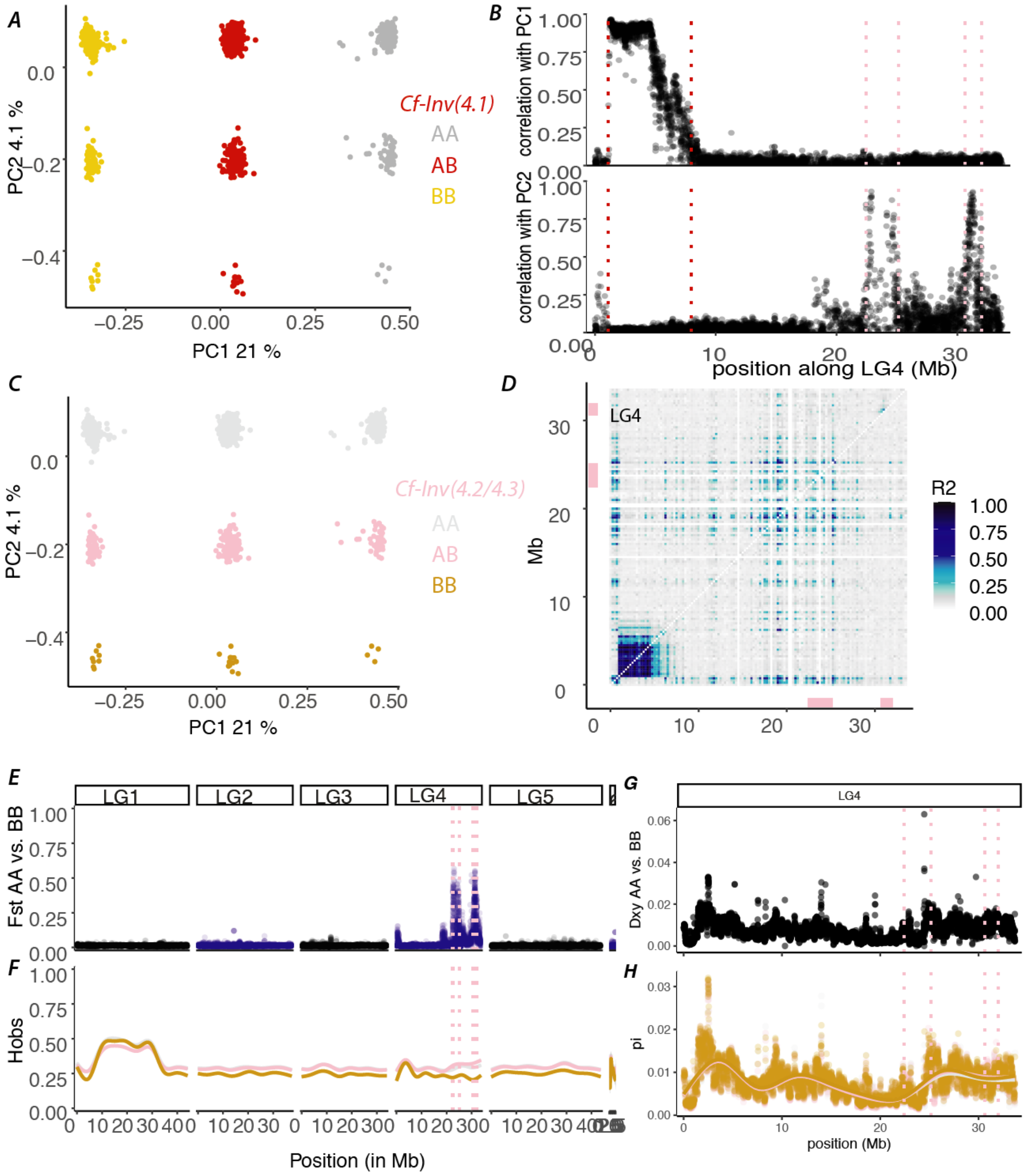

### Fig. S5: The inversion(s) *Cf-Inv(4.2)* and *Cf-Inv(4.3)* on LG4

**(A-C)** Principal component analysis (PCA) of genetic variation in LG4. Individuals are coloured by karyotypes at the inversions *Cf-Inv(4.1)* and *Cf-Inv(4.2/4.3)*. AA and BB stand for the homokaryotypes and AB for the heterokaryotypes, for each inversion. **(B)** Correlation between PC1 scores of local PCAs performed on windows of 100SNPs and PC1/PC2 scores of the PCA performed SNPs from LG4. Dashed lines represent the inferred boundaries of the inversions *Cf-Inv(4.1)* and *Cf-Inv(4.2/4.3)* **(D)** Linkage disequilibrium (LD) in LG4. The colour scale shows the $2^{nd}$ higher percentile of the $R^2$ value between SNPs summarized by windows of 250kb. The upper triangle includes all individuals and the lower triangle include individuals homokaryotypes for the most common arrangement at inversion *Cf-Inv(4.2/4.3)*. Bars represent the position of the inversion(s) *Cf-Inv(4.2/4.3)*. **(E)** FST differentiation between the two homokaryotypes of *Cf-Inv(4.2/4.3)* in sliding-windows of 25kb. **(F)** Observed proportion of heterozygotes at each SNP in the three karyotypic groups of *Cf-Inv(4.2/4.3)* smoothed for visualization. **(G)** Absolute nucleotide divergence ($d_{XY}$) between homokaryotypes of the inversions *Cf-Inv(4.2/4.3)* in sliding-windows of 25kb on LG4 **(H)** Nucleotide diversity (π) in each karyotypic group of the inversions *Cf-Inv(4.2/4.3)* in sliding-windows of 25kb on LG4.

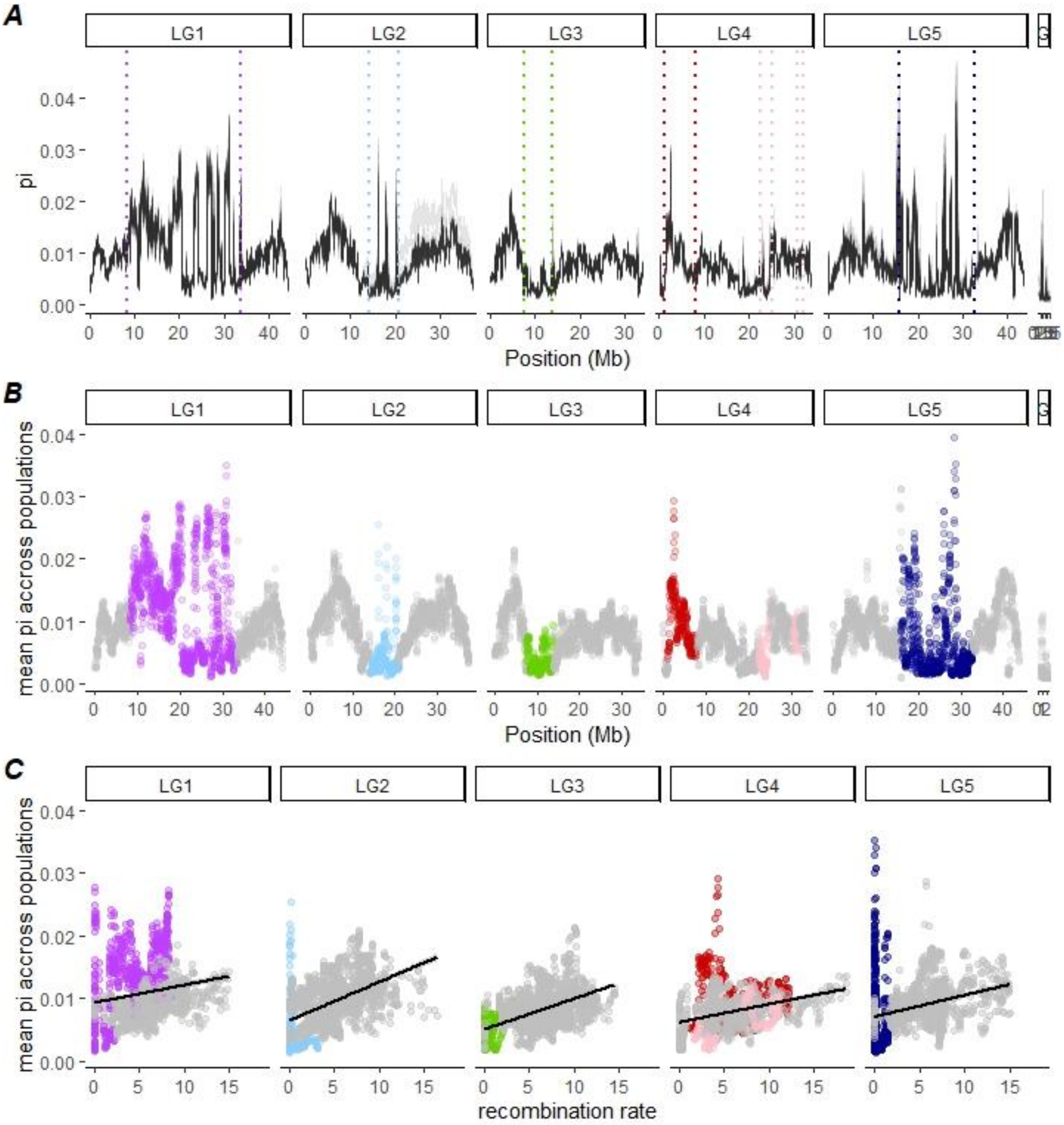

### Fig. S6: Genetic diversity along the genome and as a function of recombination

(A) Genetic diversity (π) along the genome for each population. Each line is a population. The lines (plotted with transparency) overlap showing that the diversity landscape followed the same pattern across populations, except on LG2, in which USA populations had a slightly higher pi. In the lower panels, π is average across the 16 populations and displayed along the genome (B), and then, as a function of recombination rate by sliding-windows of 100kb (C). Windows belonging to the collinear genome are plotted in grey while other windows are coloured according to the inversion or the low-recombining region they belong to.

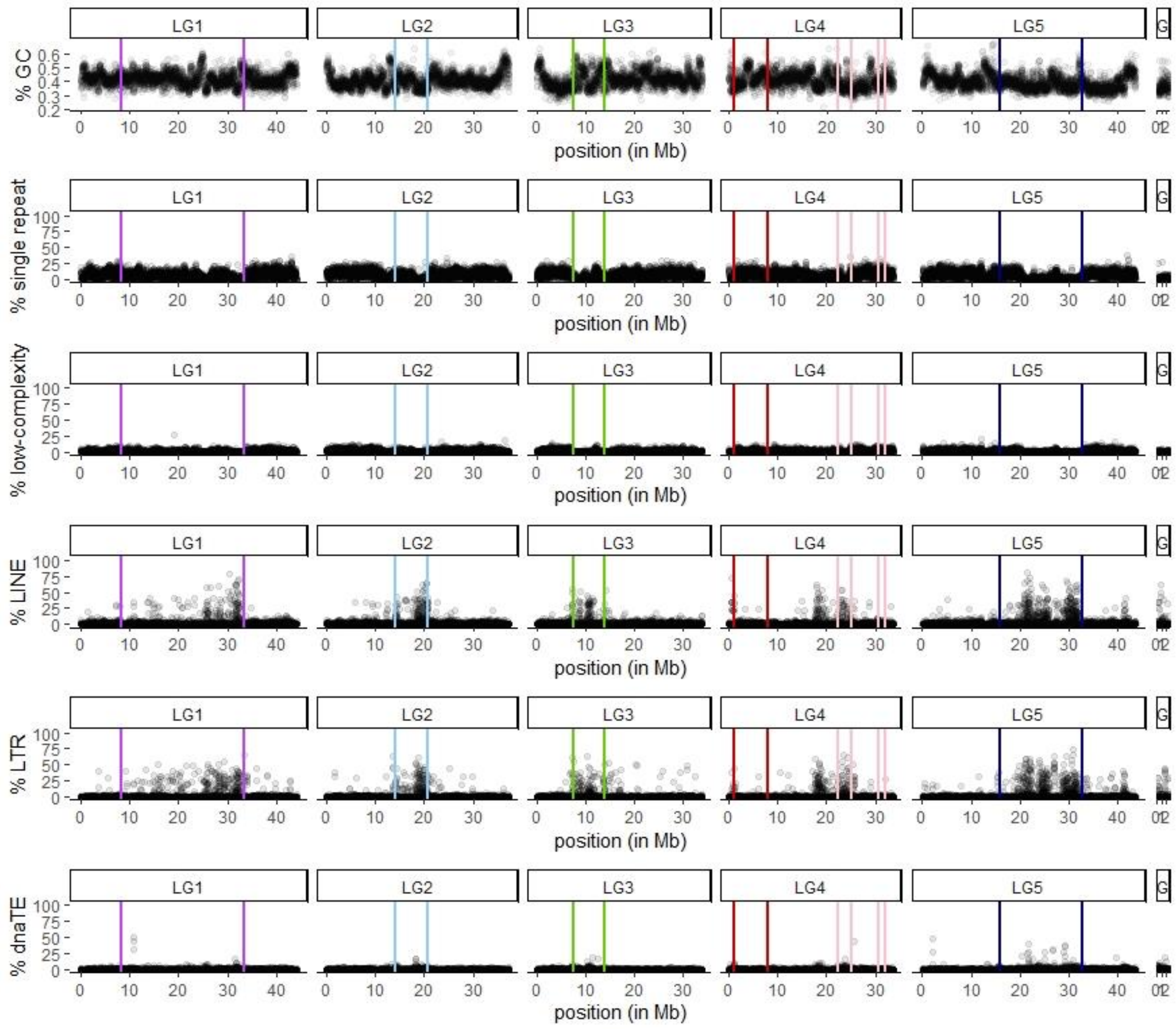

### Fig. S7: GC, repeat and TE content along the genome

Average percentage of GC or average percentage of bases belonging to repeated sequences or transposable elements, as annotated by RepeatMasker (Smit et al. 2015), by windows of 10 kb.

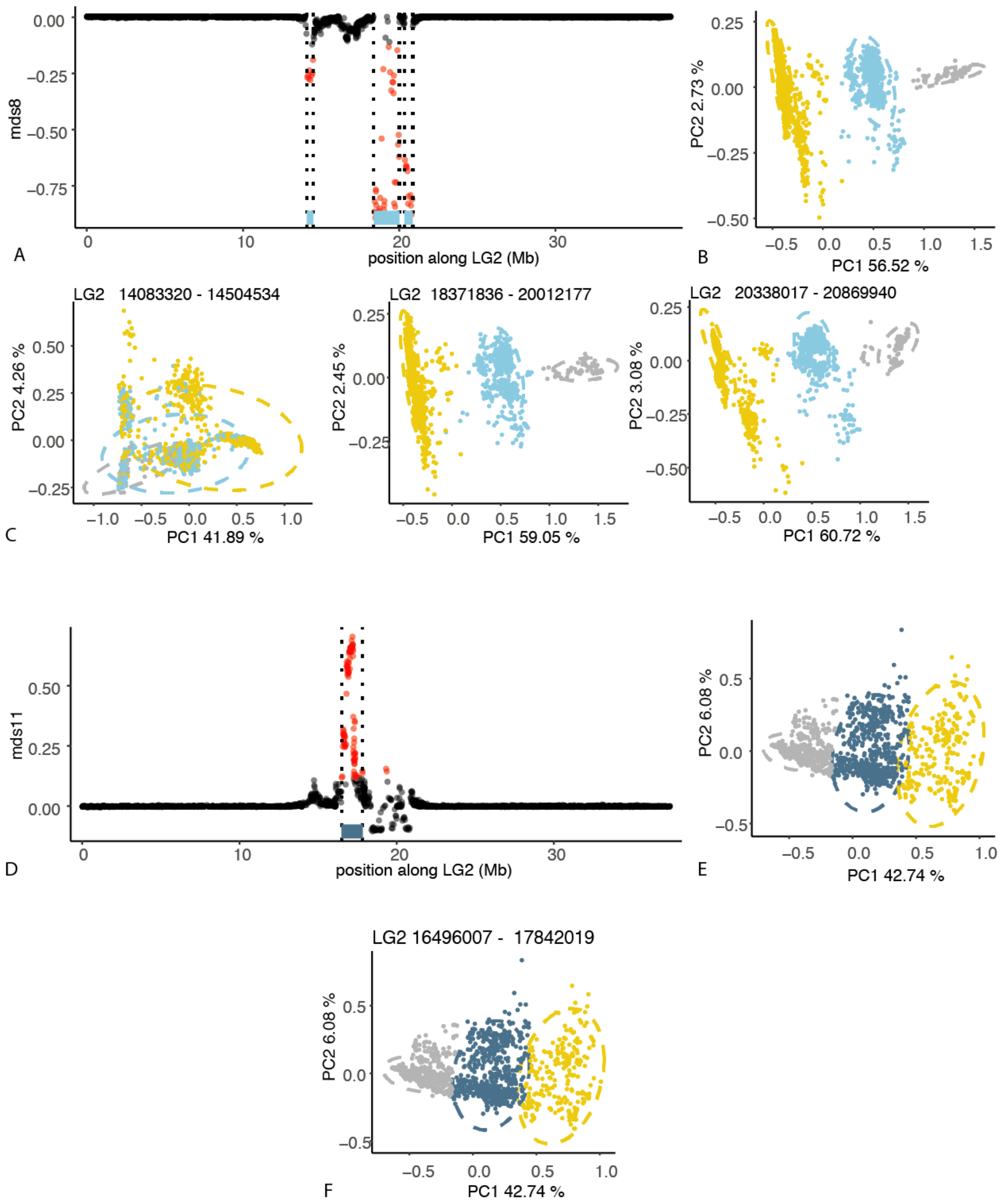

### Fig. S8: The low-recombining region on LG2 *Cf-Lrr(2)*

**(A-D)** MDS values of local PCAs along LG2 for the MDS axes on which they formed a cluster of outlier windows. **(B-E)** Principal component analysis (PCA) of genetic variation at SNPs within the cluster of outlier windows **(C-F)** Principal component analysis (PCA) of genetic variation at SNPs within each contiguous group of outlier windows, coloured by groups inferred from the PCA on the whole cluster.

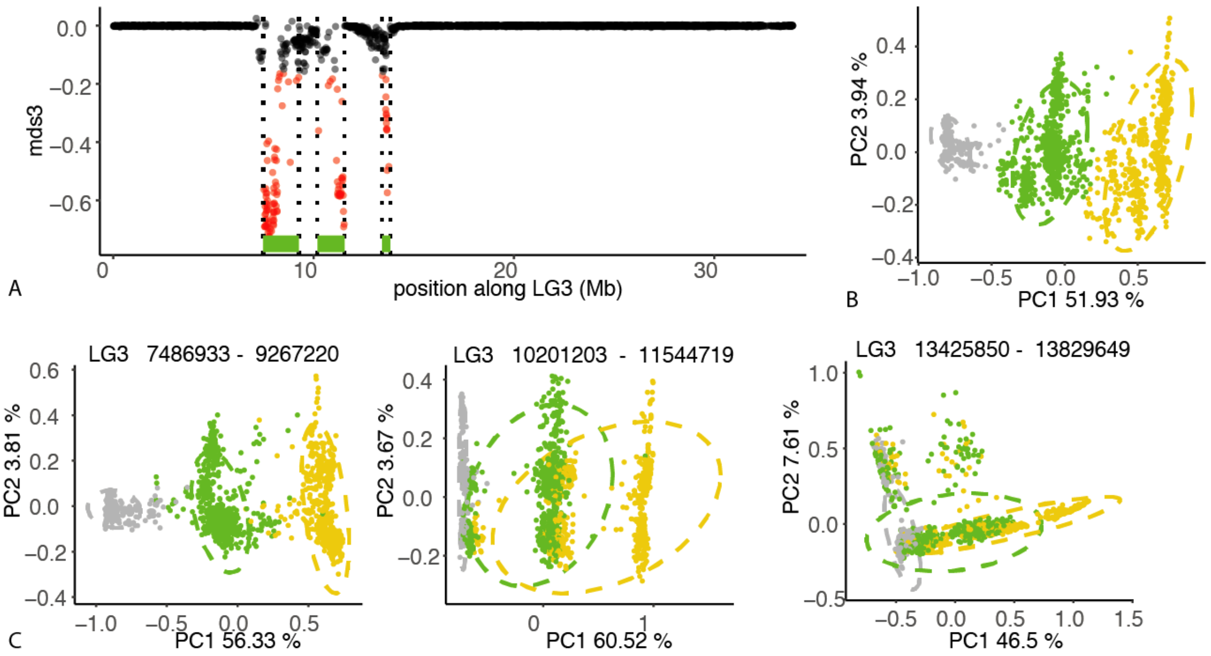

### Fig. S9: The low-recombining region on LG3 *Cf-Lrr(3)*

**(A)** MDS values of local PCAs along LG3 for the MDS axis on which they formed a cluster of outlier windows. **(B)** Principal component analysis (PCA) of genetic variation at SNPs within the cluster of outlier windows **(C)** Principal component analysis (PCA) of genetic variation at SNPs within each contiguous group of outlier windows, coloured by groups inferred from the PCA on the whole cluster.

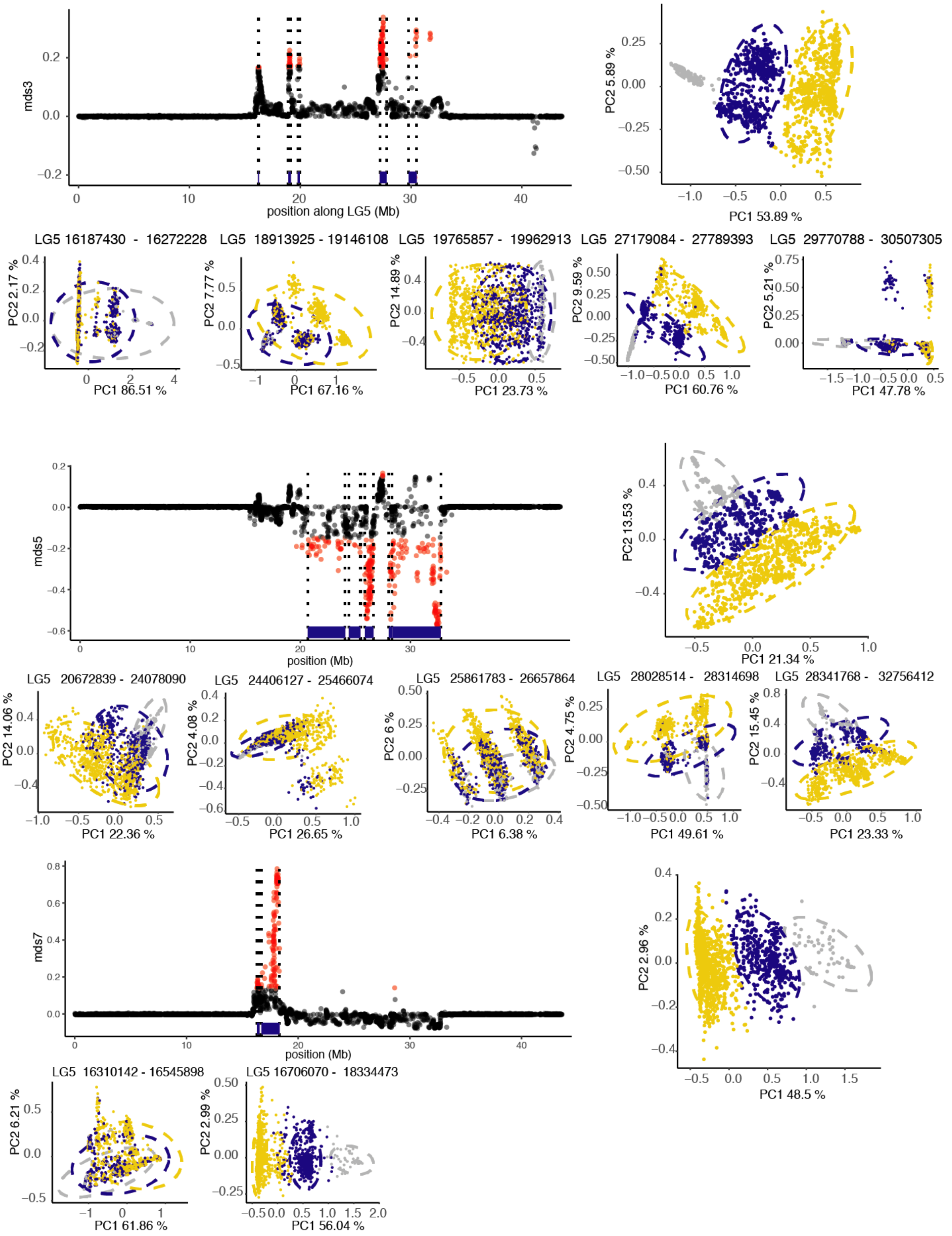

### Fig. S10: The low-recombining region on LG5 *Cf-Lrr(5)*

**(A-D-G)** MDS values of local PCAs along LG5 for the MDS axes on which they formed a cluster of outlier windows. **(B-E-H)** Principal component analysis (PCA) of genetic variation at SNPs within the cluster of outlier windows **(C-F-I)** Principal component analysis (PCA) of genetic variation at SNPs within each contiguous group of outlier windows, coloured by groups inferred from the PCA on the whole cluster. Clusters of windows from LG5 which were outliers along other MDS axis are not shown because they overlap with the same region and display similar patterns as mds3 or mds5.

| | BS | RB | BT | SI | KA | ME | GM | RC | AG | CE | SS | NB | CB | BP | HA | MA |
|---|---|---|---|---|---|---|---|---|---|---|---|---|---|---|---|---|
| BS | | 0.003 | 0.004 | 0.006 | 0.006 | 0.006 | 0.006 | 0.006 | 0.005 | 0.006 | 0.006 | 0.007 | 0.007 | 0.011 | 0.012 | 0.014 |
| RB | 0.003 | | 0.003 | 0.007 | 0.007 | 0.007 | 0.007 | 0.007 | 0.006 | 0.007 | 0.007 | 0.01 | 0.01 | 0.012 | 0.013 | 0.015 |
| BT | 0.006 | 0.004 | | 0.005 | 0.005 | 0.006 | 0.005 | 0.005 | 0.005 | 0.005 | 0.006 | 0.007 | 0.008 | 0.01 | 0.01 | 0.013 |
| SI | 0.005 | 0.007 | 0.011 | | 0.003 | 0.003 | 0.003 | 0.003 | 0.003 | 0.003 | 0.003 | 0.004 | 0.004 | 0.005 | 0.006 | 0.008 |
| KA | 0.009 | 0.008 | 0.005 | 0.01 | | 0.003 | 0.004 | 0.003 | 0.003 | 0.003 | 0.004 | 0.004 | 0.005 | 0.005 | 0.006 | 0.008 |
| ME | 0.005 | 0.007 | 0.011 | 0.002 | 0.009 | | 0.003 | 0.003 | 0.003 | 0.003 | 0.003 | 0.004 | 0.004 | 0.005 | 0.006 | 0.008 |
| GM | 0.01 | 0.013 | 0.021 | 0.004 | 0.021 | 0.004 | | 0.003 | 0.003 | 0.003 | 0.003 | 0.004 | 0.005 | 0.006 | 0.007 | 0.009 |
| RC | 0.007 | 0.008 | 0.014 | 0.002 | 0.013 | 0.002 | 0.003 | | 0.003 | 0.002 | 0.003 | 0.004 | 0.005 | 0.006 | 0.006 | 0.008 |
| AG | 0.006 | 0.006 | 0.005 | 0.004 | 0.004 | 0.004 | 0.011 | 0.006 | | 0.003 | 0.003 | 0.004 | 0.005 | 0.006 | 0.006 | 0.009 |
| CE | 0.007 | 0.008 | 0.012 | 0.003 | 0.011 | 0.003 | 0.004 | 0.002 | 0.005 | | 0.003 | 0.004 | 0.005 | 0.006 | 0.006 | 0.008 |
| SS | 0.009 | 0.01 | 0.015 | 0.004 | 0.013 | 0.004 | 0.004 | 0.003 | 0.006 | 0.002 | | 0.004 | 0.005 | 0.006 | 0.006 | 0.008 |
| NB | 0.007 | 0.009 | 0.007 | 0.005 | 0.004 | 0.005 | 0.013 | 0.007 | 0.004 | 0.006 | 0.008 | | 0.003 | 0.005 | 0.006 | 0.009 |
| CB | 0.007 | 0.009 | 0.012 | 0.004 | 0.01 | 0.004 | 0.008 | 0.005 | 0.006 | 0.005 | 0.007 | 0.004 | | 0.006 | 0.007 | 0.01 |
| BP | 0.011 | 0.012 | 0.01 | 0.007 | 0.006 | 0.007 | 0.014 | 0.009 | 0.006 | 0.008 | 0.008 | 0.005 | 0.008 | | 0.003 | 0.005 |
| HA | 0.015 | 0.014 | 0.01 | 0.011 | 0.006 | 0.011 | 0.021 | 0.014 | 0.007 | 0.012 | 0.012 | 0.006 | 0.012 | 0.003 | | 0.004 |
| MA | 0.019 | 0.019 | 0.02 | 0.011 | 0.016 | 0.011 | 0.013 | 0.01 | 0.012 | 0.009 | 0.008 | 0.012 | 0.014 | 0.007 | 0.008 | |

### Fig. S11: Pairwise $F_{ST}$ between all population pairs

Pairwise $F_{ST}$ between all population pairs, ordered by proximity from North to South and coloured by geographic regions. The values above the main diagonal shows $F_{ST}$ based on LD-pruned SNPs and those below the main diagonal show $F_{ST}$ based on all SNPs.

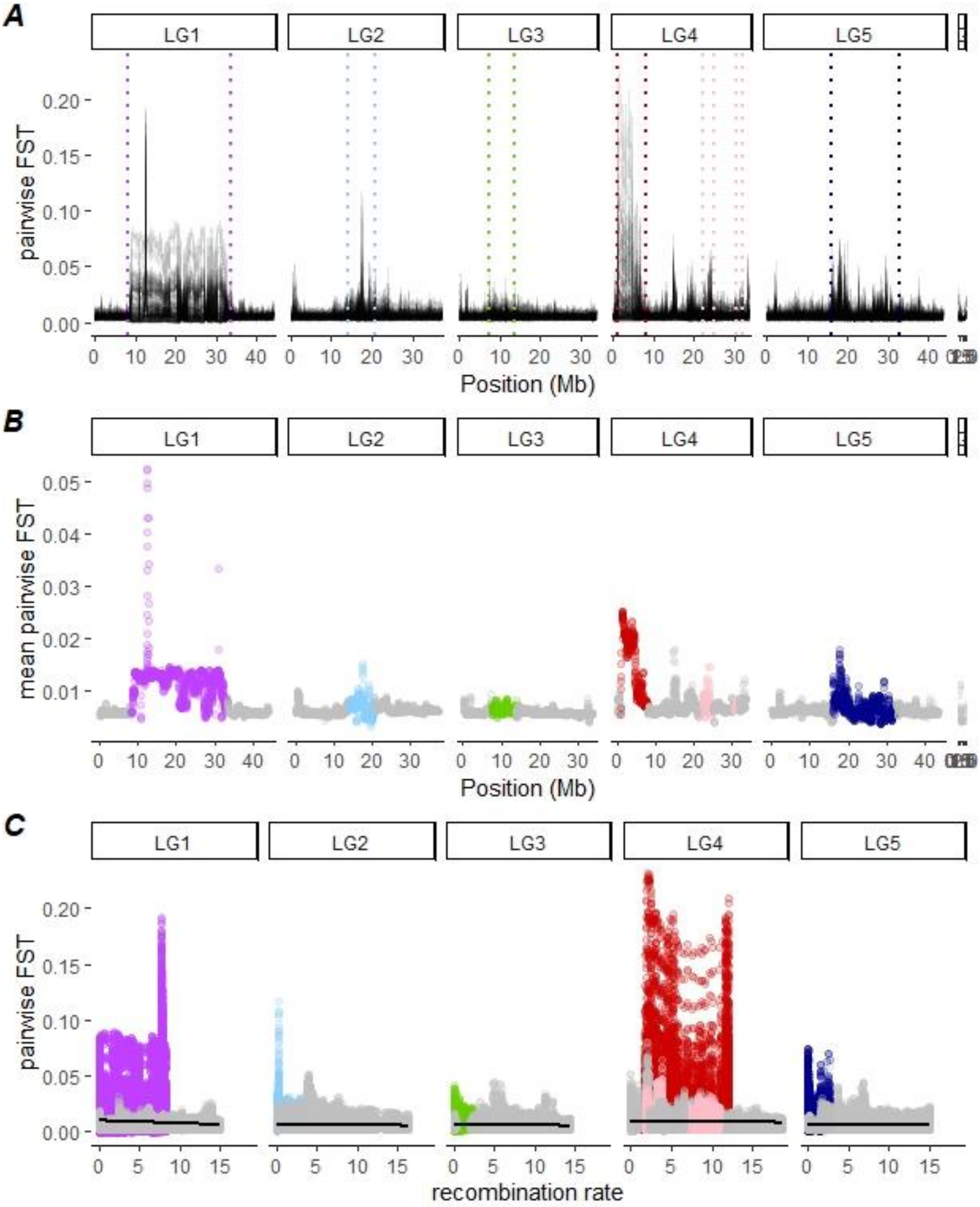

Fig. S12: Pairwise $F_{ST}$ between geographic populations along the genome and as a function of recombination

(A) Pairwise $F_{ST}$ along the genome for each pair of populations. Each line is a pair of populations (plotted with transparency) hence darker colours appear when the values for the different pairs of populations overlap. (B) Pairwise $F_{ST}$ averaged across pairs of populations along the genome and coloured for the inversions and low-recombining regions. (C) Pairwise $F_{ST}$ for each pair of populations as a function of recombination rate. All points are sliding-windows of 100kb.

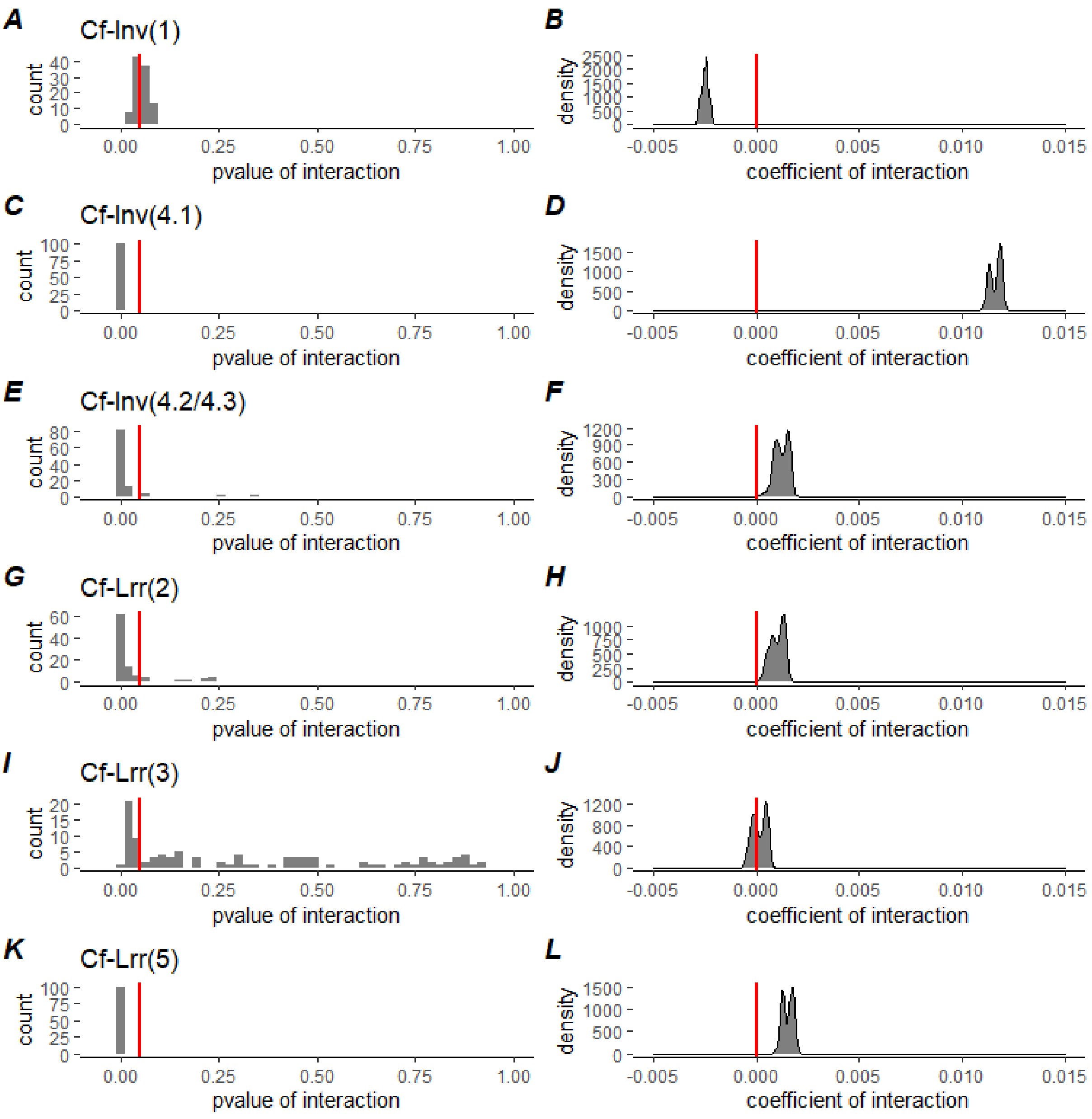

### Fig. S13: Isolation-by-Resistance full models comparing each region of interest to collinear regions

**(A-C-E-G-I-K)** Distribution of the p-value of the interaction terms in 100 models explaining genetic distance by physical distance with the type of genomic region (given inversion or collinear region) as co-variable. The red line indicates p=0.05. **(B-D-F-J-L)** Distribution of the slope coefficient of the interaction terms in 100 models explaining genetic distance by physical distance with the type of genomic region (given inversion or collinear region) as co-variable. The red line indicates a null slope (no interaction) A positive slope indicates that the IBR signal is stronger in the inversion/low-recombining region than in the collinear genome. A null slope indicates a comparable signal while a negative slope indicates a weaker signal.

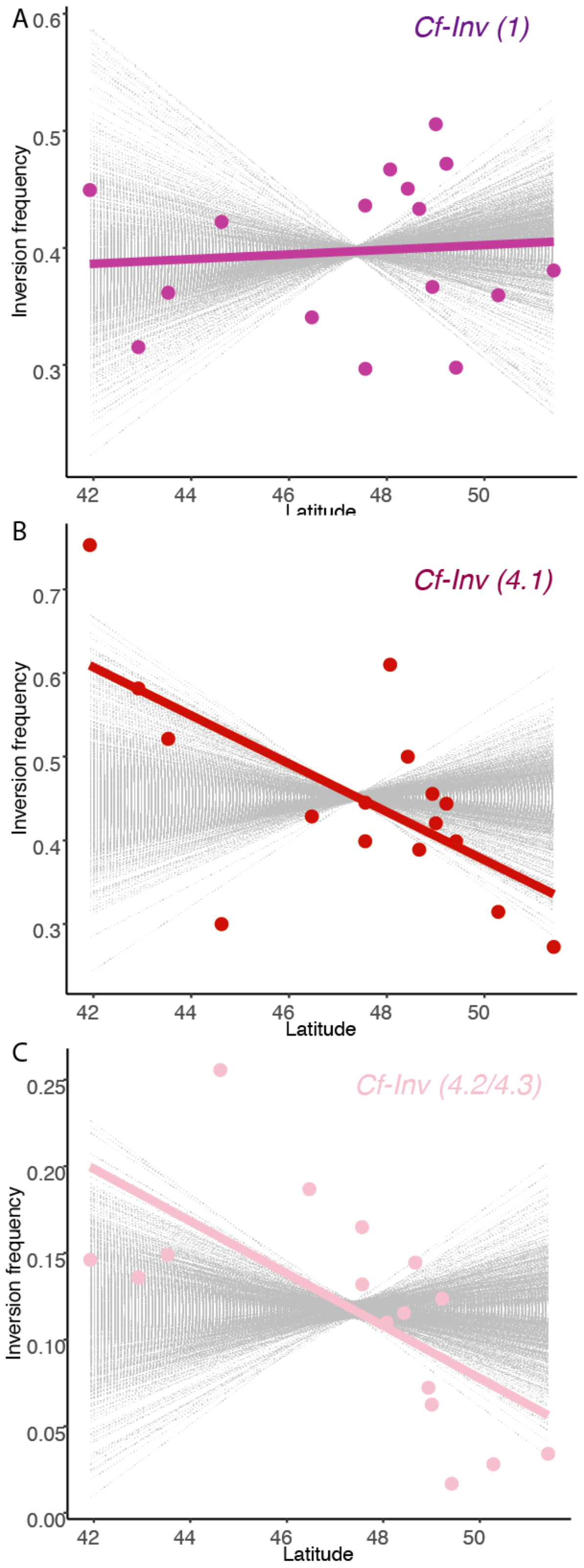

### Fig. S14: Latitudinal cline of frequencies for the major inversions

Coloured points depicted the frequencies of the rarest inversion arrangement for each location, plotted by latitude, the coloured line depicting a linear cline. Grey lines represent approximated latitudinal clines of frequencies for 1,000 random SNPs chosen to have the same average frequency as the inversion across the whole area.

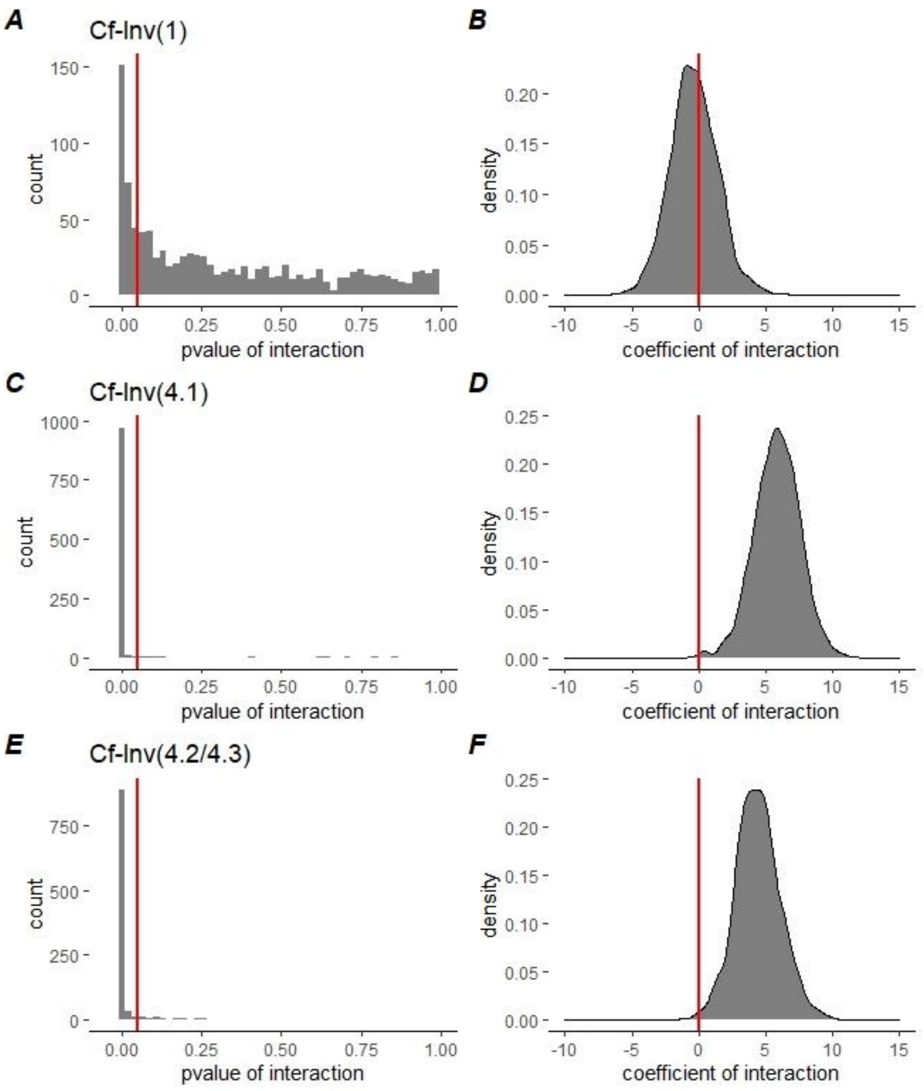

### Fig. S15: Clinal association full models comparing each inversion to collinear SNPs

**(A-C-E)** Distribution of the p-value of the interaction terms in 1000 models explaining a variant frequency by latitude with the type of variant (given inversion or randomly-picked SNP with same average frequency) as co-variable. The red line indicates p=0.05. **(B-D-F)** Distribution of the slope coefficient of the interaction terms in those 1000 models. The red line indicates a null slope (no interaction). A positive slope indicates that the cline is stronger for the inversion frequency than for SNPs. A null slope indicates a comparable signal.

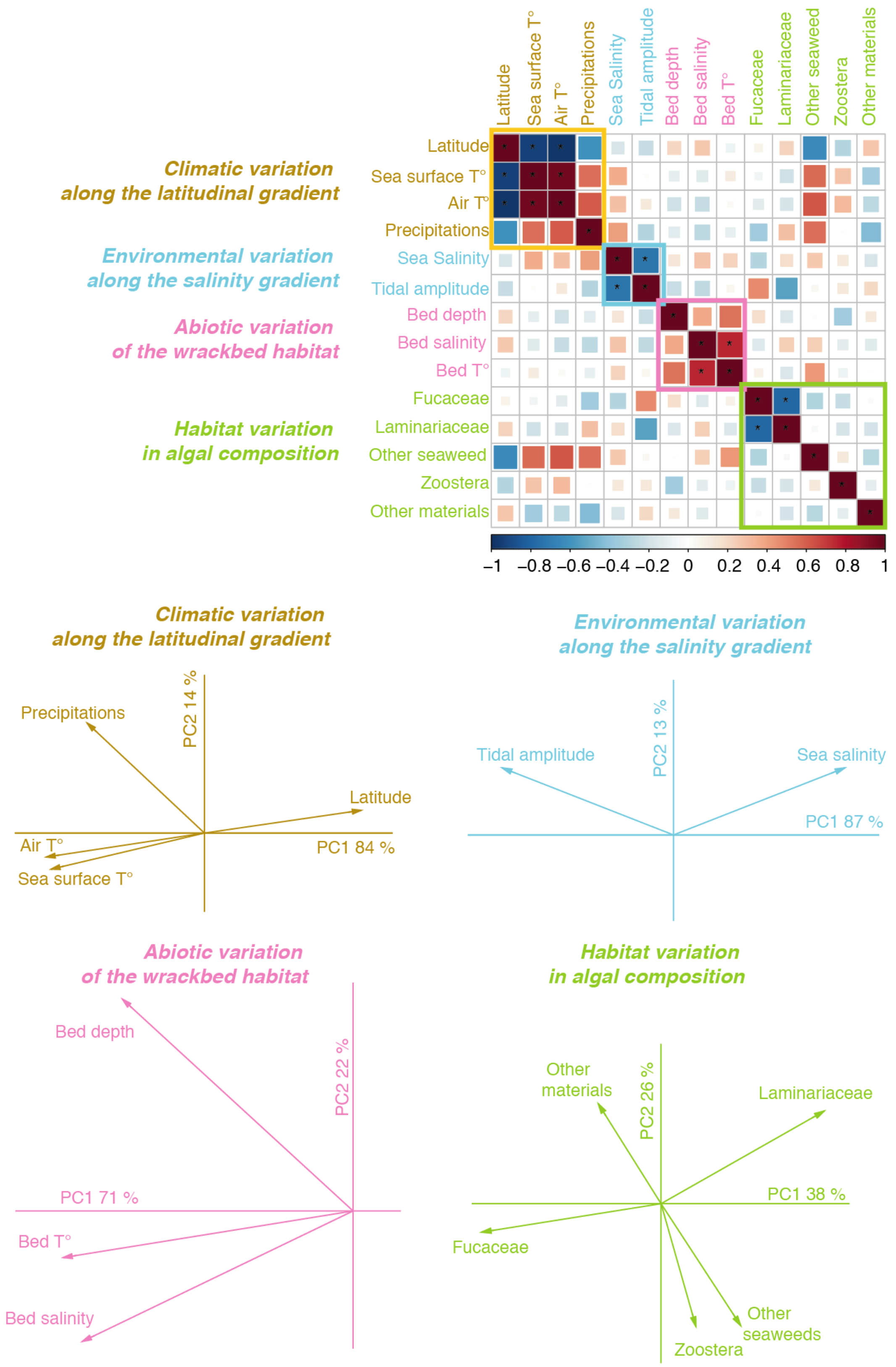

Fig S16: Correlations between environmental variables and summary variables by PCA

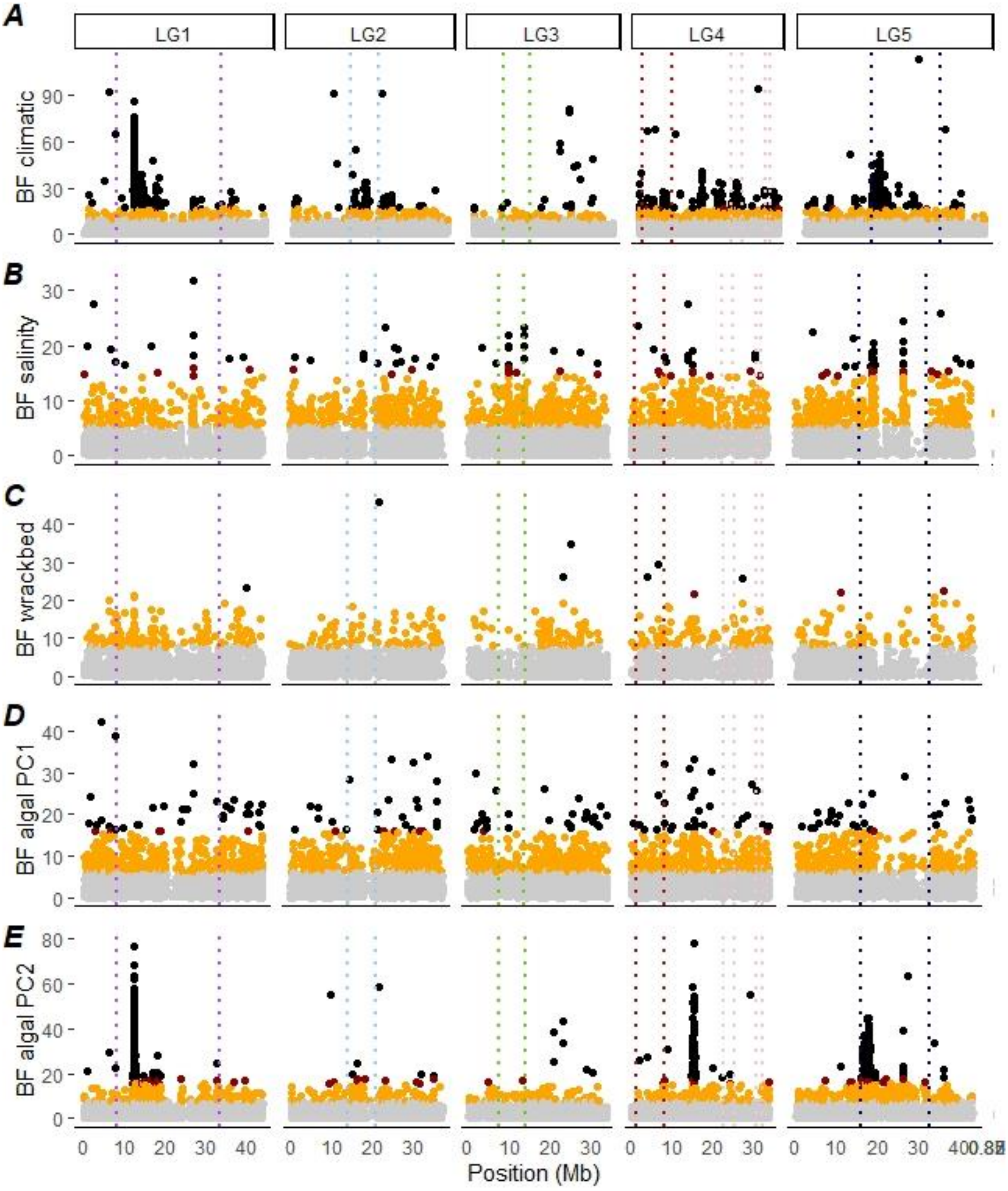

### Fig S17: Environmental associations with Baypass (uncontrolled for population structure)

The Manhattan plot shows the Bayesian factor from the environmental association analysis performed in Baypass, without controlling for population structure. Points are coloured according to false-discovery rate (black: <0.00001, red: <0.0001, orange: <0.001)

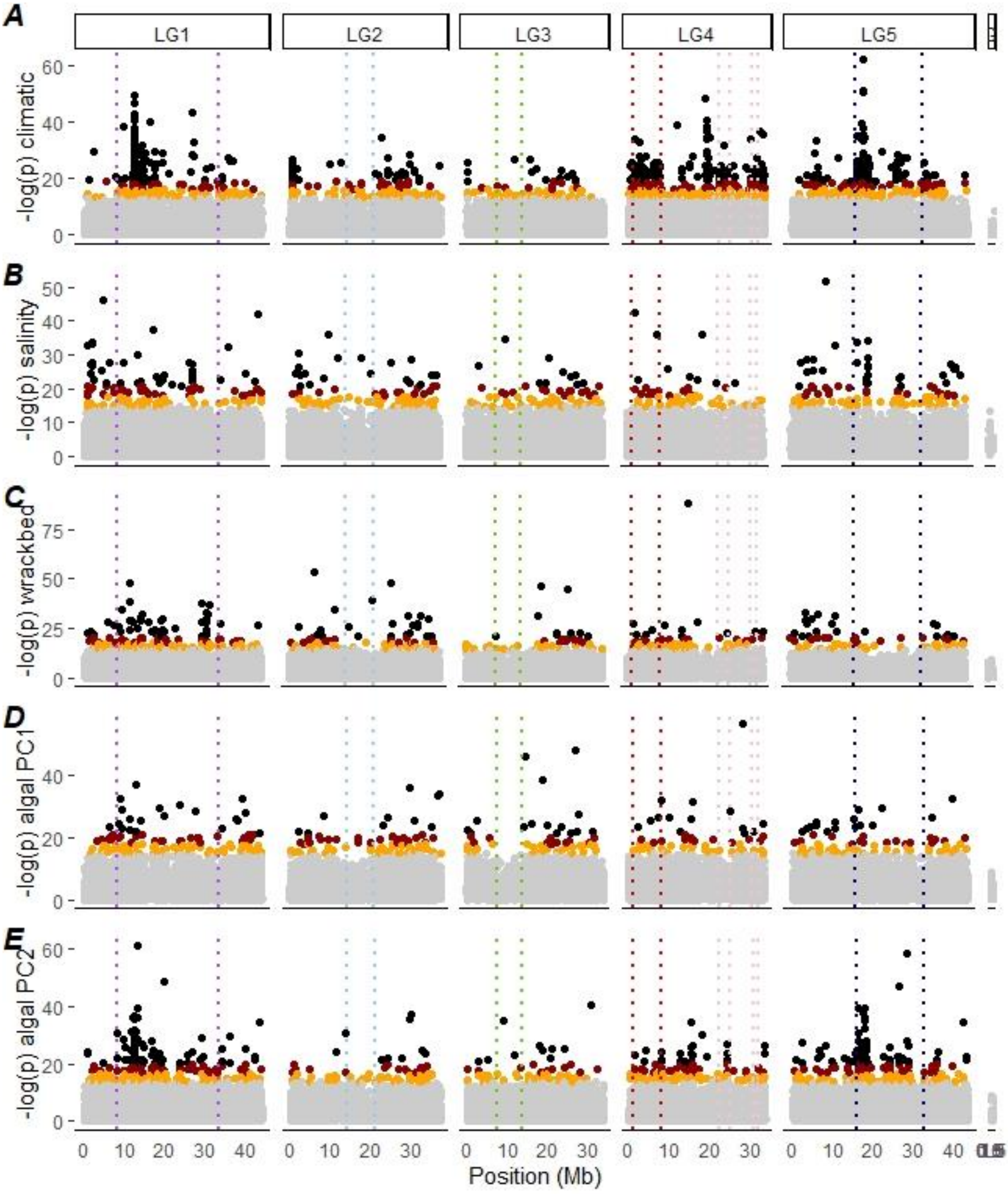

### Fig S18: Environmental associations with LFMM (K=4)

The Manhattan plot shows the log of the p-value from the environmental association analysis performed in LFMM, controlling for K=4 latent factors. Points are coloured according to false-discovery rate (black: <0.00001, red: <0.0001, orange: <0.001)

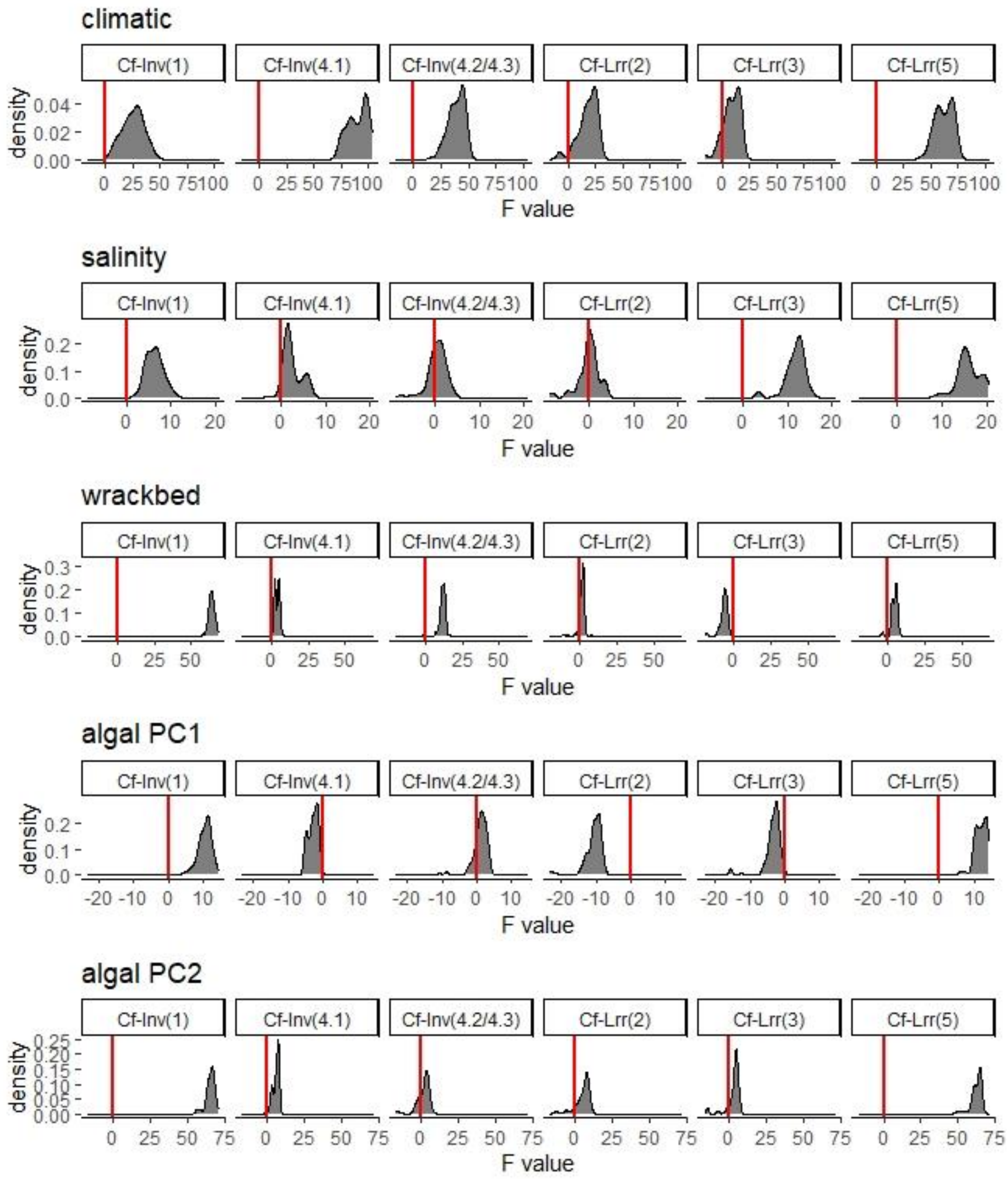

Fig. S19: Models comparing the distribution of association scores between each inversion and collinear blocks

Distribution of the F-value in 100 models comparing the distribution of association scores between SNPs and environmental predictors within inversion to within collinear blocks. The red line indicates 0, the distributions are not different while a positive value indicates that the environmental association is stronger in the inversion/low-recombining region than in the collinear genome.

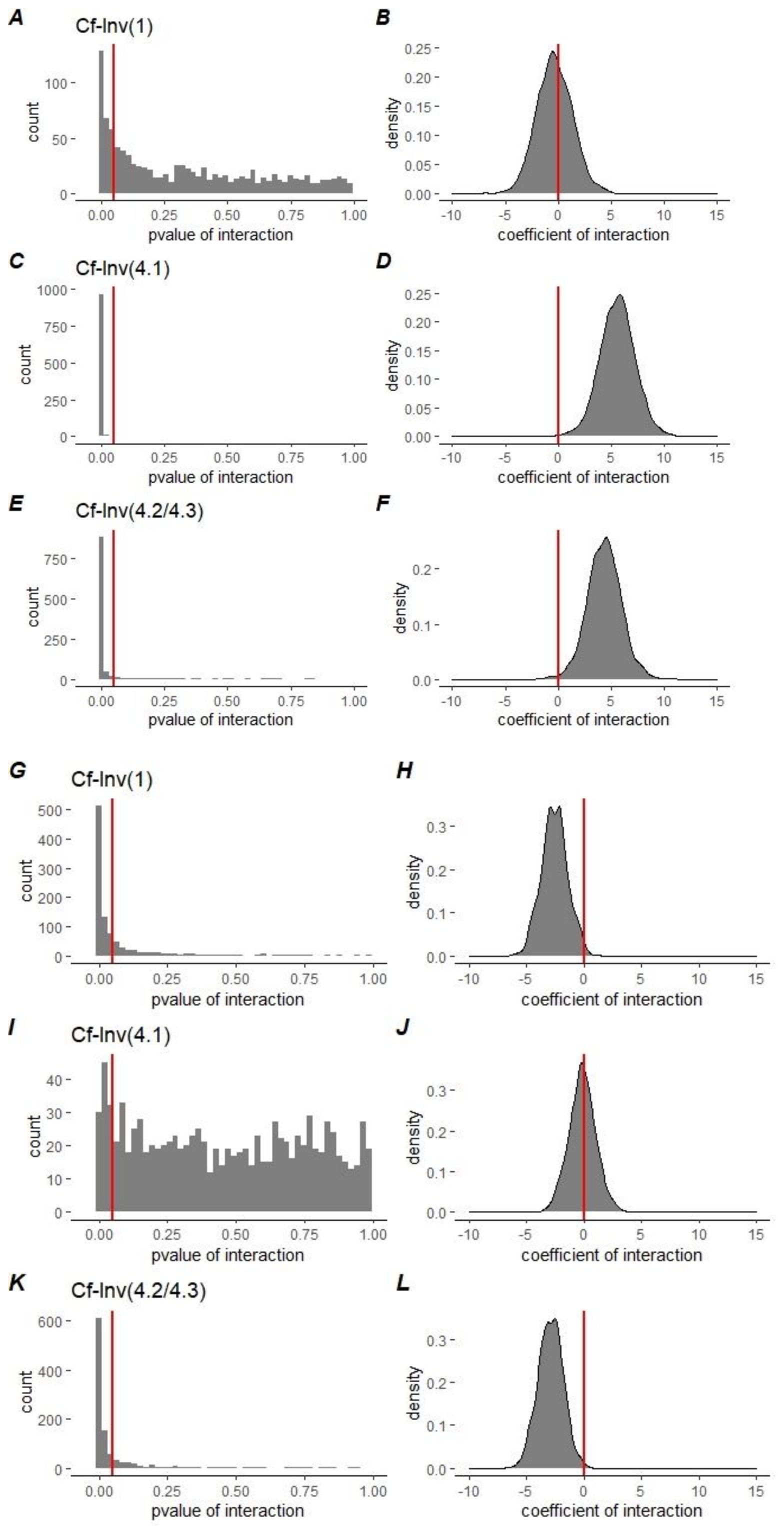

### Fig. S20: Environmental association full models comparing each inversion to collinear SNPs

Distribution of the p-value, and the slope coefficient of the interaction terms in 1000 models explaining a variant frequency by climatic variation **(A-F)** or wrackbed characteristics **(G-L)**, with the type of variant (given inversion or randomly-picked SNP with same average frequency) as co-variable. On the left (p-values, the red line indicates p=0.05). On the right, the red line indicates a null slope (no interaction). A positive/negative slope indicates that the association is stronger for the inversion frequency than for SNPs. A null slope indicates a comparable signal.

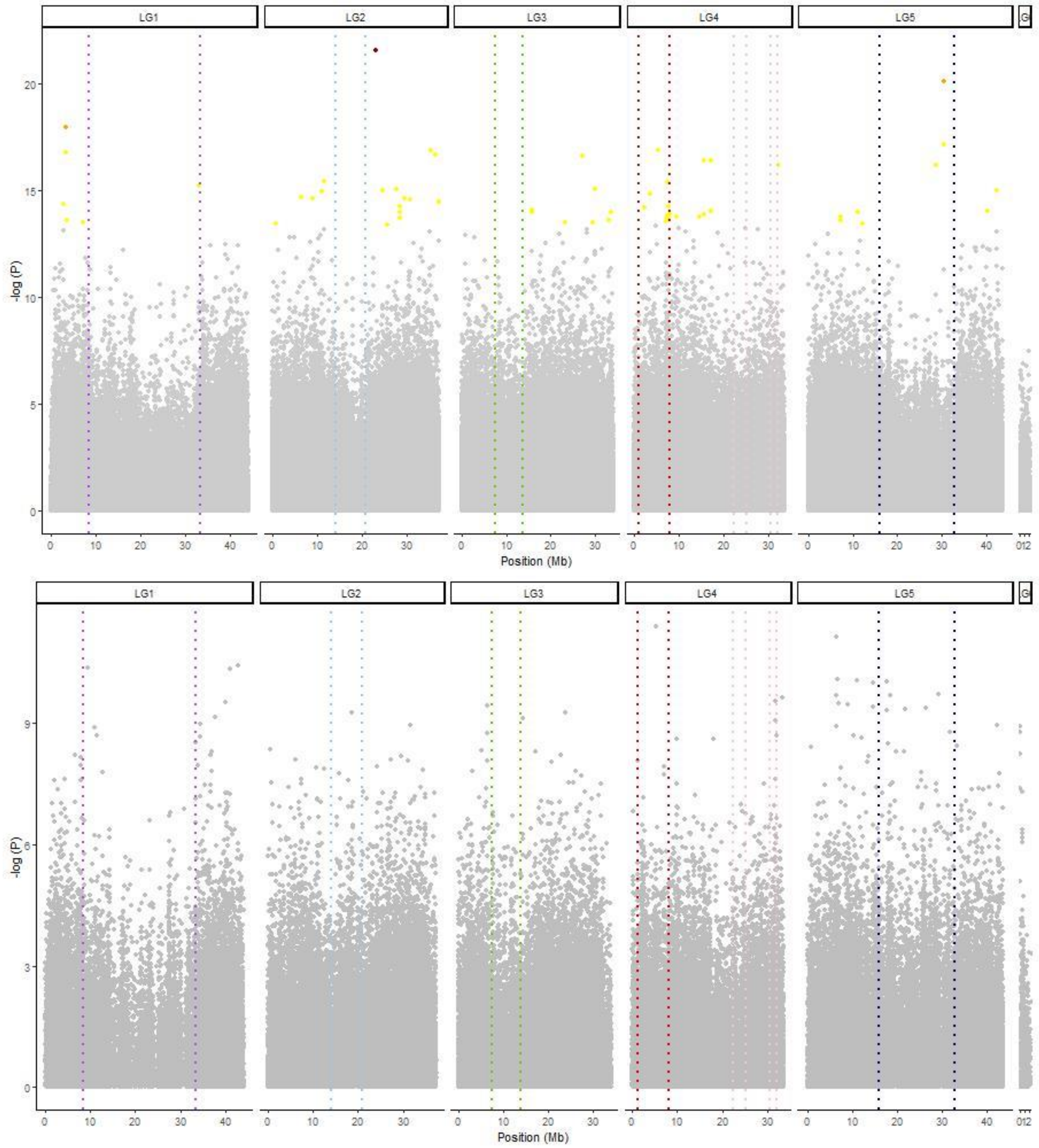

### Fig S21: GWAS on wing size within each homokaryotypes group

Above: GWAS for ββ (N= 436 sized individuals). Below: GWAS for αα (N=140 sized individuals). SNPs coloured show significant association with size with a FDR of 0.05 (yellow), 0.01 (orange), 0.001 (red). No SNP significantly associated with size was found on the αα subset.

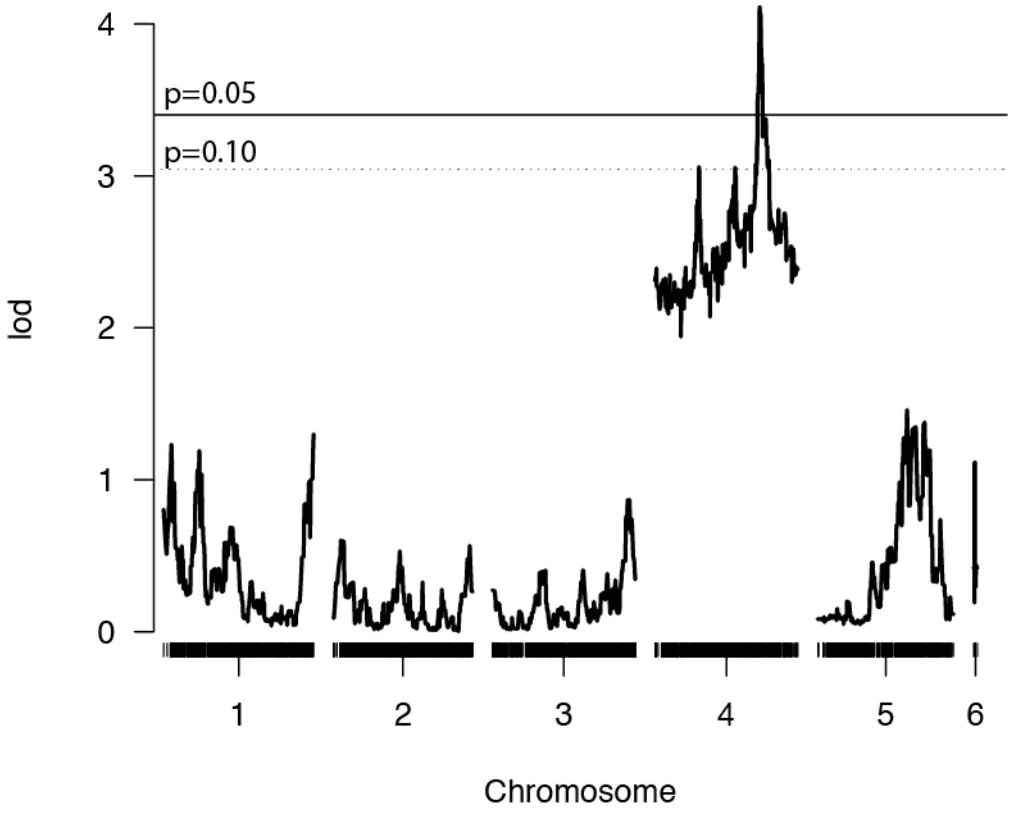

### Fig S22: QTL for chill-coma recovery

The scale on the y-axis depicts the LOD score (Logarithm of odds) for an association between each genetic marker and recovery after a cold-induced coma. The horizontal line represents the LOD genome-wide significance threshold determined through permutations with alpha = 0.05.
